# Identity and functions of monoaminergic neurons in the predatory nematode *Pristionchus pacificus* reveal nervous system conservation and divergence

**DOI:** 10.1101/2025.10.16.682888

**Authors:** Curtis M. Loer, Hyunsoo Yim, Luke T. Geiger, Yasmin H. Ramadan, Megan F. Hampton, Diana V. Bernal, Heather R. Carstensen, Jorge Morgan, Laura Rivard, Theresa Medina, Steven J. Cook, Misako Okumura, James Lightfoot, Oliver Hobert, Ray L. Hong

## Abstract

Changes in neurotransmitter usage in homologous neurons may drive evolutionary adaptations in neural circuits across animal phylogeny. The predatory nematode *Pristionchus pacificus* can be used as a model system to examine nervous system evolution by comparing neurotransmitter expression with that of *C. elegans* and other nematodes. Here we characterize *P. pacificus* neurotransmitter expression and function in specific neurons, focusing on its complete set of monoaminergic neurons. We discover patterns of conservation as well as novelties. We examine the roles of monoamines in specific behaviors using neurotransmitter synthesis and vesicular transporter mutants, finding possible differences in the control of host-finding and dispersal behavior.

## Introduction

The model organism nematode *Caenorhabditis elegans* was selected in part to study mechanisms of nervous system function and development (Brenner, 1974). As a base for such studies, the complete anatomy, invariant cell lineage, and neuronal connectivity (aka the ‘connectome’) were determined early in its emergence as new ‘model organism’ (Sulston and Horvitz, 1977) (Sulston et al., 1983) (White et al., 1986) Studying this nematode has revealed many fundamental insights into nervous system biology (Horvitz and Sulston, 1990) (Driscoll 1996) (Rand et al., 2000) (Hobert 2013) (Shaham, 2015) (Hobert, 2016) (Markaki and Tavernarakis, 2020). More recently, the ‘base’ of nematode knowledge has been expanded and strengthened by re-analysis of the *C. elegans* connectome, with the addition of multiple life stages and the adult male connectome (Cook et al., 2019) (Witvliet et al., 2021) (Jarrell et al., 2012) (Yim et al., 2023), near complete mapping of all classical neurotransmitters used by individual neurons in both sexes (Pereira et al., 2015) (Gendrel et al., 2016) (Serrano-Saiz et al., 2017) (Wang et al., 2024), and extensive single cell transcriptomes in identified neurons (Taylor et al., 2021) (Wolfe et al., 2025) (Morillo et al., 2025). In addition to studies on *C. elegans,* a second model nematode was established to facilitate study of comparative and evolutionary mechanisms, the free-living diplogastrid nematode *Pristionchus pacificus* (Sommer et al., 1995) (Felix et al., 1999) (Sternberg 2015).

*P. pacificus* is now a powerful nematode model in its own right, with advanced genetic tools and base of organismal knowledge, including a complete head connectome and embryonic cell lineage, which demonstrates a nearly 1-1 cell homology between *P. pacificus* and *C. elegans*, especially in the nervous system (Vangestel et al., 2008) (Hong et al., 2019) (Cook et al., 2025). Here we examine the monoaminergic neurons in *P. pacificus,* behaviors requiring monoamine function, and differences between *P. pacificus* and *C. elegans.* Comparisons between these distantly-related species with remarkably similar nervous systems may provide valuable insights into how neurotransmitter usage evolves among homologous neurons in species that have similar anatomical and behavioral constraints.

One striking difference between *P. pacificus* and *C. elegans* lies in the diversity of their feeding behaviors. While both species eat bacteria, *P. pacificus* is an omnivorous and predatory nematode species that can kill and feed upon larvae of other nematode species, and cannibalizes *P. pacificus* conspecifics (Wilecki et al., 2015) (Lightfoot et al., 2021;Quach and Chalasani 2022). *P. pacificus* can develop ‘teeth’ used to puncture and then feed on other nematodes. How *P. pacificus* consumes bacteria is also different: these worms lack the pharyngeal grinder used by *C. elegans* to macerate bacteria. These dramatic differences in feeding behavior and apparatus are accompanied by differences in connectivity among enteric and central neurons of *P. pacificus* and *C. elegans* (Bumbarger et al., 2013) (Sieriebriennikov et al., 2018) (Cook et al., 2025). Interestingly, both bacterial and predatory feeding behaviors in these species are modulated by serotonin signaling in *P. pacificus* (Okumura et al., 2017) (Ishita et al., 2021) but precisely which neurons are responsible, both releasing and responding to serotonin, has not yet been determined.

Biogenic amines (or monoamines) such as serotonin (5HT), dopamine (DA), tyramine (TA), octopamine (OA), epinephrine, norepinephrine, and histamine serve as neurotransmitters, neuromodulators, and endocrine factors broadly conserved among Bilaterians, controlling and modulating a wide array of behaviors [Figure 1; (Bauknecht and Jékely 2017) (Edsinger and Dölen 2018) (Moroz et al., 2021)]. For example, in higher vertebrates, serotonin plays a role in mood and satiety, while dopamine is associated with motivation. Some similar roles of monoamines such as serotonin have been demonstrated in invertebrates, including *C. elegans* (Chase and Koelle 2007). Interestingly, in the vertebrates, the autonomic neurotransmitters (or hormones) norepinephrine and epinephrine – both dopamine derivatives – seem to function in similar responses to those found in invertebrates in which the structurally similar TA and OA – both tyrosine derivatives – mediate that signaling (Roeder 2005) (Bauknecht and Jékely 2017). The vertebrate adrenergic signaling molecules norepinephrine and epinephrine are typically absent from invertebrate species.

**Figure 1.**
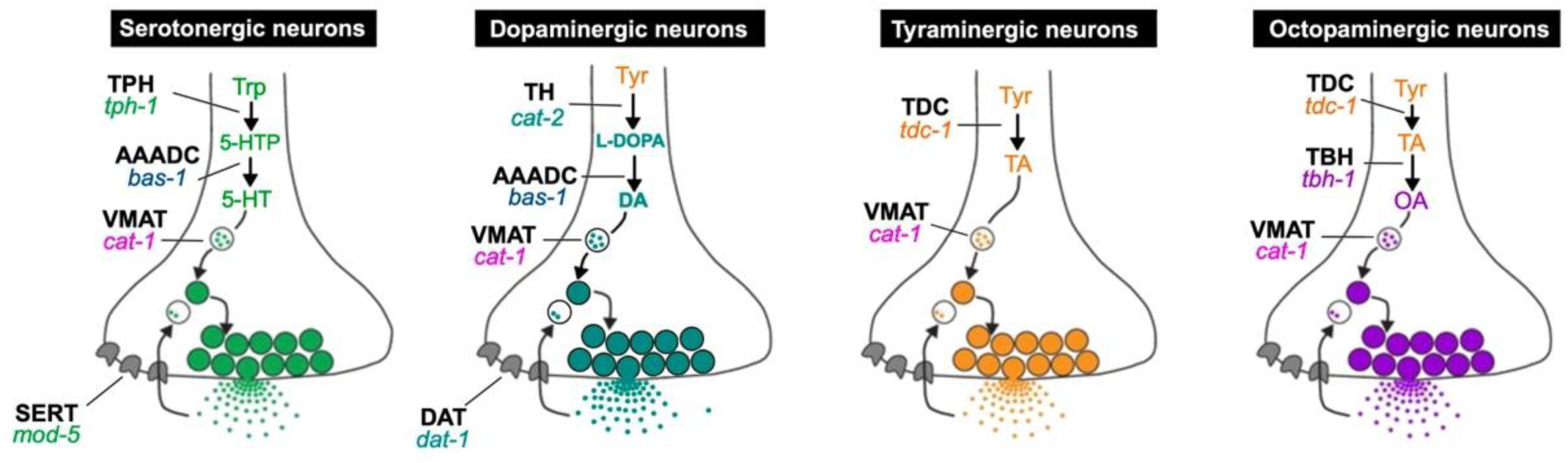
Monoamine synthesis and transport pathways. Proteins encoded by identified nematode genes are indicated in black, bold, capital letters (abbreviations: TPH – tryptophan hydroxylase, AAADC – aromatic amino acid decarboxylase, VMAT – vesicular monoamine transporter, SERT – serotonin transporter, TH – tyrosine hydroxylase, DAT – dopamine transporter, TDC – tyrosine decarboxylase, TBH – tyramine beta-hydroxylase); nematode gene names (originally from *C. elegans*) are italicized below the protein name abbreviations. Function of the *cat-1* gene, encoding VMAT, is required in all monoaminergic neurons for packaging monoamines into synaptic vesicles to allow subsequent synaptic release. Other genes pictured are required for monoamine biosynthesis or transport across the plasma membrane (‘reuptake’) to stop the action of the neurotransmitter following vesicular release at synapses (or extrasynaptically).

In the nematode *C. elegans*, serotonin, dopamine, tyramine and octopamine can be detected directly from worm extracts and via cytochemical techniques; gene expression patterns of rate-limiting biosynthesis enzymes and transporters specific to bioamines can also be detected in specific identified neurons [summarized in (Loer and Rand 2022)]. More recently, the application of single cell transcriptomics and CRISPR-based native locus reporters have expanded the list of likely monoaminergic neurons in *C. elegans* (Wang et al., 2024). A variety of neuronal classes in the *C. elegans* hermaphrodite – sensory, motor and interneurons (ADFs, AIMs, HSNs, NSMs, RIH, VC4-5) – use serotonin; several ciliated mechanosensory neurons use dopamine (ADEs, CEPs, PDEs); one pair of motor neurons uses tyramine (RIMs) and one pair of interneurons uses octopamine (RICs). Additional sex-specific neurons in the male also use these neurotransmitters. Together, these aminergic neurons in the *C. elegans* adult hermaphrodite constitute a small minority - 23 out of 302 neurons – but are involved in behaviors ranging from food satiety signaling, regulation of locomotory rates, foraging, egg-laying [(Trent et al., 1983) (Desai et al., 1988) (Lints and Emmons 1999) (Sawin et al., 2000) (Sze et al., 2000); review by (Chase and Koelle 2007)]. There are also a few neuron-like or neuroendocrine cells that likely release monoamine neurotransmitters, such as the two pairs of uterine ventral (uv1) cells in the hermaphrodite, which synthesize tyramine (Alkema et al., 2005), and male-specific glia (spicule socket cells) that synthesize dopamine (LeBoeuf et al., 2014).

In *P. pacificus*, some monoaminergic neurons have been identified previously via antibody staining [serotonin-immunoreactive (IR) cells], including the highly conserved pharyngeal NSM neurons (Loer and Rivard, 2007) (Rivard et al., 2010) (Wilecki et al., 2015). The pharyngeal I1 neurons, which are not serotonin-IR in *C. elegans,* are moderately serotonin-IR in *P. pacificus* (Wilecki et al., 2015). In the hermaphrodite ventral nerve cord, motor neurons VC1-4, positioned on either side of the vulva (egg-laying pore), are strongly serotonin-IR and appear to innervate vulval muscles (Loer and Rivard, 2007). These neurons are lineally homologous to VC3-6 in *C. elegans;* although VC4-5 are weakly and unreliably serotonin-IR in *C. elegans,* they have been shown to function in egg laying behavior via serotonin release (Duerr et al., 1999). Interestingly, the HSNs (Hermaphrodite-specific neurons), which innervate vulval muscles, and are essential for normal egg laying in *C. elegans* (Schafer, 2005), are either absent or at least do not express serotonin in *P. pacificus* (Loer and Rivard, 2007). Other serotonin-IR neurons have been observed in *P. pacificus,* but not reliably identified. Dopamine neurons in *P. pacificus* have been identified by formaldehyde-induced fluorescence (FIF) and 5HTP-induced serotonin-IR (explained elsewhere), showing the same neurons as found in *C. elegans:* head CEPs and ADEs, and mid-posterior body wall PDEs (Rivard et al., 2010). Likely tyraminergic and octopaminergic neurons have recently been identified in *P. pacificus* via reporter fusions with their synthetic enzyme genes (Eren et al., 2024).

To map all monoaminergic neurons in *P. pacificus*, we examined expression of the sole vesicular monoamine transporter (VMAT) gene *cat-1*, which shares 1-1 orthology with *C. elegans cat-1*/VMAT (Duerr et al., 1999). VMAT can transport all known monoamine neurotransmitters into synaptic vesicles (Yaffe et al., 2018). We examined *in situ* mRNA expression by hybridization chain reaction (HCR) and a *cat-1* reporter, and similarly expression of all monoamine biosynthetic enzyme genes, and their co-expression with *cat-1.* We further re-examined the expression of a previously described *tph-1* reporter and of serotonin immunoreactivity, identifying and corroborating differences in neurons expressing serotonin in *P. pacificus* vs. *C. elegans*. To gain insights into the evolutionary origins of differences seen in serotonergic neurons, we examined the staining patterns of serotonin-IR neurons in many other nematode species, including re-examining patterns from previously published studies (Loer and Rivard, 2007) (Rivard et al., 2010). We find that the pattern of serotonin-IR neurons in *C. elegans* is derived, whereas that seen in *P. pacificus* is closer to a likely ancestral pattern.

To investigate behaviors that require monoaminergic neuron function in *P. pacificus*, we characterized loss-of-function mutants in the *cat-1* gene and some monoamine biosynthetic enzyme genes. In *P. pacificus cat-1* mutants, we observed defects in egg laying in adult hermaphrodites, suppression of head movements during backward locomotion, and in nictation, a nematode host-finding and dispersal-promoting ‘waving’ behavior seen in developmentally arrested dauer larvae, including *P. pacificus* dauers (Brown et al., 2011). In *C. elegans,* cholinergic signaling has been implicated in regulating nictation; serotonin and dopamine appear not to play a role (Lee et al., 2012). Whether tyramine or octopamine regulate nictation has not been tested. We observed that both serotonin and tyramine/octopamine regulate nictation in *P. pacificus*.

## Methods

### Nematode strain growth and maintenance

*P. pacificus* and other nematode strains were maintained at ∼21°C on NGM plates seeded with *E. coli* OP50 for food as described previously (Cinkornpumin et al., 2014); these methods are derived from standard *C. elegans* culture methods (Brenner, 1974). *P. pacificus* and other nematode strains used are listed in Supplemental Table 2.

### *Ppa-cat-1* reporter strain

To make the *cat-1p::GFP* transcriptional reporter (pMH1), 2 kb of the *cat-1* promoter was amplified from PS312 genomic DNA using RHL1236 and RHL1237. Using Gibson Assembly (E2611, New England Biolab), this *cat-1* promoter was introduced upstream of codon optimized *GFP* (pZH008) (Han et al., 2020), which includes the *rpl-23* 3’UTR and the plasmid backbone. This *cat-1p::GFP* construct (0.1 ng/µl), along with PS312 genomic DNA (80 ng/µl) and *egl-20p::RFP* (2 ng/µl) were individually digested with HindIII and assembled as the injection mix. It is noteworthy that while we also isolated several F_1_ animals with either head or vulva expression from injection mixes made with higher concentrations of the *cat-1p::GFP* plasmid (0.2-3 ng/µl), only the lowest injection concentration of the reporter plasmid (0.1 ng/µl) yielded *csuEx86* with stable reporter transmission for detailed characterization, suggesting that excess *cat-1* promoter DNA results in sterility or embryonic lethality.

### Live, fixed and immunostained worm imaging

Some worms were imaged with a Leica DM6000 upright fluorescence microscope using a 40x objective and photographed with Leica K5 cMOS camera, or a Zeiss Axio Imager with Apotome and processed with Zen software. Other transgenic worms were imaged with an Olympus BX60 upright fluorescence microscope and photographed with an Olympus Magnafire CCD camera. Some image stacks were created ‘manually’ with individual focal plane images from the standard compound microscope using FIJI (various versions). Some anti-serotonin preparations were imaged with a Nikon A5 laser scanning confocal microscope. Worms processed for HCR (see below) were imaged on a Zeiss Imager Z2 upright microscope equipped with Colibri 7 LEDs (Zeiss) using a 40x oil immersion objective lens, and an inverted Zeiss LSM 980 laser scanning confocal microscope using a 40x water immersion objective lens.

### Hybridization Chain Reaction

HCR was performed essentially as described previously (Ramadan and Hobert 2024). To help preserve the structure of the pharynx, centrifugation during all processing was reduced to 1000 g instead of 2000 g. Each HCR probe consisted of an initiator sequence (B1, B2, or B3), 2 nucleotide spacer (‘TT’), and 20 nt complementary sequence. Number of probes per gene varied depending on gene length (see Supplemental Materials for probe sequences), and probe sets were ordered from IDT as OligoPools. Hairpins complementary to each initiator sequence and conjugated with AlexaFluor (AF) 647, AF 546, or AF 488 were obtained from Molecular Instruments. Briefly, mixed-stage worms grown on NGM plates were harvested in M9 buffer and fixed with 4% paraformaldehyde/1x PBS and immediately frozen at −80°C overnight. Worms were subsequently thawed for 45 min, washed, and treated with 100 µg/ml proteinase K for 15 min at 37C to optimize tissue permeability, ‘stopped’ with glycine solution, and washed again. Samples were pre-incubated in probe hybridization buffer (PHB) at 37°C for 1 hour, then probe hybridization was carried out with probes in PHB overnight at 37°C. After hybridization, samples were washed with probe wash buffer and 5x SSCT before amplification using snap-cooled hairpins in amplification buffer, with overnight incubation in the dark at room temperature. Finally, samples were counterstained with DAPI and mounted in ProLong Gold Antifade Reagent for imaging. All microscopy was performed at 40x using a compound light microscope.

### Hybridization Chain Reaction with Immunohistochemistry

The HCR protocol above was modified to incorporate antibody staining for simultaneous detection of gene expression and protein localization, specifically to use a *C. elegans* myosin antibody (DSHB 5-6). Briefly, mixed-stage worms were harvested in PBS, fixed with 4% paraformaldehyde, and placed at 4°C overnight. Worms were washed with PBST, treated with glycine to quench PFA, and permeabilized with beta-mercaptoethanol (β-ME) at 37°C for 1 hour followed by proteinase K at 37C for 15 min to enhance tissue accessibility while preserving the myosin epitope. Samples were pre-hybridized in hybridization buffer at 37°C for 1 hour, then hybridized overnight with probes at 37°C. Worms were washed with probe wash buffer and 5x SSCT, then incubated with primary antibody against myosin (1:15 in PBST-A) overnight at room temperature. After washing with PBST-B, HCR amplification was performed with snap-cooled hairpins in amplification buffer overnight in the dark at room temperature. Samples were then washed with 5x SSCT, counterstained with DAPI, and incubated with AF488-conjugated secondary antibody (1:500 in PBST-B) overnight. Finally, after extensive washing with PBST-B, worms were mounted in ProLong Gold Antifade Reagent for imaging at 63x magnification using a Zeiss LSM 980 confocal microscope.

### Anti-Serotonin Immunostaining

We followed the immunostaining protocol in Loer and Rivard (2007). In brief, mixed-stage worms, in some cases containing mostly young adults, were washed with M9 from nearly saturated cultures on NGM plates and pelleted in 1.5 ml microcentrifuge tubes. Worms were fixed in 4% paraformaldehyde in 1X PBS (Thermo Fisher Scientific J61899 or prepared from PFA solid flakes) at 4°C overnight. Fixed worms were rinsed 3 times in 0.5% Triton X-100 / 1x PBS (PBSTx), then incubated overnight at 37°C in 5% beta-mercaptoethanol in 1% Triton X-100 in 0.1 M Tris pH 7.4 (TrisTx), rinsed 3 times TrisTx, then digested with Collagenase Type IV (Sigma-Aldrich C5138) at 2000 U/ml in 1 mM CaCl_2_/ Tris Tx for 30 min - 2 hours at 37°C with gentle mixing until adult worm cuticles just began to rupture. Other nematode species were digested for shorter or longer times as needed (Loer and Rivard, 2007; Rivard et al., 2011). After rinsing 3 times in PBSTx, worms were blocked for 1-5 hr with 1% BSA in PBSTx, then incubated in 1:100 Rabbit anti-5HT (Sigma-Aldrich S5545) in 1% BSA / PBSTx overnight at room temp, then rinsed in 0.1% BSA / PBSTx, followed by a secondary antibody (either 1:200 Goat anti-Rabbit IgG Alexa Fluor 647 or Goat anti-Rabbit IgG TRITC) overnight at room temp, then rinsed with 0.1% BSA / PBSTx for 1-2 hr with 3-6 changes. For most preparations, prior to mounting, DAPI was added (∼0.1 µg/ml final concentration).

### Cholinergic Immunostaining

To see cholinergic proteins, we followed the ‘freeze-crack’ immunostaining protocol described in (Duerr et al., 2008) (Duerr 2013). The antisera used was a mixture of several anti-*C. elegans* CHA-1 monoclonal antibodies (MAbs) and an anti-*C. elegans* UNC-17 MAb as described in the previous references. The overall staining pattern observed in *P. pacificus* was quite similar to that seen in *C. elegans*. In brief, synchronized adult worms were washed off plates with M9 into 1.5 microfuge tubes, and washed with a final rinse of deionized water, then pipetted on to a single untreated microscope slide. A second slide was placed staggered on the first slide and excess liquid was wicked away until adult worms were compressed and slightly flattened, and a few worms may have burst. These slides were then frozen on an aluminum block on dry ice. After at least 15 min, the slides were removed and immediately twisted apart. The slides were plunged into ice cold acetone for 5 min, followed by ice cold methanol for 5 min. Worms were then washed off the slides with 1x PBSTx into a 50 ml centrifuge tube on ice. The 50 ml tube was spun at 1000 g for 5 min to pellet the worms, and then the supernatant was removed. A small volume with the worms was transferred to a 1.5 ml microfuge tube in which the remaining antibody staining procedure was carried out.

### Three-dimensional reconstructions of neurons

We used volumetric reconstructions of *P. pacificus* neurons from ‘Series 14’ serial section EM reconstructions of neurons (Bumbarger et al., 2013) (Hong et al., 2019) (Cook et al., 2025) to assist in neuronal identification when viewing neuronal morphologies in immunostained and reporter fusion worms. We also examined 3D rendered maps of neuronal nuclei from ‘Series 14’ worms. In both cases, .obj files were exported from TrakEM2 (a FIJI plugin) which was used for neuronal tracing and synapse identification (Cook et al., 2025) and imported into Adobe Dimension (v. 3.4.11) to make volumetric renderings of neurons shown in this article. Images were typically labeled and/or adjusted in Adobe Illustrator (28.1) or Graphic Converter 11.

### CRISPR mutagenesis generated mutants

CRISPR/Cas9 mutagenesis (Han et al., 2020) (Nakayama et al., 2020) was used to generate mutations in *cat-1*(PPA18844), *dat-1*(PPA09932), and *cha-1*(PPA11636); for *cha-1,* an in-frame 2x-FLAG insertion into the C-terminus was also generated. crRNA and primer sequences are included in Supplemental Table 3. Generation of *cat-2, tdc-1* and *tbh-1* mutants is described in (Onodera et al., 2024).

#### *cat-1* alleles

Target crRNA, tracrRNA, and Cas9 nuclease were purchased from IDT Technologies (San Diego, CA, USA). crRNA (RHL1213) and tracrRNA were hydrated to 100 µM with IDT Duplex Buffer, and equal volumes of each (0.61 µl) were combined and incubated at 95°C for 5 minutes, then 25°C for 5 minutes. Cas9 protein (0.5 µl of 10 µg/ µl) was added, then the mix was incubated at 37°C for 10 minutes. *egl-20p::RFP* was used as a co-injection marker. To reach a final total volume of 40 µl, the Cas9-crRNA-tracrRNA complex was combined with pZH009 (*egl-20p::RFP*) DNA to reach 50 ng/µl final concentration using nuclease-free water. F_1_ progeny were screened for the presence of *egl-20p::RFP* expression in the tail and candidate F_1_’s were sequenced to identify heterozygotes (Nakayama et al., 2020). *csu115* has a 22 bp insertion while *csu116* has a complex 33 bp indel mutation, both frameshift mutations result in a premature stop codon in the first exon. Each allele was outcrossed three times to wildtype before characterization.

#### *cha-1* and *dat-1* alleles

indel alleles were generated during screens intended to epitope-tag the genes. *cha-1(ot5000)* is an in-frame knock-in of 2x FLAG before the last three amino acids, while *cha-1(ot5001)* contains an 8-bp deletion with a 68-bp insertion, and *cha-1(ot5002)* contains a 20-bp deletion with a 50-bp insertion. Both *ot5001* and *ot5002* are missing the last five amino acids (LLDRE) in the C-terminus due to indel frame-shifting. For the *dat-1* alleles, the CRISPR insertion site was in the predicted N-terminal region: *dat-1(pa507)* is a 303 bp deletion around the PAM site; *dat-1(pa508)* is a 441 bp deletion/12 bp insertion; both of these should be complete loss-of-function alleles. See Supplemental methods for detailed information on generation of these alleles.

### Behavioral assays: Egg-laying

We followed the egg-laying rescue protocol used by (Okumura et al., 2017). To synchronize worms, eggs were collected by allowing gravid adult hermaphrodites to lay eggs on food at 25°C for 4 hours before being removed; the progeny were then incubated at 20°C for ∼5 days. Eggs are typically laid at the 2-cell stage of embryonic development (Lewis and Hong 2014). The egg-laying defective (Egl) phenotype in adults was defined as having 8 or more eggs in the uterus with embryos in more advanced stages of development. Adults with this phenotype were counted on days 5 and 6 to determine the percent Egl for genotypes *PS312*, *cat-1(csu115)* and *cat-1(csu116)*. To test the effects of exogenous serotonin, adult worms were placed individually in 96-well flat bottom plates containing in each well 50 μl of 10 mM serotonin (Millipore Sigma 111019150) or serotonin hydrochloride (Sigma-Aldrich H9523) in M9 buffer.

### Behavioral assay: head oscillation suppression

Head oscillation suppression was assessed following the protocol described by (Alkema et al., 2005) with minor modifications. All gentle touch assays were performed on young adult worms at room temperature. Similar to the procedure used for *C. elegans*, suppression of head oscillation was evaluated by gently stroking just posterior to the posterior bulb with a fine eyelash while the animal was outside the bacterial lawn. For each genotype, 60 animals were assayed. Upon gentle touch, *P. pacificus* typically either paused or reversed; head oscillations were counted only when reversal occurred in response to gentle touch. An assay was scored as positive when head oscillations were successfully suppressed during backward movement. The proportions of positive and negative outcomes between the two groups were compared using Pearson’s chi-square test of independence. All expected cell counts exceeded 5, satisfying the assumption for the chi-square test; therefore, no continuity correction was applied. Statistical analyses were performed using Python (version 3.10.7).

### Behavioral assay: Nictation

Dauers from ∼2-weeks old, starved cultures were ‘chunked’ in ∼1 cm^2^ agar slabs onto NGM agar plates. After agar slabs were removed, we count the total number of dauers on each plate, which usually includes 20-60 dauer larvae. The nictation assay is commenced by adding a finger pinch of white sand evenly onto the dauer plates which triggers nictation. We counted the number of waving (nictating) dauer larvae after 30 minutes.

#### Nomenclature

Throughout the manuscript, *P. pacificus* genes are referred to *without* a *Ppa-* suffix; however, if necessary for comparison to another species such as *C. elegans (Cel-)*, the *Ppa-* suffix is added.

## Results

To understand how monoamines may be expressed and function differently in *P. pacificus* from the well-described *C. elegans,* we undertook a comprehensive identification of all neurons in *P. pacificus* that may synthesize and synaptically release the monoamine neurotransmitters serotonin, dopamine, tyramine and octopamine. We generated a *cat-1/ VMAT* reporter fusion, re-examined anti-serotonin staining and a previously described *tph-1* reporter fusion, and combined this with *in situ* hybridization chain reaction (HCR) for the monoaminergic marker genes *cat-1/VMAT* (all monoamines), *tph-1* and *mod-5/SERT* (5HT-specific), *cat-2/TH* (DA-specific), *tdc-1* (OA and TA), and *tbh-1* (OA-specific) (Figure 1), which are all represented as single orthologs in the *P. pacificus* genome. We also performed double- and triple-labeling for some gene combinations, including in worms with the *cat-1* or *tph-1* reporter transgenes to corroborate those expression patterns. (All results are summarized in Table 1.)

**Table 1.** Monoaminergic neurons in *C. elegans* hermaphrodite, canonical and other likely or possible cells. Pluralized names represent bilaterally symmetric pairs, singular names are unpaired of ventral nerve cord cells. [Canonical: Serotonin / 5-HT: NSMs, ADFs, AIMs*, RIH, HSNs; Dopamine: CEPVs, CEPDs, ADEs, PDEs; Tyramine: RICs, uv1s(4); Octopamine: RIMs] *AIMs uptake 5HT via MOD-5/SERT but do not express CAT-1/VMAT; therefore, they likely do not function as serotonergic neurons. Neurons expressing the *cat-1*/VMAT gene in *C. elegans* hermaphrodite but lacking expression of a typical or proper complement of neurotransmitter synthesis genes required for either serotonin, dopamine, tyramine or octopamine synthesis; these cells may use an unknown / novel monoamine (Wang et al., 2024): Cells expressing *cat-1*/VMAT alone; these cells may use betaine as a neurotransmitter (Wang et al., 2024).

| Neuron | Confidence in *Ppa* cell ID<sup>†</sup> | Neuron Type | *Cel* NT(s)<sup>‡</sup> | *Ppa* NT<sup>‡</sup> | anti-5HT staining | anti-5HT *mod-5*(-) | *tph-1* reporter | *cat-1* reporter | *cat-1* HCR | *tph-1* HCR | *mod-5* HCR | *cat-2* HCR | *tdc-1* HCR | *tbh-1* HCR |
| --- | --- | --- | --- | --- | --- | --- | --- | --- | --- | --- | --- | --- | --- | --- |
| I1 | ••• | Pharyngeal | ACh | 5HT* | ++ | same | - | - | + | +/- | +/- | - | - | - |
| NSM | ••• | Pharyngeal | 5HT | 5HT | +++ | same | +++ | ++ | ++ | ++ | + | - | - | - |
| RIP | ••• | Interneuron | ACh | 5HT* | +++ | same | +++ | ++ | ++ | ++ | + | - | - | - |
| ADF | ••• | Sensory | ACh / 5HT | 5HT | ++ / + | same | - | ++ | ++ | ++ | + | - | - | - |
| RIH | • | Interneuron | ACh / 5HT | 5HT | + / - | ? | - | + | - | - | - | - | - | - |
| RIM | ••• | Interneuron | Glu / TA | TA | NA | NA | - | ++ | ++ | - | - | - | +++ | - |
| RIC | ••• | Interneuron | Glu / OA | OA | NA | NA | - | + | ++ | - | - | - | + | ++ |
| VC | ••• | Motor neuron | ACh / 5HT<sup>a</sup> | 5HT<sup>a</sup> | +++ / ++<sup>a</sup> | same | + | + | ++ | + | +/-<sup>b</sup> | - | - | - |
| HSN | (absent?)<sup>‡</sup> | Motor neuron | ACh / 5HT | (absent?)<sup>‡</sup> | NA | NA | NA | NA | NA | NA | NA | NA | NA | NA |
| CEP | ••• | Sensory | DA | DA | NA | NA | - | ++ | ++ | - | - | +++ | - | - |
| ADE | ••• | Sensory | DA | DA | NA | NA | - | + | ++ | - | - | +++ | - | - |
| PDE | ••• | Sensory | DA | DA | NA | NA | - | + | ++ | - | - | +++ | - | - |
| uv1<sup>§</sup> | •• | uterine<sup>§</sup> | TA | Other MA?? | NA | NA | - | - | - | - | - | - | - | - |

Abbreviations: *Ppa* – *P. pacificus*; *Cel* – *C. elegans*; NT – neurotransmitter; NA – not applicable
All columns to the right of '*Ppa* NT' column refer to *P. pacificus* cell properties.
<sup>†</sup> Level of confidence in homology assignment / cell identification in *Ppa*: High / Certain: •••, Moderate: ••, Low •.
<sup>‡</sup> Neurotransmitter abbreviations: 5HT – serotonin, ACh – acetylcholine, DA – dopamine, Glu – glutamate, OA – octopamine, TA – tyramine. Note that many *P. pacificus* neurons are likely also to have small molecule co-transmitters; cells using the three other major neurotransmitters found in *C. elegans* (ACh, Glu, GABA) have mostly not yet been identified and/or published.
\* Neurotransmitter is different from that of the *C. elegans* homologous cell
<sup>a</sup> In *C. elegans*, only the two VCs closest to the vulva (VC4, 5) are weakly and unreliably 5HT+. In *P. pacificus*, all VCs are 5HT-IR+; vulva-proximal VCs (2, 3) are always strongly 5HT-IR whereas distal VCs (1, 4) are often moderately 5HT+.
<sup>b</sup> In *P. pacificus*, the two proximal VCs (2, 3) express *mod-5*; distal VCs (1, 4) appear not to express *mod-5*.
<sup>‡</sup> In *P. pacificus*, there are no sublateral 5HT+ cells near the vulva like the *C. elegans* egg laying neurons (HSNs).
<sup>§</sup> Neuroendocrine cells (neuron-like) or neurons, 2 bilaterally symmetric pairs directly lateral to the vulval pore.

**Table 2.** Nematode strains – *P. pacificus* and other species.

| **Species + Gene** | **Allele** | **Strain (Source)** |
| --- | --- | --- |
| *Pristionchus pacificus* | wildtype | PS312 |
| *Ppa-bas-1* | *tu629* | RS2955 (Okumura et al., 2017) |
| *Ppa-bas-1* | *tu630* | RS2956 (Okumura et al., 2017) |
| *Ppa-cat-1* | *csu115* | RLH245 (this study) |
| *Ppa-cat-1* | *csu116* | RLH246 (this study) |
| *Ppa-cat-1p::gfp* | *csuEx86* | RLH271 (this study) |
| *Ppa-cat-2* | *tu1023* | RS3150 (Onodera et al., 2024) |
| *Ppa-cha-1* | *ot5000* | OH16038 (this study) |
| *Ppa-cha-1* | *ot5001* | OH16039 (this study) |
| *Ppa-cha-1* | *ot5002* | OH16040 (this study) |
| *Ppa-dat-1* | *pa507* | LC179 (this study) |
| *Ppa-dat-1* | *pa508* | LC180 (this study) |
| *Ppa-mod-5* | *tu586* | RS2853 (Sommer lab) |
| *Ppa-mod-5* | *tu587* | RS2854 (Sommer lab) |
| *Ppa-tbh-1* | *cbh32* | MOK46 (Onodera et al., 2024) |
| *Ppa-tbh-1* | *cbh33* | MOK47 (Onodera et al., 2024) |
| *Ppa-tdc-1* | *tu1007* | RS3070 (Onodera et al., 2024) |
| *Ppa-tdc-1* | *tu1009* | RS3071 (Onodera et al., 2024) |
| *Ppa-tph-1* | *tu628* | RS2946 (Okumura et al., 2017) |
| *Ppa-tph-1p::rfp* | *tuEx257* | RS3066 (Okumura et al., 2017) |
| *Ppa-sem-4* | *tu314* | RS2003 (Photos et al., 2006) |
| *Acrobelloides nanus* | wildtype | ES501 (Wiegner and Schierenberg, 1999) |
| *Acrobelloides sp.* | wildtype | PS1146 (Felix and Sternberg, 1996) |
| *Plectus sambesii* | wildtype | ES601 (Lahl et al., 2006) |
| *Plectus sp.* | wildtype | WWL501 (P. Schiffer lab) |
| *Propanagrolaimus sp.* | wildtype | LC82 (Schiffer et al., 2014) |

**Table 3.**
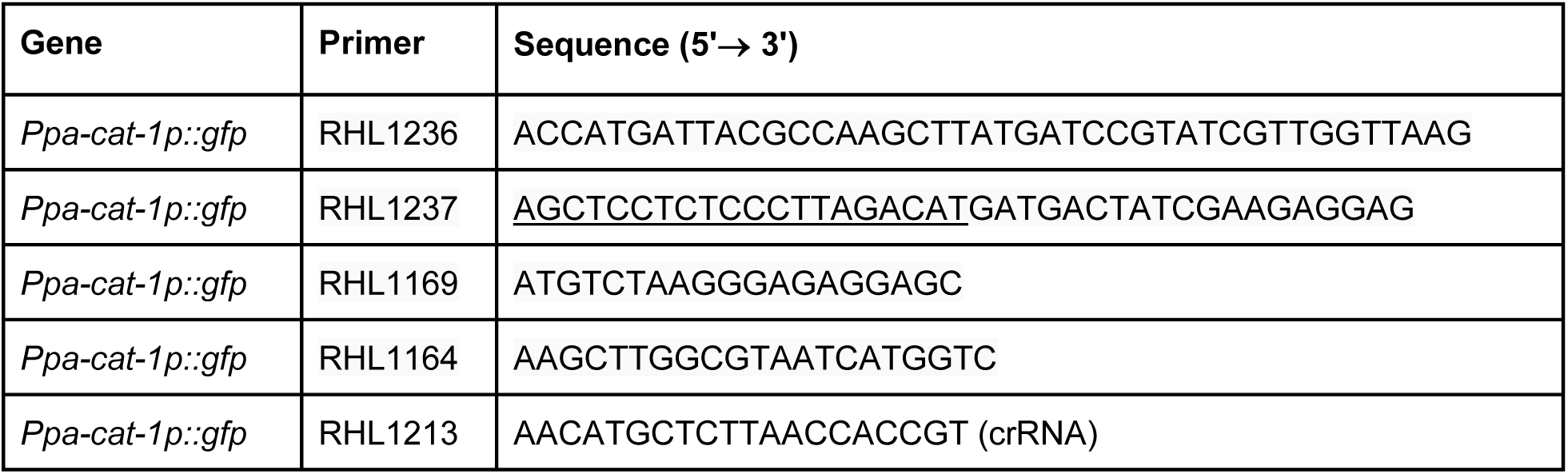
Primer sequences.

### Serotonin neurons in P. pacificus

While examining anti-serotonin staining in the *P. pacificus* head, we observed that, unlike in *C. elegans,* there are two strongly-staining somas anterior to the nerve ring, in the lateral region of the anterior ganglion (Fig. 2A, Suppl Fig S1). In *C. elegans*, all non-pharyngeal serotonin-immunoreactive (-IR) cells are located posterior to the nerve ring; only the pharyngeal NSMs are anterior to the nerve ring. We also examined a previously described *tph-1p::RFP* reporter fusion strain (Okumura et al., 2017) and observed that, along with strongly-expressing NSMs, a pair of somas anterior to the nerve ring also showed strong expression of the transgene (Fig. 2 B, C). Associated neurites showed a structure readily identifiable as belonging to the RIP neurons, having large flap-like endings terminating in the dorsal and ventral nerve ring (Fig. 2 D, E, G, H; (Cook et al., 2025). Unlike any other neurons in the anterior ganglion, RIP also extends its anterior neurite in the labial lateral bundle *contralateral* to the cell body by crossing over the dorsal midline. These anterior ganglion serotonin-IR and *tph-1* transgene-expressing cells were previously identified incorrectly as ADF neurons (Wilecki et al., 2015) (Okumura et al., 2017). ADF neurons, posterior to the nerve ring, are serotonin-IR but are not seen with this reporter. The lack of *tph-1* reporter expression in serotonin-IR head neurons such as I1 and ADF when HCR shows *tph-1* transcripts in the cells (see below) suggests that positive regulatory elements are missing from the construct.

**Figure 2.**
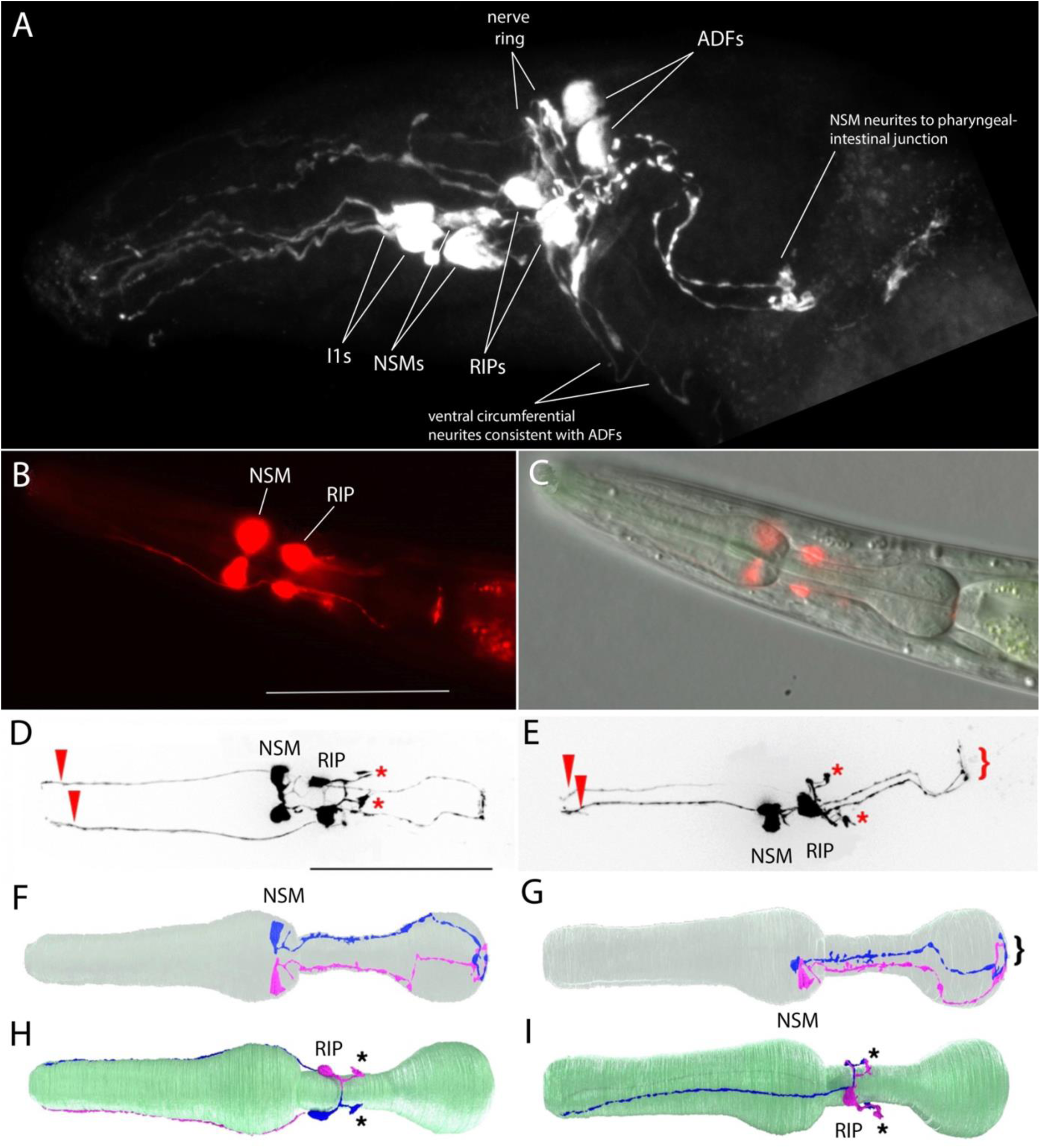
*P. pacificus* serotonin immunoreactive and *tph-1* transgene expressing neurons. Anterior is to the left in all panels. (A) Anti-serotonin staining in adult hermaphrodite head, approximately lateral view, MaxIP of confocal image. Four pairs of neurons and one unpaired neuron are apparent, as identified. Three pairs of neurites extend anteriorly to the tip of the pharynx or head, consistent with the known structures of I1, RIP and ADF. In this preparation, the pharynx is considerably kinked (as apparent by the bending of the posterior NSM neurites), making it appear that the ventral unpaired soma (RIH?) is anterior to the nerve ring when it is actually posterior to the nerve ring. (See supplemental figure S1 for a stained head showing this soma posterior to the nerve ring.) (B) Expression of *tph-1p::rfp* reporter in adult hermaphrodite head showing strong expression in a pair of pharyngeal neurons (NSMs) and anterior ganglion neurons in front of the nerve ring (RIPs). [Construct previously described in (Okumura et al., 2017).] Scale bars in B, D - 50 µm. (C) Same head as B, with DIC. (D) MaxIP with Apotome of *tph-1p::rfp* reporter showing neuronal morphologies, ventral view. Asterisks mark RIP endings in the nerve ring, arrow heads mark dendrites. (E) Left lateral view; otherwise, same as D. Bracket indicates NSM endings at pharyngeal-intestinal junction. (F-I) 3D rendering of neurons from serial section EM reconstruction (Cook et al., 2025); all left side neurons are rendered in magenta; right side neurons in blue. (F) NSML and NSMR inside a mostly transparent pharynx (gray-green), dorsal view. (G) NSMs, left lateral view. Bracket as in E. (H). RIP neurons, ventral view; pharynx (green). As in D, asterisks mark RIP endings in the nerve ring. (I) RIP neurons, left lateral view, as in H.

We also examined the expression of a *cat-1* transcriptional GFP reporter fusion with ∼2 kb upstream sequence. Although expression of this reporter was highly mosaic, we observed *cat-1p::GFP* expression in all post-hatching stages, including in dauer larvae; among the most robust expression of the *cat-1* reporter was in clearly identifiable serotonin-IR head neurons NSM, RIP, and ADF neurons (Figure 3 A-E; Suppl Fig S2). The NSMs exhibit extensive posterior processes that reach the base of the pharyngeal posterior bulb, like that observed with anti-5HT staining and as seen by serial EM reconstruction (Bumbarger et al., 2013; Cook et al., 2025); Figure 2 D-G). As with the *tph-1* reporter expression reported above, the RIP neurons’ neurite morphology revealed by the *cat-1::GFP* reporter was distinctive, standing out from other head neurons (Figure 2 D, E, H, I; Suppl Fig S3).

**Figure 3.**
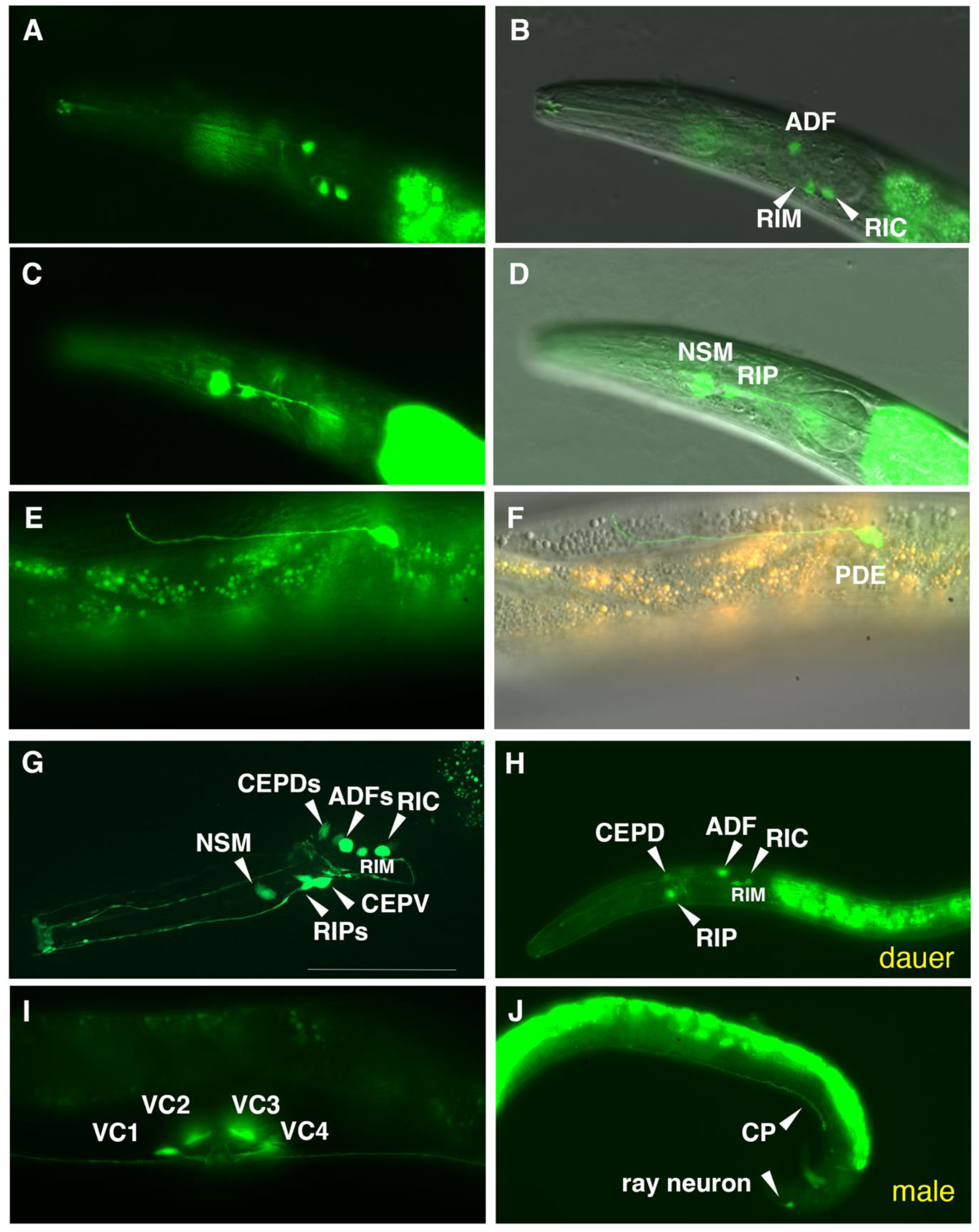
Expression pattern of a *cat-1p::gfp* reporter in *P. pacificus*. The ∼2 kb promoter fusion reporter *cat-1p::gfp* shows expression in known and other likely monoaminergic neurons. (A) GFP expression in the serotonergic amphid neuron ADF(AM9) and likely ring interneurons RIM and RIC. (B) Same with DIC. (C) GFP expression in pharyngeal neuron NSM and anterior ganglion RIP in the same worm. (D) Same with DIC. (E) Expression in mid-posterior lateral body dopaminergic PDE neuron with anteriorly extending dendrite ending near the dorsal midline. (F) Same with DIC. (G) MaxIP showing expression in likely CEPD and CEPV (dopaminergic) neurons, and NSM, RIP, ADF, RIM, and RIC neurons. (H) Expression in a dauer larva showing *cat-1p::gfp* that is similar to that seen at other developmental stages. (I) Expression in all four VC neurons (VC1-4) in the ventral nerve cord surrounding the vulva, young adult hermaphrodite. (J) A representative adult male shows expression in one likely ventral nerve cord CP motor neuron and a sensory ray neuron in the tail. Scale bar in (G) represents 50 µm in all panels. Because the reporter transgene is propagated as a complex extrachromosomal array resulting in mosaic animals from somatic loss of the array, we examined the expression pattern in many individual hermaphrodites for specific neuronal types across all post-larval developmental stages (n>100).

### Expression of serotonergic marker gene transcripts (tph-1, mod-5 and cat-1)

In the head, *in situ* hybridization via HCR for *tph-1* transcripts revealed strong signal in the pharyngeal NSM neuron and a weak signal, barely above background, in the I1 neuron, that was coincident with a strong *cat-1* HCR signal in both cells (Figure 4 A-C, G-I; Suppl Fig S3). We also observed *tph-1* co-expressed with *cat-1* laterally in the anterior ganglion, consistent with the RIP neuron (Fig. 4 A-C, G-I), and in a dorsolateral cell posterior and close to nerve ring, ADF (Fig 4 D-F). These *in situ* patterns match the 5HT antibody staining results.

**Figure 4.**
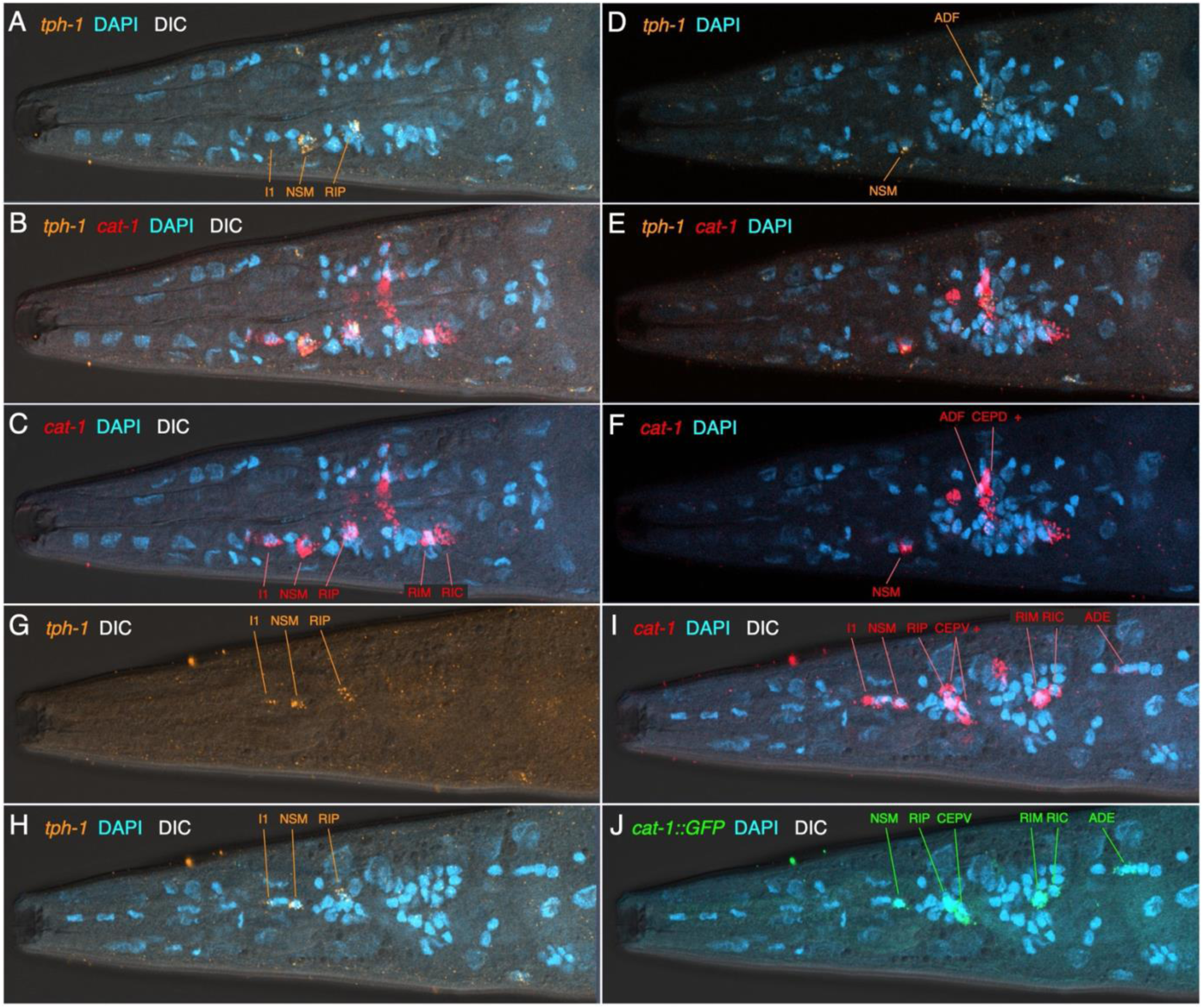
Co-expression of *tph-1* and *cat-1* transcripts in the head of *cat-1*::GFP strain. Anterior is to the left in all images; each is a single focal plane. Panels include (as noted in upper left corners) HCR for *tph-1* transcripts (orange) and *cat-1* transcripts (red), GFP (green), DAPI (cyan). (A-C) Ventro-lateral view (same focal plane) showing co-expression of *tph-1* and *cat-1* in serotonergic neurons I1, NSM, RIP, and lack of expression in other *cat-1*-positive lateral neurons (RIM, RIC). Outline of the pharynx is clear via the DIC, showing the locations of the cells (and DAPI-stained nuclei) in the ventral pharynx (I1, NSM) and anterior ganglion (RIP). (D – F) More dorsal view (single focal plane) showing co-expression of *tph-1* and *cat-1* in dorso-lateral serotonergic neuron ADF, and adjacent other *cat-1*-positive dorsal neurons (CEPD, + other unidentified nearby cell). Note that the strong signal from a ventral NSM is still visible in this focal plane. (G-J) Another ventro-lateral view (single focal plane) showing serotonergic neurons on the other side of the head, including (G) *tph-1* HCR signal alone (with DIC). (I, J) Match of c*at-1* HCR signal (I) with mosaic transgene *cat-1*::GFP (J) in a subset of *cat-1* transcript-positive cells. [In this preparation, HCR for the gfp gene was included, using an amplifier having the same excitation/emission properties as GFP. This is apparent by the particulate signal in the cells.] In the ventral region of the anterior ganglion, there are additional *cat-1* transcript-positive cells (+) adjacent to the CEPVs.

We did not observe *tph-1* transcripts in the other *cat-1*-positive cells in the head (described below). We observed a match between the mosaic *cat-1p*::GFP expression and *cat-1* transcript expression when HCR was performed in the *cat-1* reporter transgenic strain (Fig 4 I, J, Suppl Fig S3). In the adult hermaphrodite mid-body, we also observed *tph-1* transcript expression associated with four VNC nuclei (presumptive VC1-4) co-expressed with *cat-1* transcripts (Suppl Fig S4); like in the head, we did not see *tph-1* transcripts in any other *cat-1* transcript-positive cells, including any possible HSN homolog in the central body region. We observed *cat-1p::GFP* reporter expression near the vulva in the strongly serotonin-IR VC1-4; this mosaic expression was sometimes coincident with *cat-1* transcript expression by HCR (Suppl Fig S4). The *tph-1p::RFP* reporter was also sometimes expressed in VC1-4 neurons surrounding the vulva (Suppl Fig S5).

In *C. elegans,* three serotonin-IR head neurons posterior to the nerve ring, the bilaterally paired AIMs and unpaired RIH, require uptake for their serotonin – in *Cel-mod-5/SERT* mutants or in wild type worms treated with serotonin reuptake blockers, AIMs and RIH lose serotonin staining, while other serotonin neurons are not affected (Jafari et al., 2011). AIMs and RIH in *C. elegans* do not express the *tph-1* gene (Sze et al., 2000; Serrano-Saiz et al., 2017; Wang et al., 2024). To see whether any serotonin-IR neurons in *P. pacificus* depend on serotonin uptake, we examined serotonin staining in *mod-5* mutants. We found no change in staining among head and VNC serotonin-IR neurons, indicating none of the cells acquire serotonin solely via uptake (Suppl Fig S6).

As another typical marker of serotonin neurons, we also examined expression of *mod-5* transcripts by HCR. We observed clear expression, although often close to background levels, in the pharyngeal neurons I1 and NSM, in the anterior ganglion neuron RIP, and in a likely ADF (Suppl Figs. S7, S8); occasionally, we observed also expression above background in the 6-fold symmetric IL2 neurons (IL2D, IL2 lateral, IL2V, Suppl Fig. 8) In the midbody VNC, we observed *mod-5* expression in the VCs most proximal to the vulva (VC2, 3) that was coexpressed with *cat-1* transcripts; we observed no clear expression above background levels in the distal VCs (VC1, 4) that, like the proximal VCs, are strongly *cat-1* transcript positive (Suppl Fig. S9).

### Dopaminergic neurons and expression of dopaminergic markers (cat-2/TH and cat-1)

Neurons in the head and body of *P. pacificus* likely using dopamine have been identified previously via FIF staining and 5HTP-induced serotonin immunoreactivity (Rivard et al., 2010). In the head, we observed consistent *cat-1p::GFP* expression in identified bilaterally paired dopaminergic neurons CEPVs (ventral anterior ganglion) and CEPDs (dorsal) with cell bodies near the nerve ring and dendritic processes that span wider and extend more anteriorly than the ADF dendrites (Figure 3 G, H). We observed these same cells expressing *cat-2* transcripts colocalized with *cat-1* transcript expression (Figure 5). Further posterior in the head, laterally, alongside the posterior pharyngeal bulb, we observed another bilateral pair of dopaminergic cells, the ADEs (Fig 5 C-E). All of the *cat-2* transcript signals colocalized with *cat-1* transcripts. We also observed these 3 bilateral pairs of cells in preparations with *cat-2* HCR alone (Suppl Fig S10). Previously, staining with FIF, a rarely observed possible ventral unpaired DA neuron in the head was reported (Rivard et al., 2010); observed saw no evidence of such a cell via *cat-2* HCR.

**Figure 5.**
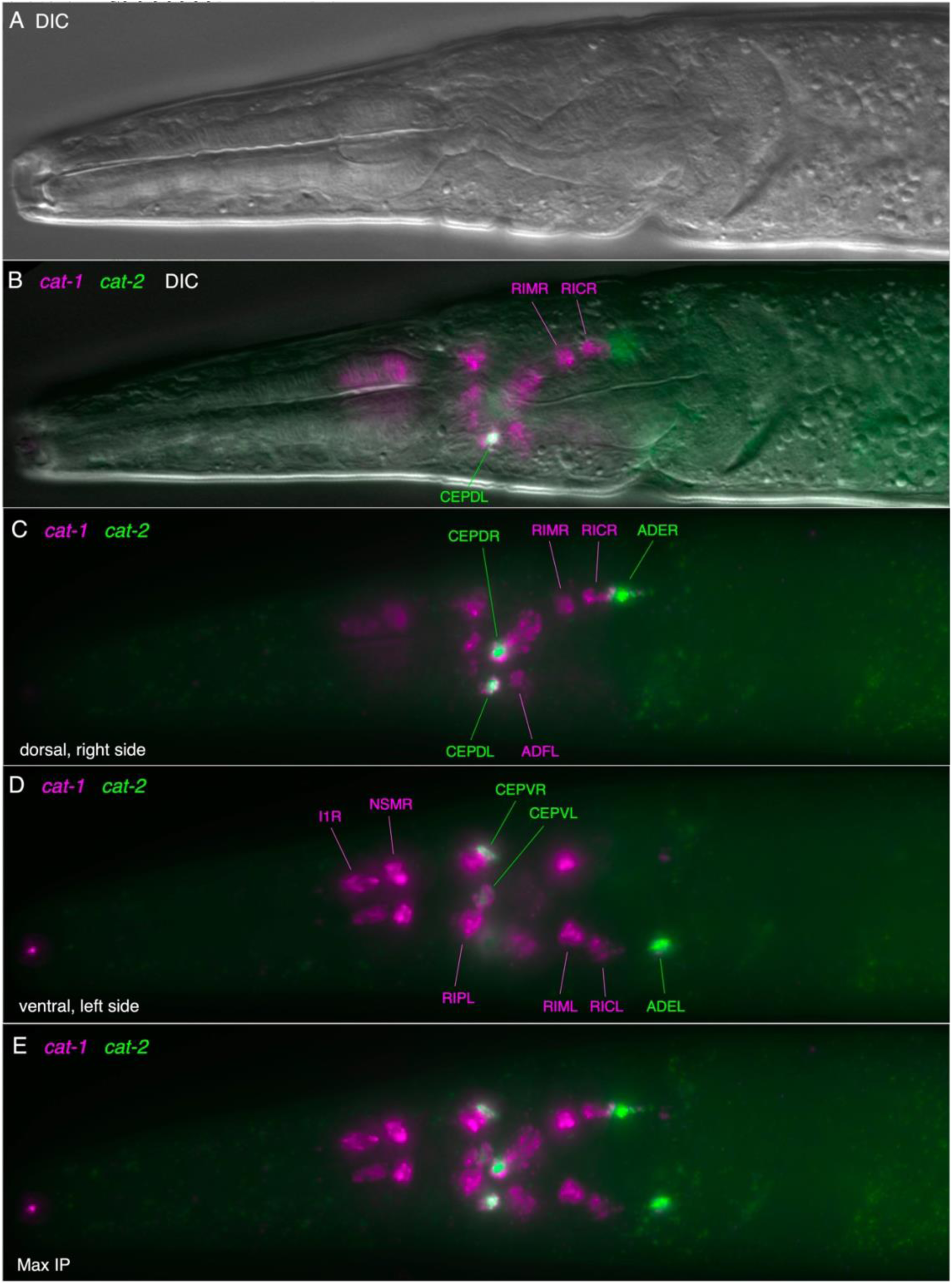
Co-expression of *cat-1* and *cat-2* transcripts in head dopaminergic neurons. Anterior is to the left in all images, all images of the same head. (A) DIC view of head, single focal plane to show outline of pharynx. (B) Single focal plane with *cat-1* (magenta), *cat-2* (green), DIC, showing colocalization in CEPVL. (C) MaxIP of several planes in dorsal and right side lateral; colocalization in CEPDs, ADER. (D) MaxIP of several planes in ventral and left side lateral; colocalization in CEPVs, ADEL. (E) MaxIP with all planes shown in C, D.

In the mid-posterior of the body, we observed expression in the bilaterally paired ciliated sensory neuron PDE located subdorsally in the body wall expressing the *cat-1::GFP* reporter (Fig. 3E-F). As reported previously, the *P. pacificus* PDE was noticeably different from that seen in the PDE of *C. elegans* and several other rhabditid nematodes: the cell’s dendrite is much longer and extends anteriorly before turning dorsally (Rivard et al., 2010); the soma is also located somewhat more posteriorly. We observed *cat-1* transcripts alone in the posterior body PDEs (Suppl Fig S10), colocalized with the *cat-1::GFP* reporter (Suppl Fig S11), and also observed PDE neurons co-expressing *cat-2* and *cat-1* transcripts (Suppl Fig S12).

### Other possible dopaminergic cells in the mid-body

Because serotonergic and dopaminergic neurons share the enzyme used in the second step of synthesis (AAADC / BAS-1, Figure 1), if worms are treated with the immediate precursor of serotonin (5HTP, 5-hydroxytryptophan), dopaminergic neurons can take up 5HTP and convert it into serotonin (5HT), becoming serotonin immunoreactive. FIF typically shows DA somas only and rarely neurites; whereas DA cells rendered serotonin-IR via 5HTP treatment can have well-stained neurites. Therefore, this technique is superior for seeing DA cells with neurites particularly if they are well-isolated from 5HT cells (which also stain with anti-5HT). Using this technique, we previously were able to observe the distinctly different sensory dendrite structure of the *P. pacificus* PDE neuron (Rivard et al., 2010). In such 5HTP-treated preparations, we also noted two bilateral pairs of cells in the lateral body wall flanking the vulva, which extend short processes toward the midline at about the same region where serotonergic VC neurons innervate vulval muscles (Suppl Fig S13, S14A). By the location and morphology of these possibly dopaminergic cells, they are likely homologs of the *C. elegans* neuroendocrine ‘uterine ventral’ uv1 cells. We observed that both uv1s and PDEs require *bas-1/AADC* function to convert 5HTP to serotonin: in *bas-1* mutants, there is no serotonin-IR in uv1s and PDEs after 5HTP exposure (Suppl Fig S14B). Staining of uv1s and PDEs is unchanged in *tdc-1* and *tbh-1* mutants (Suppl Fig S14C, D). Therefore, it is possible that uv1s use some other amine as a neurotransmitter. In these experiments, we did note some serotonin-IR staining of VC neurons in *bas-1* mutants (Suppl Fig S14B); this staining does not require addition of 5HTP. Therefore, the available *bas-1* mutants may have some residual function; both mutations disrupt only the downstream copy of a tandem duplication of the locus (Suppl Fig S15).

In *C. elegans,* the uv1 cells are tyraminergic; egg-laying is inhibited by tyramine (Alkema et al., 2005). Surprisingly, we observed no expression of either *cat-2* or *cat-1* transcripts in the putative uv1s, suggesting that these cells are not actually dopaminergic. In *C. elegans*, the tyraminergic uv1 cells also do not express *cat-1,* indicating non-vesicular release of the neurotransmitter (Wang et al., 2024). Although we did not observe *cat-1* transcripts in these cells, we did observe *cat-1* expression in nearby vulval cells (described below).

### Tyraminergic & octopaminergic neurons and marker expression (tdc-1, tbh-1 and cat-1)

On each side of the *C. elegans* head, alongside the pharyngeal posterior bulb, is a single tyraminergic ring interneuron pair (RIM L/R) and an adjacent octopaminergic neuron pair (RIC L/R); homologous bilaterally paired neurons are readily identifiable by serial section EM reconstruction in *P. pacificus* [(Cook et al., 2025), Suppl Fig S16]. We observed several head neurons posterior to the nerve ring expressing *cat-1p::GFP*, including laterally-positioned cell bodies that are likely RIMs and RICs (Fig. 3 B,G,H). RIM and RIC cell bodies are distinguished by their conserved position and their neurites extending posteriorly and ventrally that loop back before joining the nerve ring (Fig. 3G, Suppl Fig S16). Using HCR, we observed a strongly *tdc-1* transcript-positive cell on each side consistent with RIM, and immediately posterior to that, another *tdc-1* transcript-positive cell that also expressed *tbh-1* transcripts, RIC (Figure 6). The presumptive RIC cell consistently had a lower *tdc-1* signal by HCR than the RIM cell. HCR for *cat-1* transcripts posterior to the nerve ring included a likely RIM and RIC (Figures 4, 5, 9); in some preparations we observed co-localization of *cat-1* transcripts with *tdc-1* and/or *tbh-1* (Suppl Fig S17).

**Figure 6.**
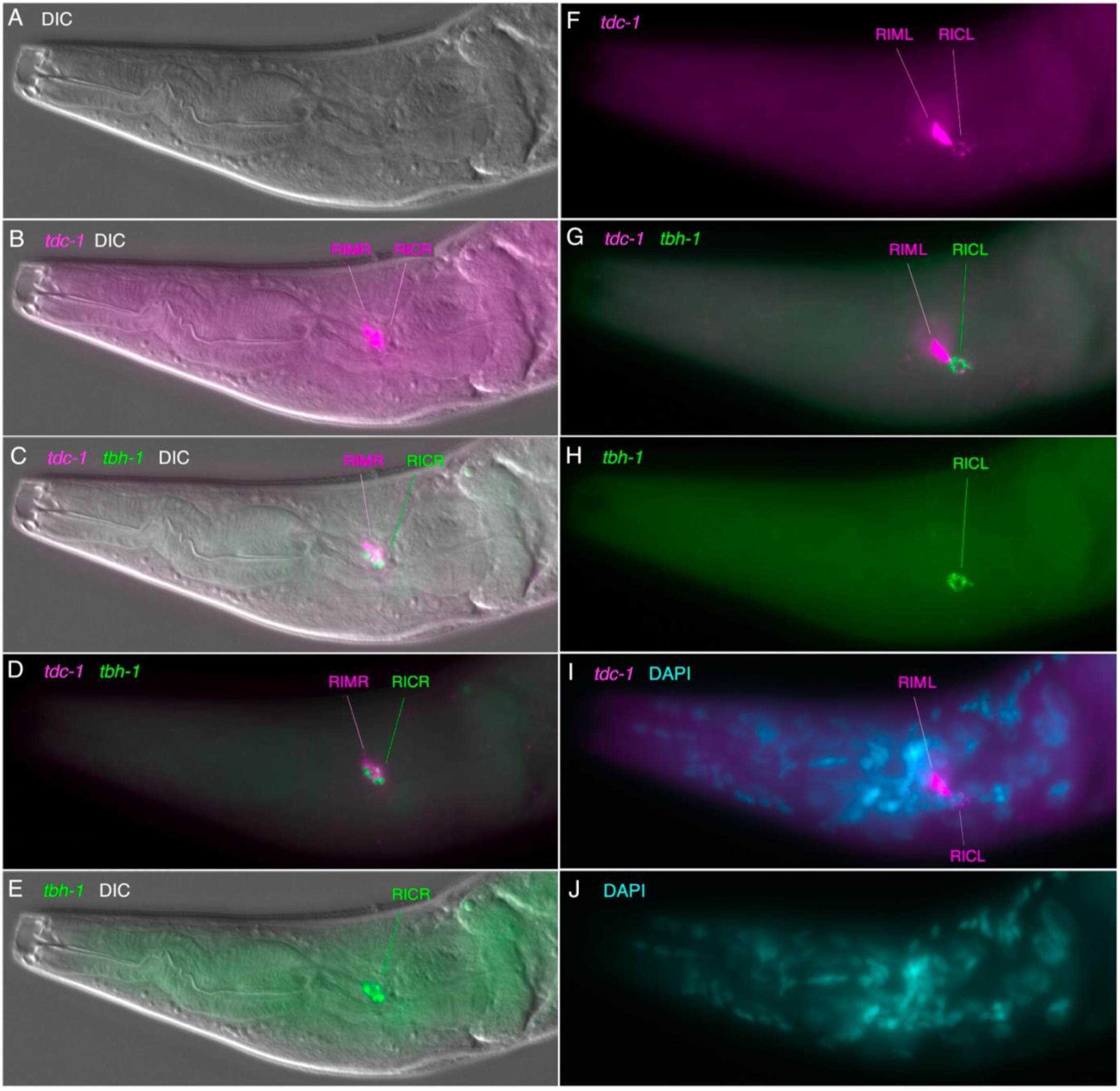
Expression of *tdc-1* and *tbh-1* transcripts in the head. Anterior is to the left and ventral down, all images of the same head; *tdc-1* transcripts (magenta), *tbh-1* transcripts (green). (A-E) Same single focal plane, right side. (A) DIC view, to show outline of pharynx. (B) *tdc-1* and DIC, showing two cells in the right lateral ganglion, presumptive RIMR and RICR. (C, D). Same focal plane as B, *tdc-1*, *tbh-1*, with (C) and without DIC (D) showing colocalization in the posterior cell (RICR). (F-J) Same single focal plane, left side. (F) *tdc-1*, showing two cells in the left lateral ganglion, presumptive RIML and RICL. (G). *tdc-1*, *tbh-1* showing colocalization in the posterior cell (RICL). (H) *tbh-1* alone, showing signal *only* in the posterior cell (RICL). (I) *tdc-1,* DAPI (blue, due to cyan + magenta background). (J) DAPI (cyan) alone, showing adjacent nuclei associated with *tdc-1* signal (I).

### Non-neuronal cells in the body expressing tdc-1, tbh-1 or cat-1 transcripts

Within the body, we also observed expression of *tbh-1* transcripts in the gonad, and *tdc-1* and *cat-1* expressed separately in different adjacent cells near the vulva in young adults. In the two arms of the hermaphrodite gonad, in the dorsal, near to the sharp ‘elbow’ where the gonadal arms bend down to the ventral, we observed a region of strong *tbh-1* expression in likely somatic gonadal sheath cells (Figure 7), which differ morphologically and positionally from those in *C. elegans* (Rudel et al., 2005). Close to the uterus in the ventral gonad, we observed moderate-to-weak expression of *tbh-1* in cells between the gonad bend and vulva that is likely part of the spermatheca (Figure 7, Suppl Figs S18, S19). In adults with developing embryos and clear sperm nuclei, this staining appeared to be associated with cells in the constriction just distal to the sperm nuclei. Interestingly, we did not see expression of *tdc-1* or *cat-1* in likely uv1 cells lateral to the vulva, which in *C. elegans* are tyraminergic (i.e., the cells seen in Suppl Fig S13).

**Fig. 7.**
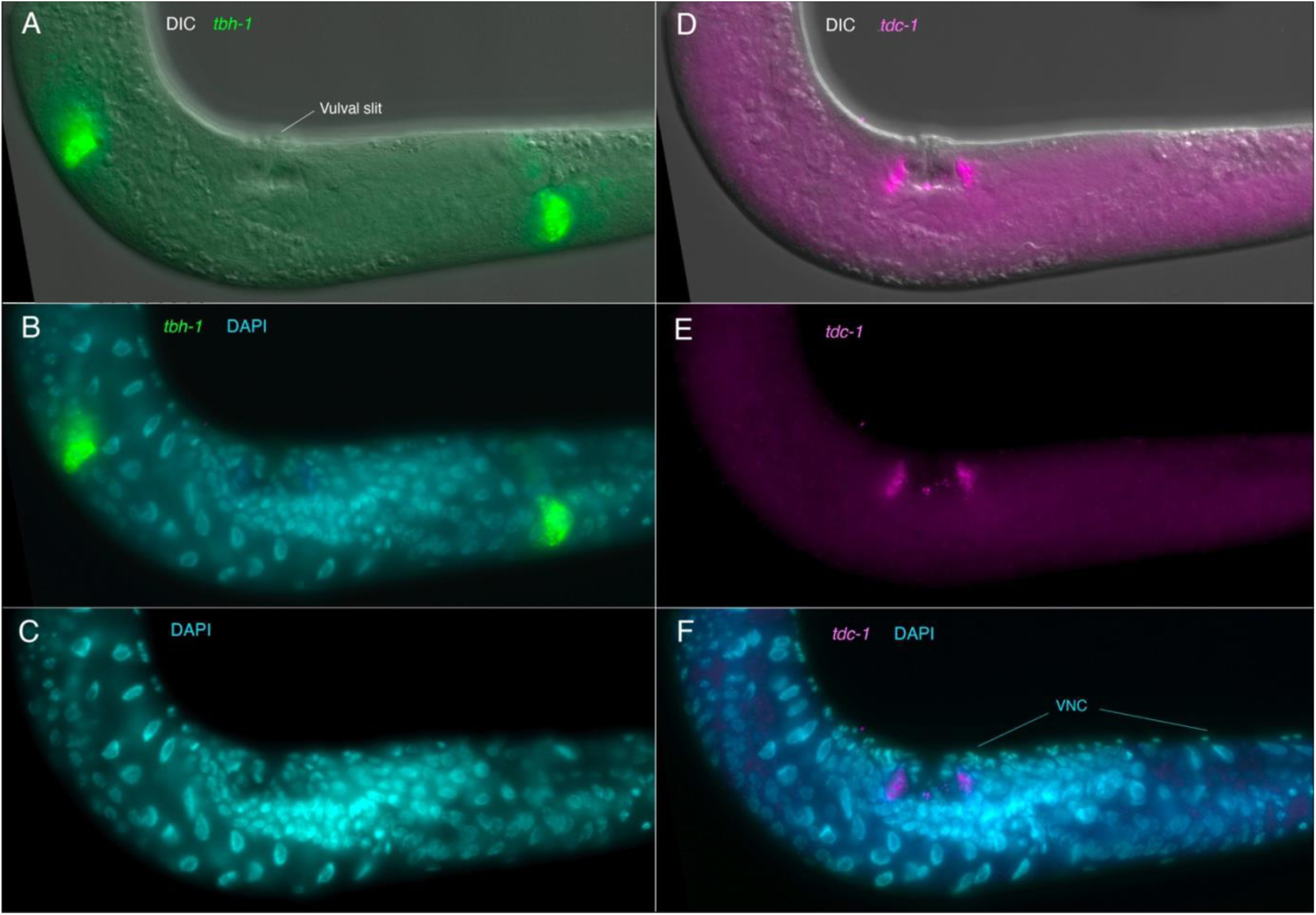
Expression of *tdc-1* and *tbh-1* transcripts in the non-neuronal gonadal & vulval cells. Anterior is up and to the left, ventral up as seen by the location of vulva and ventral nerve cord (VNC). (A-C) Same focal plane. (A) Expression of *tbh-1* transcripts (green) in the gonad, likely in gonadal sheath cells, with DIC. (B) *tbh-1* (green) with DAPI staining of nuclei (cyan). (D-F) Same focal plane. (D) Expression of *tdc-1* transcripts (magenta) in vulval cells, with DIC. (E) *tdc-1* transcripts alone (magenta). (F) *tdc-1* (magenta) with DAPI staining of nuclei (cyan/blue). See also Supplemental Figures for closeups of gonadal and vulval expression.

Upon closer examination, *tdc-1* and *cat-1* transcripts appear to be expressed in separate, adjacent cells (Figure 8, Suppl Fig S20); both the *tdc-1-* and *cat-1-*expressing non-neuronal cells are also clearly distinct from the four *cat-1-*expressing VC neurons in the VNC (Suppl Fig S21). We subsequently co-stained worms with *tdc-1* and *cat-1* HCR and a *C. elegans* myosin antibody that strongly stains vulval muscle, which showed clearly that the *tdc-1* and *cat-1* transcript-positive cells are not vulval muscle, but are likely vulval epidermal cells (Figure 8, Suppl Fig S20, S20). The fact that the cells are anterior-posterior and left-right symmetrical supports their identification as vulval cells, given the manner in which vulval precursor cells divide to generate the mature vulva (Sommer and Sternberg, 1996). Immunostaining of TBH-1 protein shows gonadal sheath expression in *C. elegans* (Alkema et al., 2005); therefore, gonadal sheath cell expression of *tbh-1* is conserved between the two species, despite the considerable differences in gonad structure. Conversely, antibody staining of *C. elegans* TDC-1 protein seen in oocytes within the gonad is not conserved – no *tdc-1* transcript expression is seen in the gonad in *P. pacificus*.

**Figure 8.**
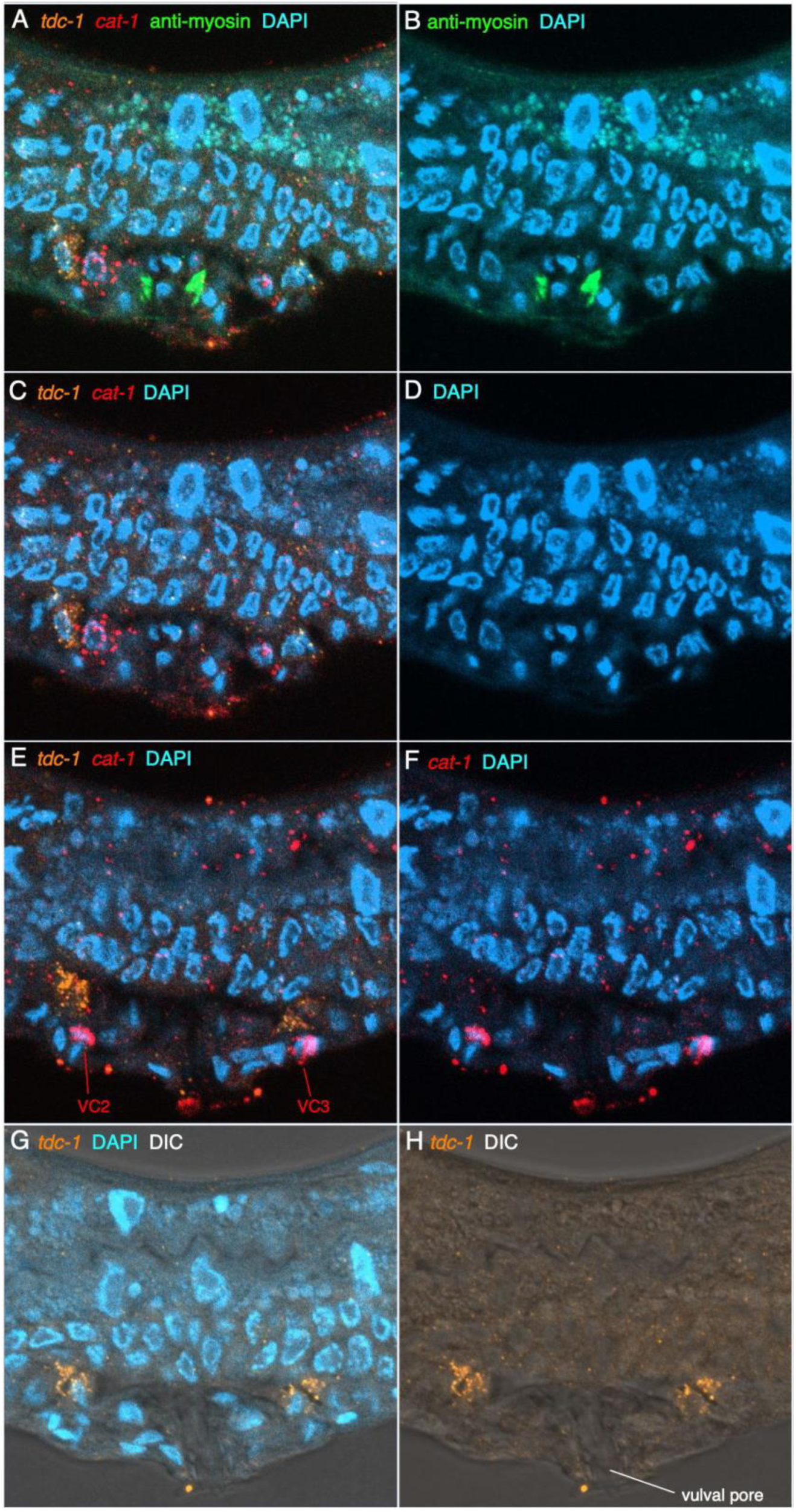
*cat-1-* and *tdc-1*-expressing cells near the vulval opening are not vulval muscles. (A-D) same focal plane, more superficial. On the left side, adjacent *tdc-1* (orange) and *cat-1* (red) expressing cells, presumptive vulval epidermal cells, are particularly apparent, and distinct from vulval muscles showing strong myosin immunoreactivity (green) centered and close to the vulva. (The *tdc-1* and *cat-1* expressing cells can also be seen on the right side but are less clear.) (E, F) deeper focal plane at level of the VNC, showing VC neurons that flank the vulval pore. (G,H) slightly deeper focal plane showing continuation of tdc-1 transcript-positive cells anterior and posterior to the vulva.

### Other cat-1-expressing cells in the head

We observed additional *cat-1-*transcript-positive cells in the head that did not match up with known serotonin, dopamine, tyramine or octopamine cells expressing marker genes for those neurotransmitters (Figure 9). In the dorsal anterior ganglion, a bilateral pair was seen just anterior to the nerve ring, possibly OLQD neurons (Figure 9C, D; Suppl Fig S22). In the lateral ganglion, another more weakly-stained bilateral pair was seen adjacent and/or ventro-lateral to presumed CEPD and ADF cells (Figure 9C, D); possible candidates include URX or AWA neurons. In *C. elegans,* none of the homologous neurons expresses *cat-1* as assessed by either fosmid-based reporter or the least stringent scRNA expression threshold (Wang et al., 2024).

**Fig. 9.**
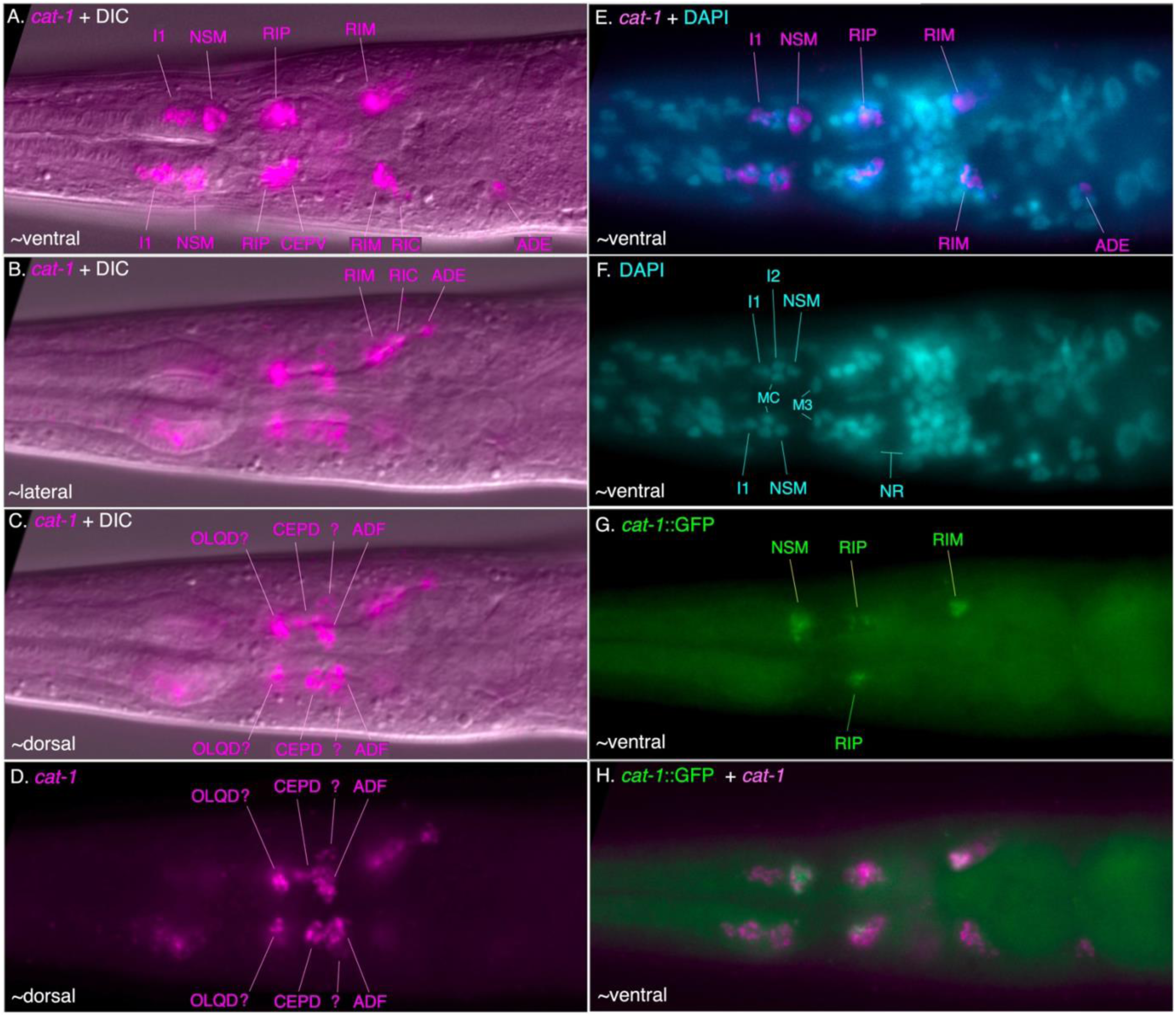
Expression of *cat-1* transcripts in the head of *cat-1*::GFP strain. Anterior is to the left in all images; each is a single focal plane. (A – D) HCR for *cat-1* transcripts (magenta) in 3 different focal planes – ventral / ventro-lateral (A), lateral (B), dorsal / dorso-lateral (C, D). Both *cat-1* HCR fluorescence and DIC (A-C); *cat-1* alone (D), same focal plane as C. Panels E – H are the same focal plane as panel A. (C, D) Dorso-lateral unidentified *cat-1-* transcript positive cells anterior and posterior to the nerve ring – possible OLQDs in anterior ganglion, and unidentified cell adjacent to CEPD and ADF; candidates include URX, AWA neurons (see also Supplemental Figure X). (E-F) Nuclear staining with DAPI shows stereotyped ‘diamond’ arrangement of ventral pharyngeal anterior bulb nuclei as labeled. *cat-1* HCR fluorescence and DAPI (E); DAPI alone (F). NR = nerve ring location. (G, H) Ventro-lateral GFP+ cells also show *cat-1* HCR fluorescence. Transgene expression *cat-1*::GFP alone (G); *cat-1*::GFP and *cat-1* HCR fluorescence colocalization.

### Monoaminergic neurons in P. pacificus males

Although our examination of monoaminergic neurons was primarily focused on hermaphrodites, we also examined males in the *cat-1*::*GFP* reporter strain. Males have four strongly serotonin-IR CP neurons and two weakly-stained CA neurons in the VNC, with a single bilaterally paired serotonin-IR sensory ray neuron (RN) in the tail (Loer and Rivard, 2007). We saw *cat-1* expression in occasional presumptive CP neurons in the VNC and in RNs in the tail (Figure 3J). Although we observed only a single RN in any given male examined (∼10 males with an RN / 56 males examined), their variable cell body positions in the tail suggest more than one RN type expresses *cat-1p::GFP.* In *C. elegans,* among the 18 bilateral pairs of RNs, there are 8 that use monoamines: 3 serotonergic, 3 dopaminergic, and 2 tyraminergic (Wang et al., 2024). Neurotransmitters for other *P. pacificus* ray neurons have not yet been identified, but the pattern of *cat-1::GFP* expression suggests there are also many monoaminergic RNs like in *C. elegans*.

### P. pacificus neurons I1 and RIP are ancestrally serotonergic

Differences in serotonin-IR neurons in *P. pacificus* and *C. elegans* prompted us to examine other nematode species of known relationship to assess which of these patterns are ancestral versus derived. Patterns of serotonin-IR neurons in several other nematode species have been reported previously (Henne et al., 2017); however, considering the clear identifications of several serotonin-IR neurons in *P. pacificus,* we decided to re-examine previous findings, and look at the patterns in additional free-living species, especially among outgroups of those examined previously, almost all of which are members of nematode ‘clade V’ [B - (Blaxter et al., 1998)] or ‘clade 9’ [H - (Holterman et al., 2006)]. We performed an extensive re-examination of head images from all the species we analyzed previously (Loer and Rivard, 2007) (Rivard et al., 2010), including larvae, adult male, female/hermaphrodite, and detached heads in which the sex could not be determined (Figure 10, Suppl Figs S23, S24, S25). As we have previously observed, patterns of serotonin-IR head cells in larvae and adults of different sexes were the same, except for *Poikilolaimus oxycera*, which was excluded from our analysis here. There were some slight differences in the relative positioning of cells along the anterior-posterior axis in larvae vs. adults, consistent with growth of head structures during larval development. Although we were not always able to be certain about the ‘complete’ pattern of cells given staining variability, particularly among the weakly-staining cells, we were able to see consistent patterns indicating the likely presence or absence of serotonin-IR in most cells known from *C. elegans* and *P. pacificus*.

**Figure 10.**
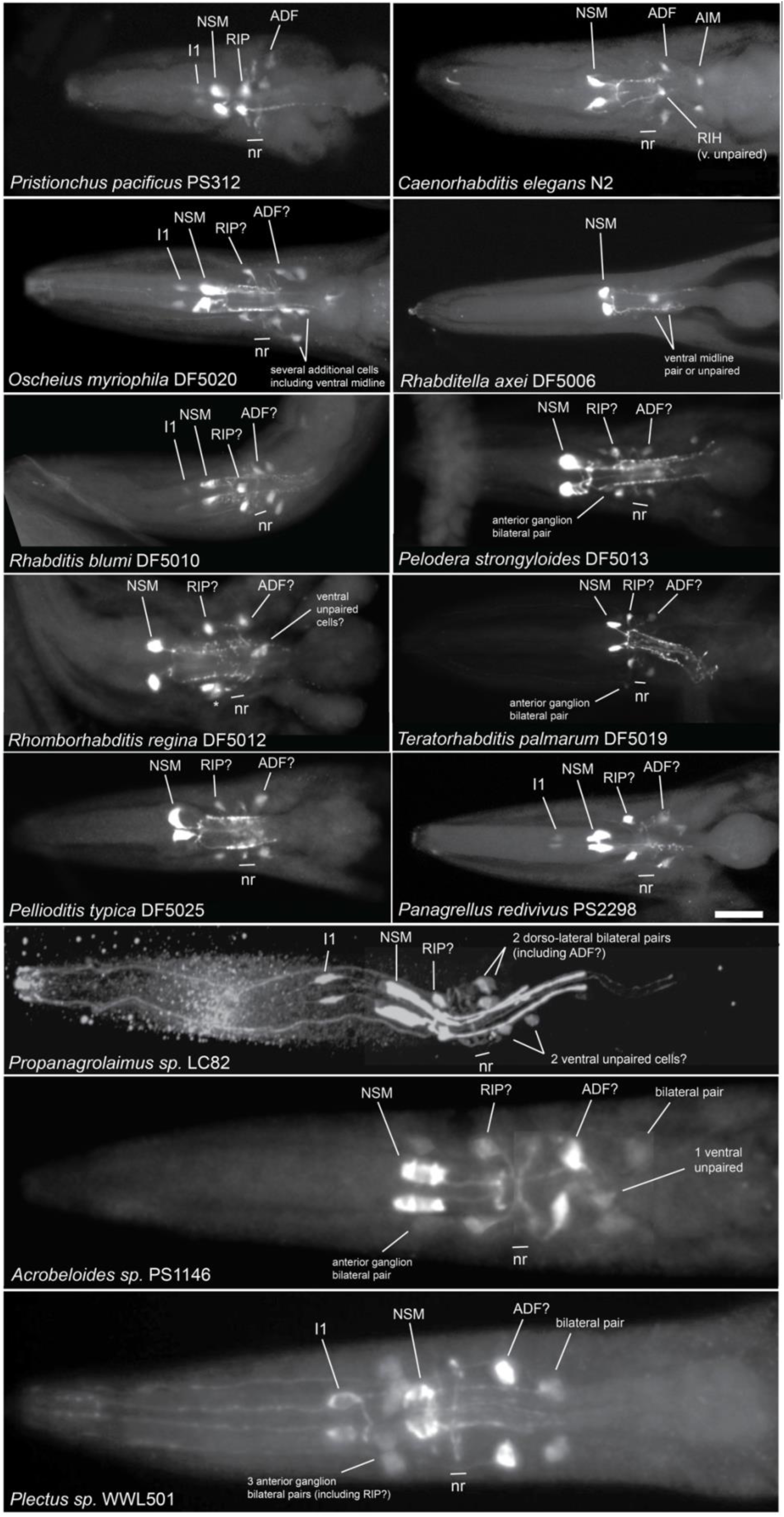
Serotonin staining in 13 different nematode species. Serotonin immunoreactivity in the heads of representative species examined or re-examined. Likely identifications of cells based on location are indicated. Cells in the pharynx directly anterior to NSMs and sending a neurite anteriorly to the tip of the pharynx were identified as I1s. Single pairs of neurons located dorsolaterally in the anterior ganglion were identified as likely RIPs. In the case of *Acrobeloides* and *Plectus* species, the presence of multiple bilaterally-paired neurons in the anterior ganglion suggested that the cells might be members of a 4- or 6-fold symmetric class of neurons, possibly not including RIPs; the numbers are listed under ‘other’ in Figure 14, which includes the two possible RIPs.

We also newly examined three more species in clade IV (B) – *Propanagrolaimus* [clade 10 (H)] and 2 *Acrobeloides* [clade 11 (H)], and two species of *Plectus* [clade 6 (H)]. The pattern of head neuron staining we observed in many species was more like that of *P. pacificus* than of *C. elegans.* While the pharyngeal NSM neurons remain the most easily recognizable and clearly conserved serotonin-IR neurons in all the species examined, we also found species in clades 9, 10 and 6 (H) that had pharyngeal neurons just anterior to the NSMs that are very likely I1 homologs (Figures 10, 11), often having anteriorly-directed neurites within the pharynx. We also observed lateral anterior ganglion neurons (anterior to the nerve ring) that are likely RIP neurons. In most cases, at least some stained worms had visible dendrites extending anteriorly as expected for RIPs. In several species, including *Acrobeloides* and *Plectus*, we found 1 – 3 additional bilateral pairs of cells in the anterior ganglion, or even further anterior. In some cases (e.g., *Plectus sp.* WWL501 and *Acrobeloides nanus* ES501), the 3 bilateral pairs appear to be 6-fold symmetric, suggesting that one pair of the cells may *not* be RIP (Figure 10). Assuming conservation of cell numbers and basic location, there are three sets of 6-fold symmetric neurons in the anterior ganglion (in both *C. elegans* and *P. pacificus*): IL1s, IL2s, and OLLs + OLQs. Given the further anterior placement of the dorsal and ventral pairs seen in *Acrobeloides*, this may indicate staining in the IL2 neurons, since such anterior placement of IL2D & IL2V is seen in *P. pacificus*.

**Figure 11.**
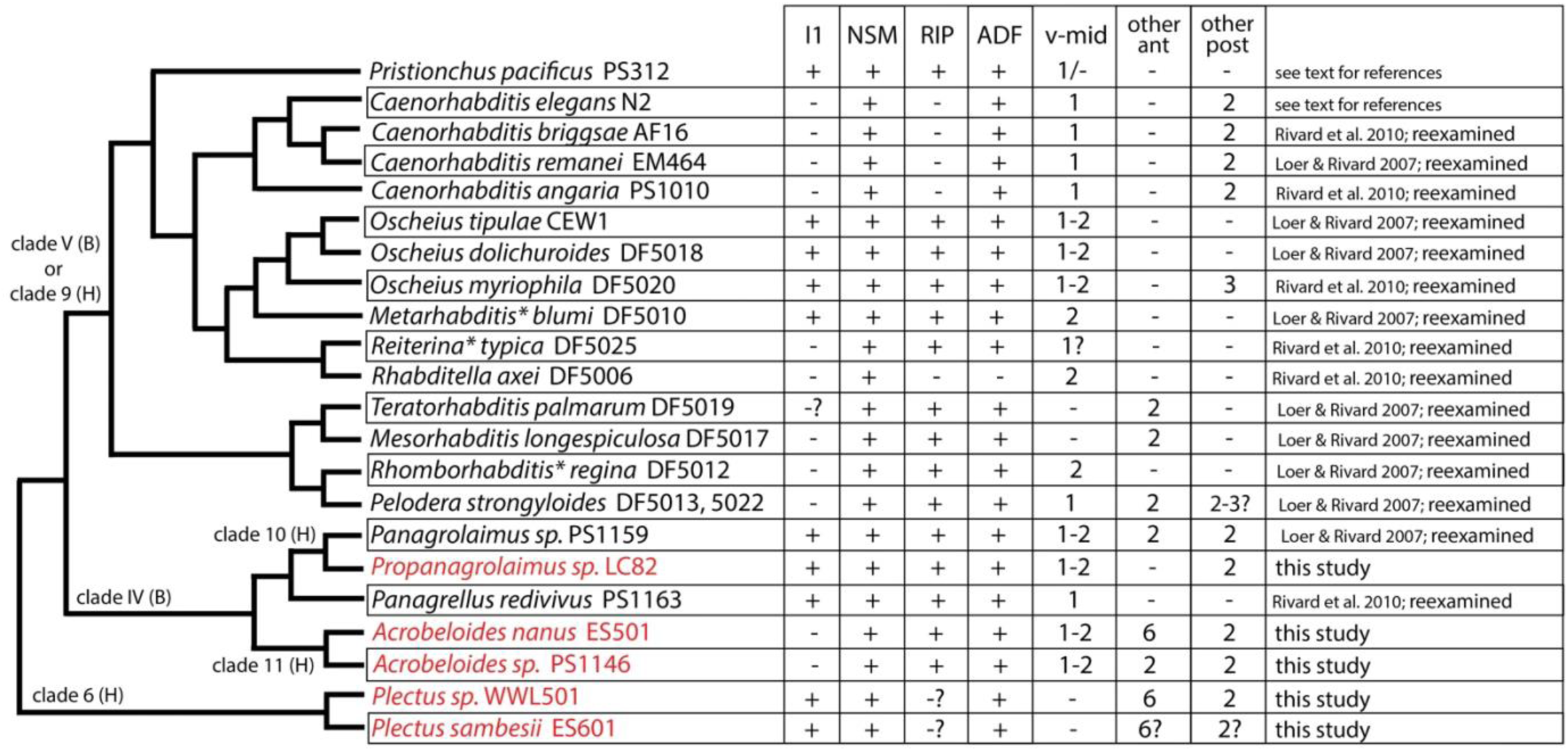
Cladogram of species with 5HT cells; serotonin-IR in I1 and RIP neurons is ancestral. Cladogram of species examined or re-examined for serotonin-IR head neurons. (Branch lengths do not indicate phylogenetic distance.) Symbols: ‘ +’ = bilateral pair of cell(s) present; ‘ -’ = cell(s) absent. ‘ -?’ = indicates that a possible matching cell was seen in only a few of the heads examined. For these purposes, if a single pair of cells was observed in the correct location (e.g., a likely RIP in the lateral portion of the anterior ganglion), then the cell is marked as present. Other columns: v-mid = ventral midline cells posterior to the nerve ring – either 1 unpaired cell or 2 cells (that might be a bilateral pair or 2 unpaired cells); in *C. elegans,* this cell is RIH. Other ant = additional unidentified cells anterior to the nerve ring (i.e., not RIP). Other post = additional unidentified cells posterior to the nerve ring (i.e., not ADF). These are all cells not matching an identified serotonin-IR cell known from *C. elegans* or *P. pacificus*. For numbers of ‘other’ cells, a ‘2’ indicates a single bilaterally symmetric pair. For ‘other ant’ cells in clades 6 and 11 (H), these appear to be 6-fold symmetric cells, and may be IL2 neurons. For ‘other post’ cells, in the genus *Caenorhabditis*, these are AIM neurons, seen only in this genus, with the possible exception of *O. myriophila* and *P. strongyloides.* For ‘other post’ cells in clades 6, 10, and 11 (H), these are bilaterally symmetric pairs of cells adjacent to ADF in the dorsolateral ganglion. Species in red are presented for the first time in this study. Species marked with an asterisk (*) have been given new genus names (i.e., since Loer & Rivard, 2007 and Rivard et al., 2010); the matching species can be identified by the strain number. Clade designations (left side of cladogram) are from either (H) Holterman et al., 2006 or (B) Blaxter et al., 1998; *Plectus* is not within a B-designated clade number.

In most species, we also observed at least one bilateral pair of 5HT-positive somas located dorso-laterally and posterior to the nerve ring, which send a neurite anteriorly to the tip of the ‘nose,’ and which show ventrally directed circumferential axons typical of ADF amphid sensory neurons; in a few cases, we could see a ‘double neurite’ ending typical of ADF as seen in both *C. elegans* and *P. pacificus* (Hong et al., 2019). The presence of serotonin in apparent ADF neurons is nearly as strongly conserved as with the NSMs – we found only one species lacking serotonin-IR presumptive ADFs – *Rhabditella axei* (DF5006), the species with the fewest serotonin-IR cells in the head. In several species, there was a second bilateral pair posterior to the putative ADF, sometimes more weakly-stained. More variable among the species is the number of additional moderately to weakly stained somas in the head that are also posterior to the nerve ring. We frequently observed one or two ventral apparently unpaired somas, one of which could be homologous to the *C. elegans* RIH neuron. As we have noted in *P. pacificus,* however, without seeing associated neurites, these identifications must be rather tentative.

### Monoaminergic function via VMAT / CAT-1 is required for P. pacificus egg-laying behavior

To identify behaviors requiring monoamine neurotransmitter function, we generated two mutant alleles in the *cat-1* gene, each with a premature stop codon in the first exon [*cat-1(csu115)* and *cat-1(csu116),* Suppl. Fig. S26A, S27]. We examined their phenotypes first to determine whether *cat-1*/VMAT shares an evolutionarily-conserved role in mediating nematode egg-laying behavior. In *C. elegans,* serotonin is required for normal egg-laying (Schafer 2005), and loss of serotonin in *P. pacificus tph-1* mutants also disrupts egg-laying (Okumura et al., 2017). Both *cat-1(csu115)* and *cat-1(csu116)* adult hermaphrodites showed a severe egg-laying defective phenotype (Egl), retaining excess unlaid eggs (>8); wild type adult hermaphrodites typically have 2 eggs in the uterus (Figure 12A, B). To confirm that the Egl defect in the *cat-1* mutants is indeed due to the lack of serotonin transport, we applied exogenous serotonin and rescued the slow egg-laying phenotype of *cat-1* mutants (Figure 12D), and to a lesser extent, the milder Egl defect of *bas-1* mutants which should be deficient in both serotonin and dopamine biosynthesis.

**Figure 12.**
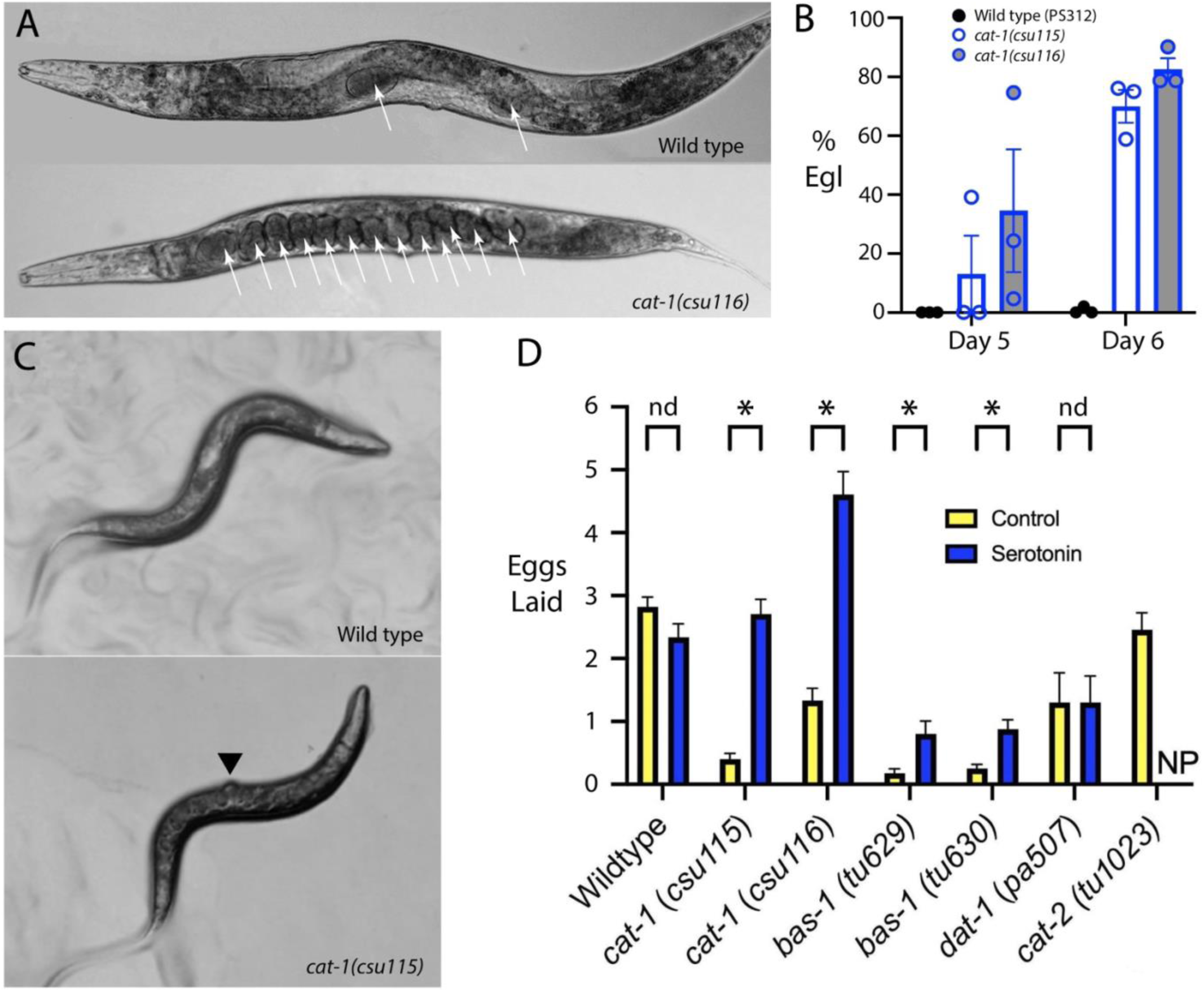
Egg laying is defective in *cat-1* mutants but is increased by exogenous serotonin. (A) *P. pacificus* wild type adult hermaphrodites typically retain 2 eggs; *cat-1* mutants accumulate many eggs (in both worms, white arrows indicate internal eggs). The *cat-1(csu116)* mutant shown has 13 internal eggs. (B) Percentage of adult hermaphrodites that are Egl (with ≥8 eggs) in wild type and two cat-1 alleles, in young adults (5 days after individual isolated as a newly laid egg), and 1 day later (6 days post egg isolation). Mean ± SEM, 3 tests each condition (n = 50-100 worms per test). (C) Many *cat-1* mutant adults have a protruding vulva (Pvl, black arrowhead), indicating abnormal vulval development, which may contribute to the Egl phenotype. (D) Number of eggs laid by individual *P. pacificus* adult hermaphrodites in liquid media, counted over two hours. Control is M9 buffer (yellow) vs. 10 mM serotonin in M9 (blue), mean ± SEM shown (n = 68-72 worms for each condition). Statistics: Multiple unpaired t-test; * indicates False-Discovery Rate for significant difference between control versus serotonin treated worms (P<0.001). “nd” denotes no discovery and “NP” indicates that the serotonin treatment for *Ppa-cat-2* was not performed due to the lack of an egg laying defect.

In many adult hermaphrodite *cat-1* mutant worms (both alleles), we also observed a protruding vulva (Pvl) phenotype (Figure 12C). Expression of *cat-1* transcripts in vulval epidermal cells suggests a possible role for the gene in normal vulval development. An abnormal vulva could also account for some of the Egl phenotype in *cat-1* mutants, although not so severe that it cannot be rescued by exogenous serotonin. Mutations in *cat-2* (dopamine biosynthesis) and *dat-1* (dopamine re-uptake, Suppl. Fig. S26B) that specifically affect dopamine availability did not affect the rate of egg-laying in this assay (Figure 12D). These results indicate that monoamine release, specifically serotonin, is crucial for regulating proper egg-laying behavior in *P. pacificus*.

### Monoamines are required for modulation of head movement during locomotion

We next tested whether monoamines are required for responses in other behaviors. In *C. elegans*, gentle anterior touch evokes an immediate reversal accompanied by a transient suppression of head oscillations/exploratory head movements that occur during forward locomotion (Chalfie et al., 1985; Alkema et al., 2005). In *tdc-1* and *cat-1* mutants, gentle touch reliably induces reversal, but suppression of head oscillations is defective, indicating that monoamines such as tyramine are involved in gentle touch–evoked suppression of head oscillations (Alkema et al., 2005). We examined whether this behavior is conserved in *P. pacificus*. In wild-type animals, reversal in response to gentle touch was accompanied by successful suppression of head oscillations in 69.8% of trials (37/53 animals). In contrast, *tdc-1(tu1007)* and *tdc-1(tu1009)* mutants showed suppression in only 39.1% (18/46, χ² test, p < 0.005) and 30.8% (16/52, χ² test, p < 0.001) of trials, respectively. Similarly, *cat-1(csu115)* and *cat-1(csu116)* mutants showed suppression in only 23.1% (12/52, χ² test, p < 0.001) and 24.5% (13/53, χ² test, p < 0.001) of trials, respectively. No significant differences were detected between alleles of the same gene or between *tdc-1* and *cat-1* mutants. These results indicate that, as in *C. elegans*, both *tdc-1* and *cat-1* functions are important for gentle touch–evoked suppression of head oscillations in *P. pacificus*.

### Monoamines are required for nictation behavior

We also tested whether monoamines are required for nictation, a nematode host-finding and dispersal-promoting behavior in which developmentally arrested dauer larvae, formed in crowding and starvation conditions, ‘stand’ on their tails and ‘wave’ their bodies in order to attach to passing animals to find new locations with food (Lee et al., 2012). The behavior is also observed in *P. pacificus* dauers when sand is added to plates (Brown et al., 2011). In our experiments, we observed that ∼46% of wildtype *P. pacificus* dauer larvae nictated by bracing themselves against grains of sand to stand on their tails and wave. We found that both *cat-1* mutants exhibited a strong reduction of nictation behavior, indicating that transport of monoamine(s) into vesicles is crucial for nictation (Figure 13A).

**Figure 13.**
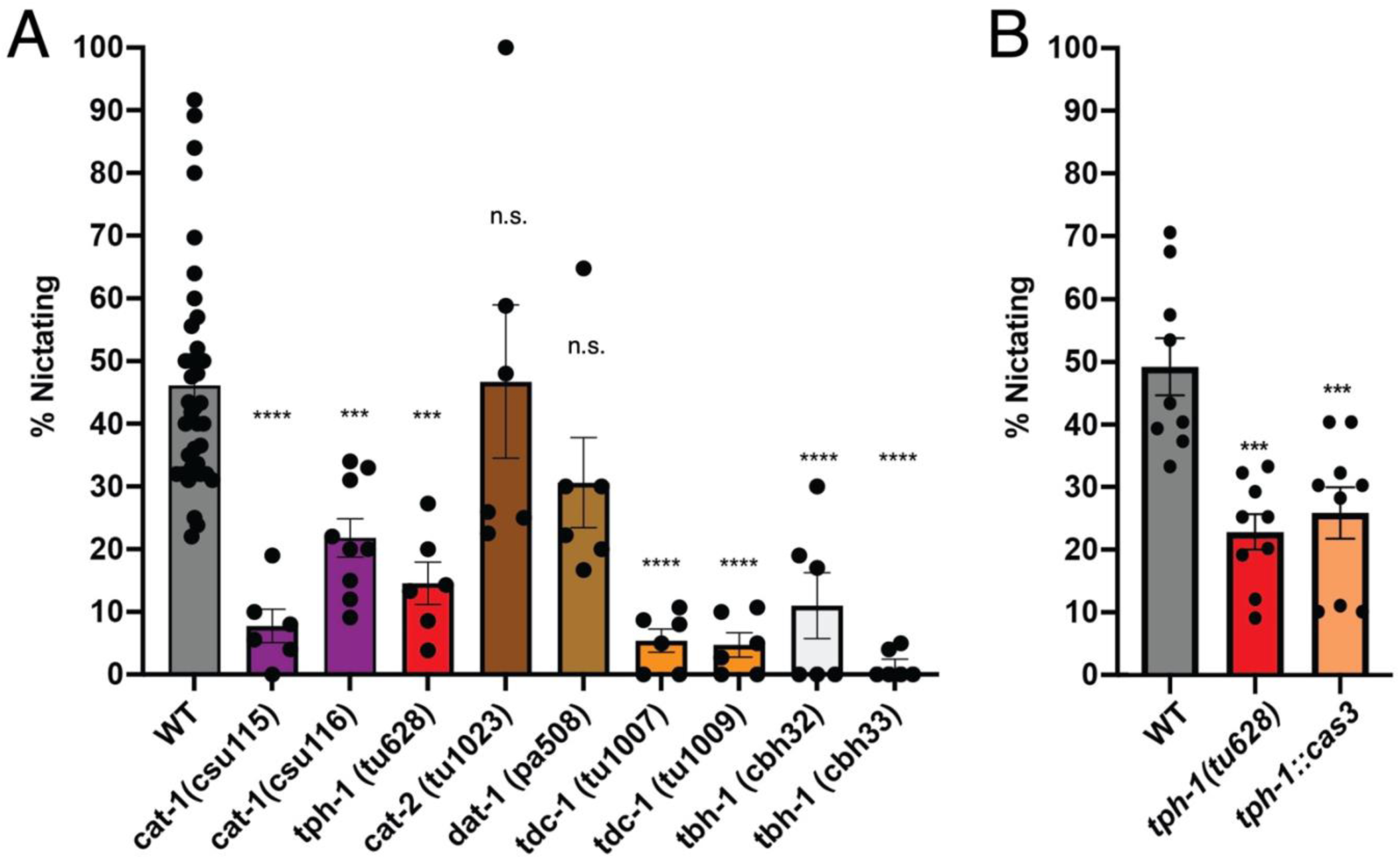
Multiple monoamines are required for nictation in *P. pacificus*. (A) The loss of *cat-1*/VMAT or genes for biosynthetic enzymes for serotonin (*tph-1*), tyramine (*tdc-1*), or octopamine (*tdc-1, tbh-1*), reduced the percentage of nictating dauer larvae. (B) Genetic ablation of the RIP and NSM neurons in *tph-1::cas-3* transgenic dauer larvae phenocopied the *tph-1(tu628)* loss of function nictation defect. Between 30-60 animals were used in each nictation assay and at least 6 assays were performed for each genotype. *P<0.05, ***P<0.001, ****P<0.0001 Dunnett’s multiple comparisons test show significant difference to wildtype (WT).

To determine which specific monoamines are required for nictation, we surveyed the responses of loss- and reduction-of-function mutants (some previously described and others generated and described first in this article) in genes encoding the rate-limiting step enzymes in the biosynthesis of these neurotransmitters (see Figure 1): *tph-1* for serotonin synthesis (Okumura et al., 2017); *cat-2* and *dat-1* for dopamine synthesis and reuptake, respectively; *tdc-1* for tyramine and octopamine synthesis; and *tbh-1* for octopamine synthesis. We observed a significant reduction in the percentage of nictating dauers in *tph-1* mutants (15% versus 46% in wildtype), comparable to the phenotype of the *cat-1* mutants, suggesting that serotonergic neurons contribute to nictation behavior (Figure 13A). Strikingly, we found that *tdc-1* and *tbh-1* mutants, reduced to 0-5% nictating worms, even more closely phenocopy the strongest effects seen in the *cat-1* mutants with a near complete loss of nictation (Figure 13A). These results show that at least two monoaminergic neurotransmitters are necessary for generating proper nictation behavior – serotonin and octopamine. Given the requirement of *tdc-1* for both tyramine and octopamine synthesis, it is difficult to rule out a role for tyramine as well.

In *C. elegans,* the inner labial IL2 sensory neurons are key regulators of nictation in the dauer larva (Lee et al., 2012) (Yang et al., 2020); these neurons are cholinergic (Pereira et al., 2015; Wang et al., 2024). To see whether *P. pacificus* IL2 neurons are also cholinergic, we examined expression of cholinergic markers using antibodies against *C. elegans* CHA-1 (ChAT, synthesis) and UNC-17 (VAChT, vesicular transport) proteins (Suppl Figure S27); we also examined anti-FLAG staining in a *P. pacificus cha- 1* C-terminal 2x-FLAG epitope tag insertion strain and observed the same pattern. The overall pattern of staining observed throughout the body with either method is quite similar to that seen in *C. elegans,* and the essential match between native locus epitope tag knock-in staining and *C. elegans* cholinergic antibody staining validates the use of these antibodies. (Detailed characterization of cholinergic staining in *P. pacificus* will be reported elsewhere). Although most individual somas can be difficult to identify (and most staining is found in neurites vs. cell bodies), we observed clear staining in the dorsal and ventral IL2 neurons in the head (IL2D & IL2V, Suppl Figure S29). These neuronal somas are displaced anteriorly (relative to their location in *C. elegans)* in front of anterior bulb of the pharynx and are the only neurons in this location (Cook et al., 2025). Although we also tested whether cholinergic function was necessary for nictation using reduction-of-function *cha-1* alleles, results were inconclusive because of the mutants’ pleiotropic phenotypes, including locomotion defects (Suppl Figures S30, S31).

In *C. elegans,* RIP neurons are the main synaptic outputs of IL2 neurons, required for nictation in dauer larvae (Lee et al., 2012) (Yang et al., 2020). RIP neurons in *P. pacificus* similarly receive strong inputs from IL2 neurons (Cook et al., 2025). Since *P. pacificus* RIP neurons express serotonin, to determine whether a subset of serotonergic neurons (those expressing the *tph-1p::RFP* transgene) are specifically required for nictation, we genetically ablated NSM and RIP neurons using the same *tph-1* promoter expressing human Caspase-3, as previously described (Okumura et al., 2017). Ablation of NSM and RIP phenocopied the *tph-1(tu628)* mutant nictation defect (Figure 13B), indicating a potential role for serotonin in nictation, possibly from serotonin released by RIP in response to IL2 activity.

## Discussion

Mechanisms underlying behavioral adaptations can vary at many levels including sensory perception, neuronal circuitry, and neuromodulation. Neuromodulation by classical neurotransmitters has been compared at deep taxonomic levels among model organisms such as *Drosophila melanogaster*, *Caenorhabditis elegans*, and the mouse (Flames and Hobert 2009) (Lloret-Fernández et al., 2018) (Aimon et al., 2023). The complete map of the *C. elegans* nervous system, including single-cell transcriptomes, synaptic and neurotransmitter connectomes, as well as the neuropeptide ‘wireless’ connectomes can now enable detailed comparisons to other nematode species at the resolution of individual neurons (Taylor et al., 2021) (Ripoll-Sánchez et al., 2023). *P. pacificus* behavior is driven by at least two forms of developmental plasticity− a genus-specific irreversible mouth-form dimorphism that enables interspecies predation and cannibalism (Lightfoot et al., 2021), and another phylum-specific and reversible type of plasticity on the dauer developmental decision that results in stage-specific host-finding behaviors (Casasa et al., 2020) (Carstensen et al., 2021) (Banerjee et al., 2023). To understand how such different behaviors evolve in nematodes, system level efforts have been made in *P. pacificus* to describe the pharyngeal and chemosensory neuron connectomes (Bumbarger et al., 2013) (Hong et al., 2019), and more recently, a complete head connectome (Cook et al., 2025). As a part of this ongoing effort, our comparative study addresses to need to identify aminergic neurons in *P. pacificus* to describe more precisely their differences in developmental patterning and behavior. This comprehensive cell identification is grounded in three complementary approaches: (1) endogenous mRNA expression of several key conserved neurotransmitter synthesis and transport genes; (2) transcriptional reporters that allow for the visualization of neurite morphology; (3) direct detection of monoamines by immunostaining. Our key findings follow and are summarized in Table 1.

### Monoaminergic neurons in P. pacificus: Serotonin

Expression of mRNAs from serotonin-related genes *tph-1, mod-5* and *cat-1* seen by HCR, and *cat-1p::GFP* and *tph-1p::GFP* reporter expression corroborate serotonin-IR staining patterns for I1, RIP, NSM, ADF and VC1-4. Three head neurons seen with anti-5HT staining (ADFL/R and an unpaired ventral midline cell, possibly RIH) did not express the *tph-1p::RFP* reporter; lack of expression presumably reflects missing regulatory elements in the 4.7 kbp upstream *tph-1p::RFP* promoter or a lack of genomic context in the extrachromosomal array. Although the three neuron types (I1, NSM, RIP) have also been observed in earlier immunostaining and transcriptional reporter studies, the RIPs were misidentified as the ADF amphid neurons (Wilecki et al., 2015) (Okumura et al., 2017). We did not observe 5HT staining in cells that did not express *tph-1* transcripts or the *cat-1p::GFP* reporter, which might suggest serotonin production in a parallel, phenylalanine hydroxylase/PAH-1-dependent pathway (Yu et al., 2023). We found that most or all serotonin-IR cells in *P. pacificus* synthesize serotonin endogenously, as opposed to relying on serotonin uptake via the serotonin transporter MOD-5/SERT. This contrasts with *C. elegans*, in which some neurons, such as RIH and AIM, depend on serotonin reuptake (Jafari et al., 2011). In these experiments in *P. pacificus,* however, we did not reliably observe serotonin-IR or marker gene expression in a ventral midline unpaired neuron (possibly RIH) that has been seen previously (Rivard et al., 2010); this cell may be an uptake-dependent cell.

### Monoaminergic neurons in P. pacificus: Dopamine

The previously identified dopamine neurons CEP, ADE and PDE express transcripts from the *cat-2/TH* and *cat-1/VMAT* genes, as expected, and we find no additional dopaminergic neurons. This set of cells is the same as seen in *C. elegans* and appears to be highly conserved in rhabditid nematodes (Rivard et al., 2010). Although vulval region uv1 cells may use dopamine (based on 5HTP conversion to 5HT), they do not express either *cat-2/TH* or *cat-1/VMAT* genes. Since this 5HTP-induced serotonin-IR in the uv1 cells is dependent on the *bas-1/*AAADC gene, it is possible that they use a different, unknown monoamine, such as tryptamine. Lack of *cat-1/VMAT* expression in ‘neuroendocrine’ uv1 cells does not preclude function: in *C. elegans,* the uv1s inhibit egg laying via tyramine release, but lack *cat-1/VMAT* expression like in *P. pacificus* (Alkema et al., 2005).

### Monoaminergic neurons in P. pacificus: Tyramine & Octopamine

As with the case for dopaminergic neurons, TA and OA neurons RIM and RIC appear to be the same as found in *C. elegans,* expressing the expected biosynthetic enzyme genes *tdc-1* (TA, OA) and *tbh-1* (OA). In *C. elegans*, octopamine biosynthesis requires tyrosine decarboxylase (TDC) to convert tyrosine into tyramine, and tyramine β-hydroxylase (TBH) to convert tyramine into octopamine (Alkema et al., 2005) (Flames and Hobert 2011) (Yu et al., 2023). Although the two biosynthetic pathways overlap, immunostaining with TDC-1 and TBH-1 shows tyraminergic neurons are distinct from octopaminergic neurons, and that tyramine is not just a precursor for octopamine biosynthesis. *Tdc-1* is expressed in the *C. elegans* RIM and nearby RIC neurons, as well as the neuroendocrine vulval uv1 cells and gonadal sheath cells, whereas *tbh-1* is primarily expressed in the RIC neurons and the gonadal sheath cells (Alkema et al., 2005). In *P. pacificus,* the *cat-1p::GFP* reporter also showed expression in two pairs of neurons with strong resemblance to RIM and RIC neurons in their neurite trajectory and cell body positions. It is important to note, however, that two other neurons share similar cell body positions and neurite projections, such as the RIB and AIZ neurons (Serrano-Saiz et al., 2013) (Cook et al., 2025), but such interpretation would not be parsimonious as it would require the simultaneous loss of *cat-1* expression in RIM or RIC, with a concomitant gain of *cat-1* expression in RIB or AIZ.

### Other possible roles of CAT-1/VMAT

In the head, we observed expression of *cat-1* transcripts and the *cat-1p::GFP* reporter in at least two pairs of additional cells that do not express markers for serotonin, dopamine, tyramine or octopamine. Therefore, these cells may use an unknown monoamine or the small molecule betaine, recently found to function as a neurotransmitter. In *C. elegans,* CAT-1/VMAT has been shown able to transport the ‘non-classical’ small molecule betaine, more commonly known as an osmolyte, into synaptic vesicles (Peden et al., 2013;Hardege et al., 2022). Any of the cells expressing *cat-1* in *P. pacificus* could also use betaine as a transmitter or co-transmitter. In *C. elegans,* betaine functions as a co-transmitter in the tyraminergic RIM neuron, helping to regulate foraging behavior (Hardege et al., 2022).

### Possible non-neuronal use of monoamines

As also seen in *C. elegans, P. pacificus tbh-1* expression was observed in gonadal sheath cells. In the absence of co-expression of *tdc-1* (required for octopamine synthesis) or *cat-1* (required for vesicle loading), however, the functional significance of such expression is unclear. The same can be said for the lack of co-expression of *tdc-1* and *cat-1* in separate, adjacent vulval epidermal cells surrounding the vulval pore, although the vulval defects we observed *in cat-1* mutants suggest a functional role for *cat-1* expression. It is possible that these Along with the potential uv1 homologs, this does suggest there are potentially even more ‘neuro-endocrine’ cells at the vulva in *P. pacificus* than in *C. elegans*.

### Serotonin signaling in egg-laying

Our results show that vesicular monoamine transport via CAT-1/VMAT is essential for egg-laying behavior in *P. pacificus*, similar to the role of serotonin in *C. elegans*. Loss-of-function mutations in *cat-1* resulted in severe egg-laying defects, which could be rescued by exogenous serotonin, confirming a key role for serotonin signaling in *P. pacificus*. In *C. elegans* hermaphrodites, the egg-laying neuromuscular circuit includes a bilateral pair of Hermaphrodite-specific neurons (HSNs) that are both cholinergic and serotonergic, and six cholinergic Ventral ‘C-type’ motor neurons (VC1-6). The two VCs closest to the vulva (VC4, VC5) take up serotonin and are sometimes weakly serotonin immunoreactive; they subtly influence egg-laying by serotonin release (Duerr et al., 2008) (Duerr et al., 1999) (Pereira et al., 2015). The strongly serotonin-IR HSNs, in contrast, are essential for normal egg-laying; loss of the cells causes a severe Egl phenotype (Schafer 2005). Since there is no serotonin-IR HSN in *P. pacificus,* and four VC neurons (VC1-4) flanking the vulva are strongly serotonin-IR (Loer and Rivard, 2007); it is likely that these VC neurons are the primary regulators of egg-laying in *P. pacificus,* in contrast to what is seen *C. elegans*.

### Monoaminergic neurons in nictation behavior

While a role in egg-laying for VC neurons appears to be conserved in *P. pacificus* and *C. elegans*, serotonin in I1 and RIP neurons in *P. pacificus* appears to represent the ancestral state in nematode species outside of the *Caenorhabditis* genus. Notably in *P. pacificus*, serotonin-signaling appears to be an important component of the dauer-specific nictation behavior, since *tph-1* mutants and *tph-1::cas-3* genetic ablation of NSM and RIP resulted in nictation defects. Given that the IL2 to RIP synaptic connection is conserved in *P. pacificus* (Cook et al., 2025), the lack of a nictation phenotype in *C. elegans tph-1* mutants (Lee et al., 2012) and the difference in RIP neurotransmitter are consistent with an interpretation that RIP neurons are responsible for facilitating nictation behavior in *P. pacificus*.

Interestingly, the RIP neuron is a ‘rich club’ highly interconnected neuron in *P. pacificus* but is not so in *C. elegans* (Cook et al., 2025). Information flow via RIP neurons, the only neurons that synaptically connect the somatic and pharyngeal (enteric) nervous systems in both *P. pacificus* and *C. elegans,* is substantially different. Whereas in *C. elegans,* the nerve ring neurites of RIP only receive inputs, in *P. pacificus,* the RIP neurons’ more extensive nerve ring neurites provide synaptic outputs on to several different neurons (Cook et al., 2025).

Serotonin has been shown to play a key role in predatory feeding behavior in *P. pacificus* as serotonergic neurons are critical for controlling predation-related motor patterns. These include the precise coordination of the tooth movement with pharyngeal pumping that is necessary for killing behavior. Consistent with this, mutants defective in serotonin synthesis also exhibit severe impairments in their ability to kill prey efficiently (Okumura et al., 2017) (Ishita et al., 2021). More recently, both octopamine and tyramine have also been implicated in regulating predatory behaviors in *P. pacificus*.

Octopamine promotes aggression, while tyramine instead biases animals toward more docile behaviors. Together, they establish an antagonistic framework for balancing these behavioral drives (Eren et al., 2024). Interestingly, in *C. elegans* there is little evidence for such antagonism. Tyramine is instead linked to escape behavior while octopamine regulates fasting (Pirri et al., 2009;Churgin et al., 2017). Despite this, the RIC octopaminergic and RIM tyraminergic neurons themselves appear to be conserved across both species. Taken together, our findings suggest that *P. pacificus* may have co-opted or modified the outputs of the modulatory circuits, including serotonergic RIP neurons to support the evolution of predatory behaviors. In this view, serotonergic RIP neurons may have provided a pre-adapted substrate, upon which novel behavioral functions could evolve.

In addition to serotonin signaling, other aminergic neurons appear to also collaborate in the execution of nictation in *P. pacificus.* Since *P. pacificus cha-1* expression (choline acetyltransferase) is conserved in IL2 neurons, it is possible that we have uncovered an evolutionary shift in downstream targets of the *P. pacificus* nictation circuit consisting of the mechanosensory IL2 neurons with input to the interneurons RIP, RIM, and RIC. In contrast, compared to other developmental stages, the *C. elegans* IL2 neurons form their strongest connection to the RIG interneurons (glutaminergic), which are required for nictation (Yim et al., 2024). In *P. pacificus,* dauer regulation pathways have diverged significantly from those found in *C. elegans.* None of the mutations in each of the seven *P. pacificus daf-7* paralogs resulted in the dauer formation constitutive phenotype (Daf-c) (Lo et al., 2023), and *tph-1* mutants do not have a Daf-c phenotype unlike their *C. elegans* counterpart (Sze et al., 2000), so it is likely that the specific neurons involved in the remodeling of the nervous system during dauer development may also have diverged between these two nematode species.

### Evolutionary origins of differences in serotonin-IR neurons

Our examination (and re-examination) of anti-serotonin staining in other nematodes of known relationship to *P. pacificus* confirms the highly conserved neurotransmitter expression in the pharyngeal NSM neuron and sensory ADF neuron. Furthermore, our observations indicate that expression of serotonin in the pharyngeal I1 and anterior ganglion RIP neurons is ancestral and that loss of serotonin in these neurons in the genus *Caenorhabditis* is derived. Only a few species that we examined lack serotonin-IR RIP neurons; the I1 neurons, however, appear to have lost serotonin expression independently in at least three other groups. It is interesting to note that although no reporters in *C. elegans* show expression of serotonin marker genes in I1 (Wang et al., 2024), scRNA data indicate a low level of *tph-1* transcript expression (Taylor et al., 2021), perhaps a vestigial feature. Bolstering a determination of serotonin in I1 and RIPs as being ancestral, our examination includes outgroups to *Pristionchus* and *Caenorhabditis* (clade V or 9), including newly examined cephalobs (*Propanagrolaimus* and *Acrobeloides*) and plectids (*Plectus*). We note that *Plectus* (*P. aquatilis* and *P. acuminatus)* has been examined previously for serotonin-IR neurons, showing beautifully stained NSMs, ADFs, and an anterior pharyngeal neuron identified as possibly I2 (Henne et al., 2017); this cell is almost certainly I1. These authors sometimes observed a second dorso-lateral pair posterior to ADF, which we also saw frequently, but did not describe the 2 - 6 anterior ganglion neurons including cells that could include RIPs. We speculate that in some species we examined, serotonin-IR neurons in the anterior ganglion (or further anterior) may be IL2 homologs. Although we have not observed serotonin-IR in IL2 neurons in *P. pacificus,* we did sometimes observe weak expression of *mod-5* transcripts in IL2s. Expression of the *mod-5/SERT* gene is all that would be required for a cell to become serotonin-IR. Becoming a serotonin ‘absorbing’ cell could be a stepping-stone to becoming a functional serotonergic neuron; after acquiring uptake, expression of *cat-1/VMAT* would suffice.

## Acknowledgements

We thank J. Cardenas and I. Dimov for technical assistance. We thank Chi Chen for generating nematode strains. Some strains were provided by the CGC, which is funded by NIH Office of Research Infrastructure Programs (P40 OD010440). We thank Philipp Schiffer for sharing *Acrobeloides* and *Plectus* nematode strains. We thank Kevin Collins for sharing information about uv1 cells in *C. elegans.* We thank David Rudel for examining and commenting on images of *tbh-1* staining in gonads.

## Funding

This research was funded by NIH SC1GM140970 to RLH, NIH BUILD PODER UL1 GM1 18976 to MFH, and T34GM136450 to DVB. An endowment from the Fletcher Jones Foundation and NSF MRI award 1229443 (LSCM acquisition) supported CML. This work was funded by the Howard Hughes Medical Institute (OH).

## Author contributions

Conceptualization: CML, RH, OH

Methodology: CML, HY, LG, YR, RH

Investigation: CML, HY, LG, YR, MFH, DVB, HRC, JM, LR, TM, SJC, MO, JL, RH

Visualization: CML, HY, LG, LR, SJC, RH

Funding acquisition: CML, OH, RH

Project administration: CML, RH, OH

Supervision: CML, RH, OH

Writing – original draft: CML, RH

Writing – review & editing: CML, HY, MO, JL, RH, OH

## Competing interests

Authors declare that they have no competing interests.

## Supplemental Materials and Methods

*dat-1* and *cha-1* alleles: These indel alleles were generated during screens intended to epitope-tag the genes. Injection mixes were prepared using the method described in (Dokshin et al., 2018) for *C. elegans,* with the addition of lipofectamine (Adams et al., 2019): Cas9 - 0.5 μl at 10 μg/μl, tracrRNA – 5 μl at 0.4 μg/μl, 2.8 μl at 100 µM. This mixture was incubated for 10 min at 37°C. Then 0.6 µl Lipofectamine RNAiMax reagent (final conc 3%, Thermofisher 1377803) and 1 µl single stranded repair template at 100 µM were added, plus 12.1 µl nuclease free water to an injection mix final volume of 20 µl. For the *cha-1* screen, 16 young adult hermaphrodites (worms with ≤2 eggs) were injected and placed on individual plates; these parents were removed after ∼24 hrs of laying eggs. Three days later, ≤ 20 F1s were singled from each parental plate (some P0’s had fewer than 20 progeny). Two or more days later, after the adult had many progeny eggs and larvae, screening by PCR was performed using a modified ‘single worm’ PCR method (Williams et al., 1992). The F1 adult plus several larvae and eggs were picked into 2 µl PCR / Proteinase K lysis buffer (0.5 mg/ml proteinase K final concentration, prepared fresh) in the lid of a PCR tube. Lysis buffer with worms was spun down, frozen in liquid nitrogen, then immediately thawed in a water bath at 37-65°C, then repeated; the rapid freeze-thaw cycle was performed a total of 3 times. PCR tubes were put in a Thermocycler, digested at 65 C for 60 min, heat-inactivated at 95 C for 15 min, then cooled to 4°C, and within ∼1 hr, PCR mix with appropriate primers was added, and PCR performed with primers flanking the expected cut & insertion site. Clone plates yielding PCR bands with increased sizes consistent with repair template insertion were selected for further subcloning to isolate homozygous mutants. Among the homozygotes isolated were indel mutants. For the *cha-1* screen, 271 F1 clones were screened, yielding one proper 2x FLAG epitope C-terminal insertion and 3 indel mutants. One mutant was apparently homozygous larval lethal, with heterozygotes throwing ∼ ¼ early larval arrest coilers (a phenotype like seen in *C. elegans cha-1* null homozygotes); this strain was eventually lost. For the *dat-1* screen, 15 worms were injected; parents removed after ∼48 hrs. Because of a need to delay F1 singling, those plates were then put at 15°C for 4 days, at which time ∼ 270 F1s were singled and subsequently grown at standard temperature. 158 clones were screened by PCR; from an initial screen of 94, ∼17% appeared to have a genomic alteration (from one ‘jackpot’ P_0_ plate, 50% of F1’s showed genomic alteration). Therefore, the remainder of clones were not followed up. Subcloning and rescreening resulted in the isolation of 5 independent strains with the proper N-terminal HA tag insertion, 1 strain with an indel mutation and partial tag insertion, and 3 strains with indel mutations, all resulting in likely nulls.

## Abbreviations

5HT, TA, OA, DA, Ach

**To be submitted to eLife**

## Supplemental Figures

**Suppl Fig S1.**
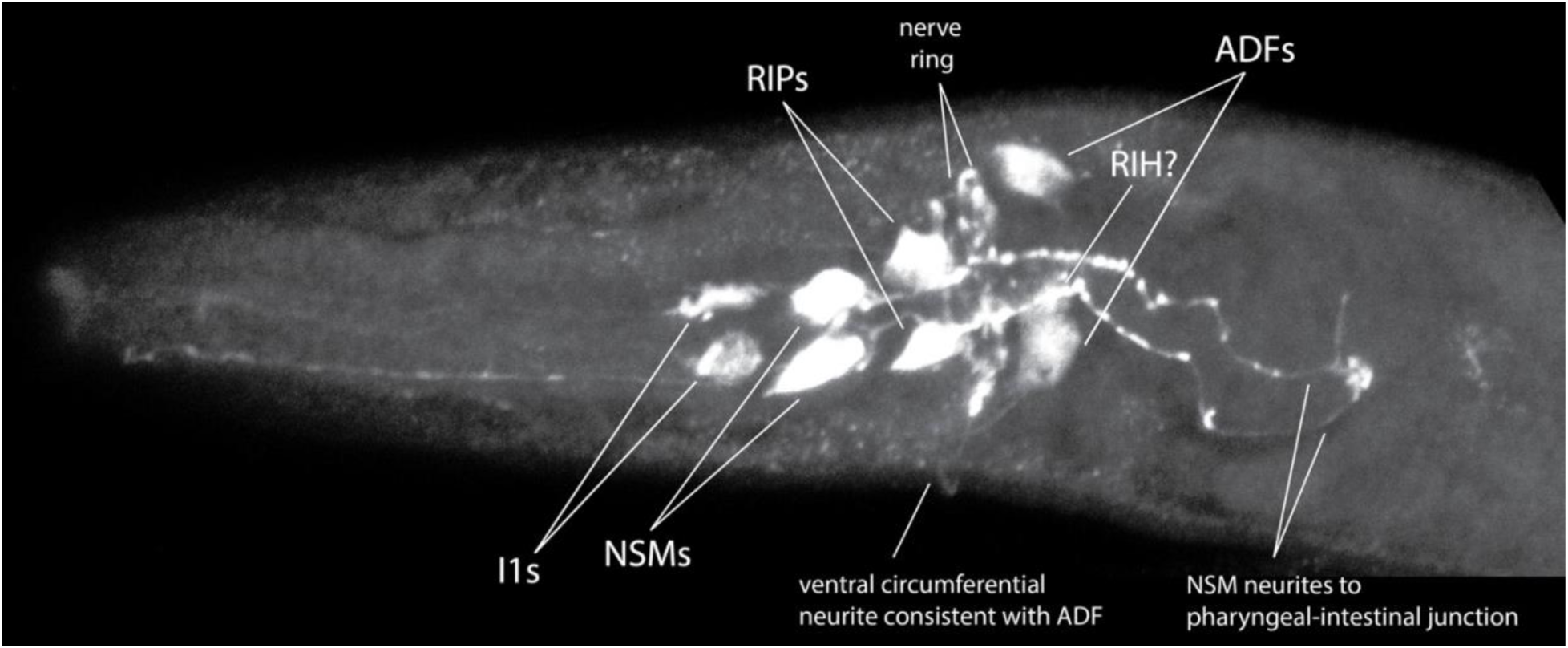
Anti-5HT staining of wildtype *P. pacificus*, larval head showing all serotonin-IR head neurons as in Figure 1 adult, Max IP, anterior to the left, ventral view. In the pharynx: I1s, NSMs; anterior ganglion in front of the nerve ring: RIPs; posterior to the nerve ring: ADFs and unpaired possible RIH. Because the pharynx is not kinked as it Figure 5, the unpaired neuron is shown clearly to be posterior to the nerve ring.

**Suppl Fig S2.**
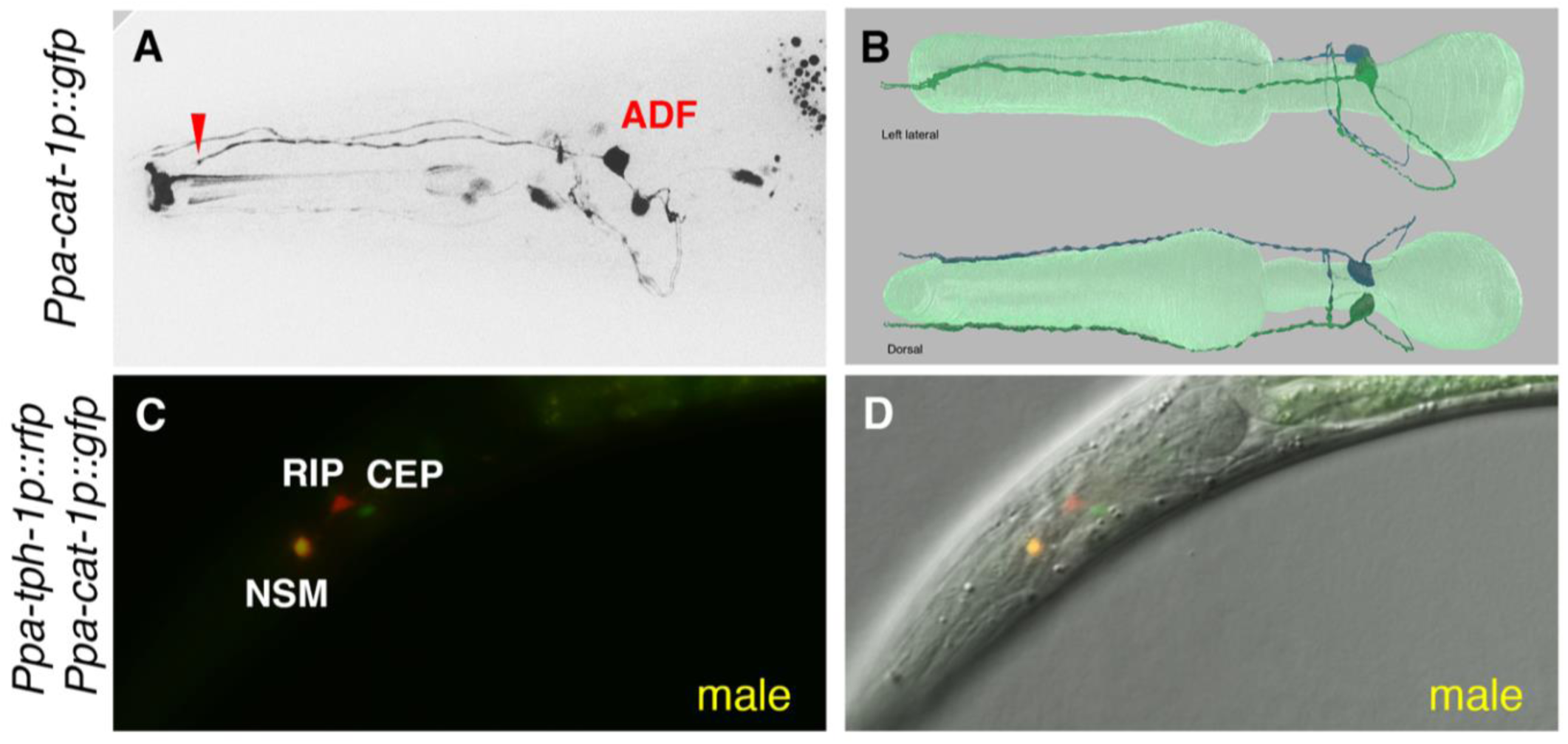
Additional reporter expression in aminergic neurons in *P. pacificus.* (A) *cat-1p::gfp* expression in the amphid neuron homolog ADF(AM9) with double ciliated ending (arrowhead). Anterior is to the left. (B) 3D-rendering of the ADF neuron pair in lateral and dorsal orientations. (C-D) F_1_ hermaphrodite adult showing overlap of *cat-1p::gfp* and *tph-1p::rfp* reporter expression in the NSM neuron.

**Suppl Fig S3.**
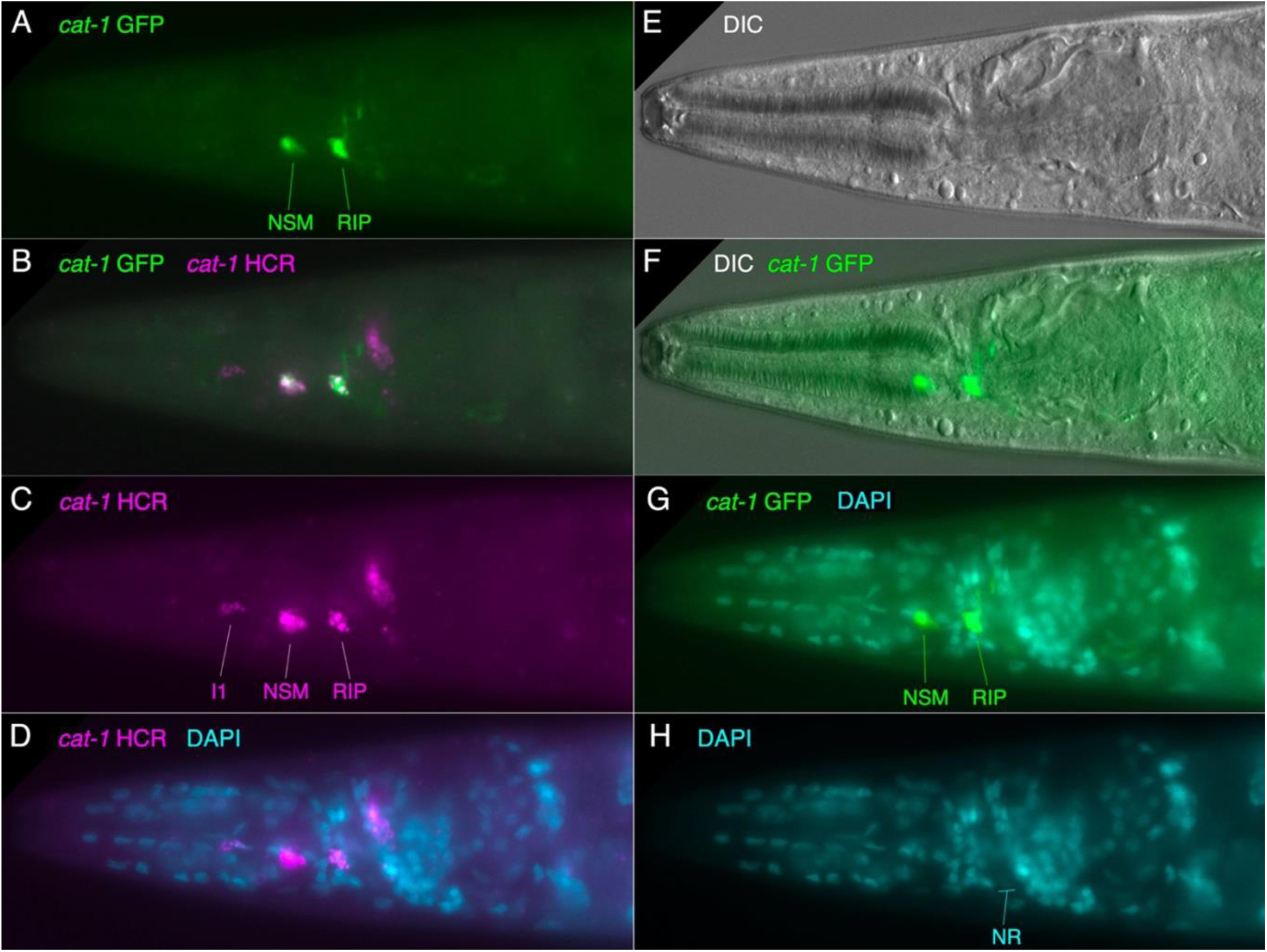
Expression of *cat-1* transcripts via HCR in head in *cat-1*::GFP strain showing RIP neuron in anterior ganglion with neurite to dorsal nerve ring. Anterior to the left, ventro-lateral view of head. All images are the same focal plane, except (E), slightly different to show pharynx outline better. (A) Mosaic expression of *cat-1::GFP* reporter in NSM and RIP on one side of the head. Note the characteristic morphology of RIP neurites. (B) Colocalization of *cat-1* transcripts with *cat-1::GFP* in NSM and RIP. (C) *cat-1* transcripts (magenta). (D) *cat-1* transcripts and DAPI showing RIP in anterior ganglion anterior to nerve ring (nucleus-free region, marked in H). (E) DIC alone (F) DIC with *cat-1::GFP*. (G) *cat-1::GFP* with DAPI showing RIP ending in dorsal nerve ring. (H) DAPI showing nuclei and location of nerve ring (NR).

**Suppl Fig S4.**
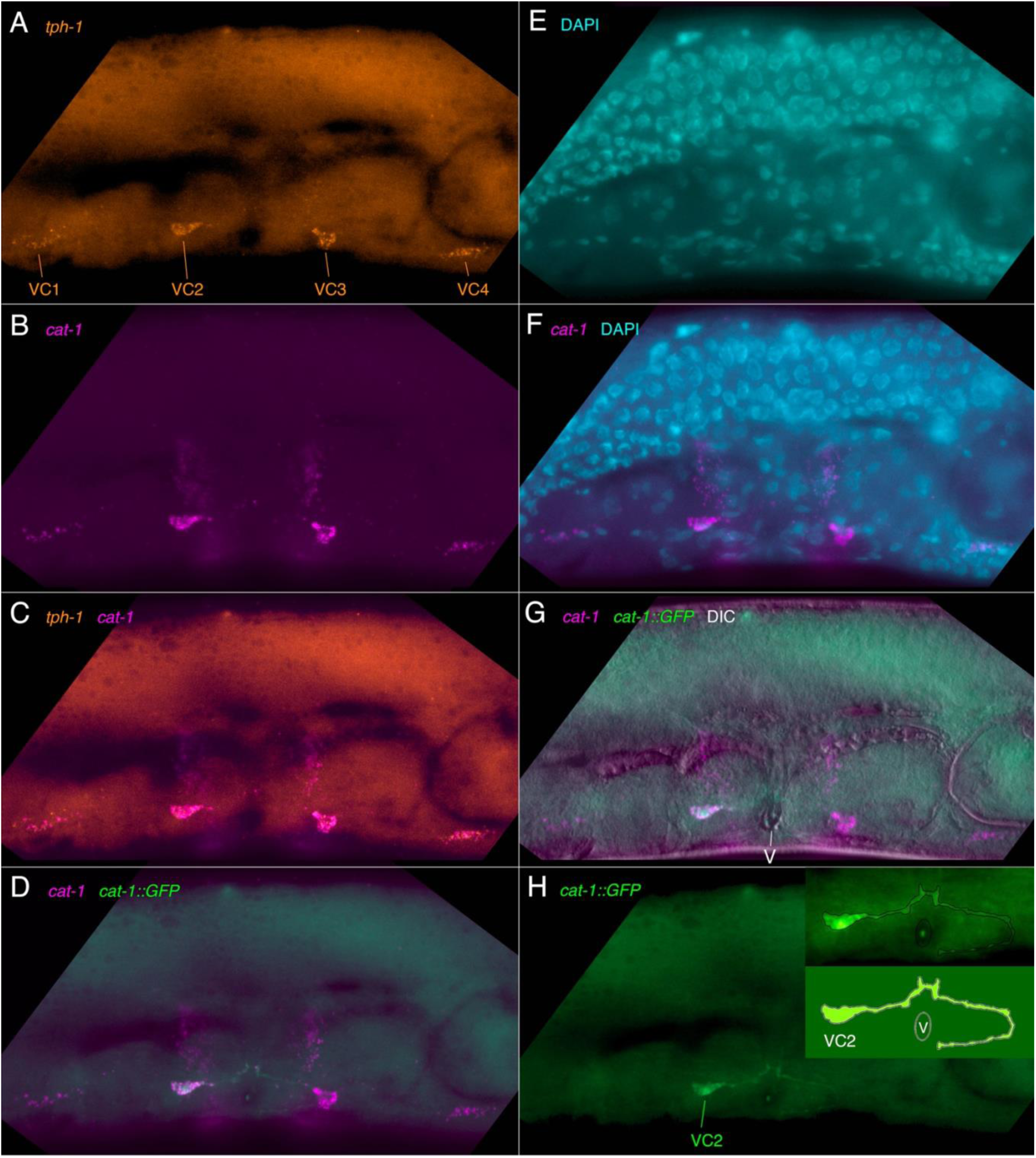
Coexpression of *tph-1* and *cat-1* transcripts in vulval region in *cat-1*::GFP strain. Anterior to the left, ventral down, all images of the same focal plane. (A) *tph-1* transcripts expressed in VC1-4 in the VNC (B) *cat-1* transcripts expressed in VC1-4 and in vulval cells in ventrolateral body wall. (C) *tph-1* and *cat-1* transcripts colocalized in VC1-4. (D) *cat-1* transcripts colocalized with *cat-1*::GFP reporter fluorescence of a single labeled VC neuron, VC2. (E) DAPI staining of vulval region, showing compact nuclei of VNC. (F) *cat-1* transcripts colocalized with DAPI-stained VNC nuclei. (G) Including DIC shows the location of vulval pore between VC2 & VC3 (marked with ‘V’). (H) *cat-1*::GFP alone, showing VC2 neuron. Interestingly, the neurite crosses to the other side of the vulval pore and grows back anteriorly with varicosities. Inset: schematic outline of VC2 major branches. (Note that fine branches may not be seen.)

**Suppl Fig S5.**
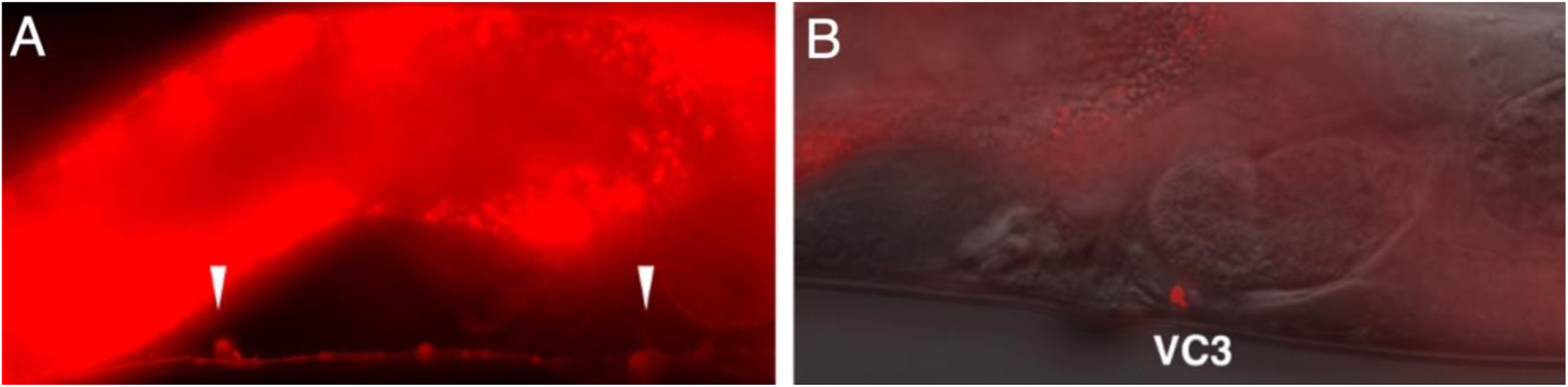
Cells expressing the *tph-1p::rfp* reporter in the mid-body of *P. pacificus*. (A, B) VC neurons in the VNC express the *tph-1p::rfp* reporter. (A) Arrowheads indicate somas in ventral nerve cord near the vulva; neurites are also apparent. (B) VC3 neuron soma just posterior to the vulva, including DIC.

**Suppl Fig S6.**
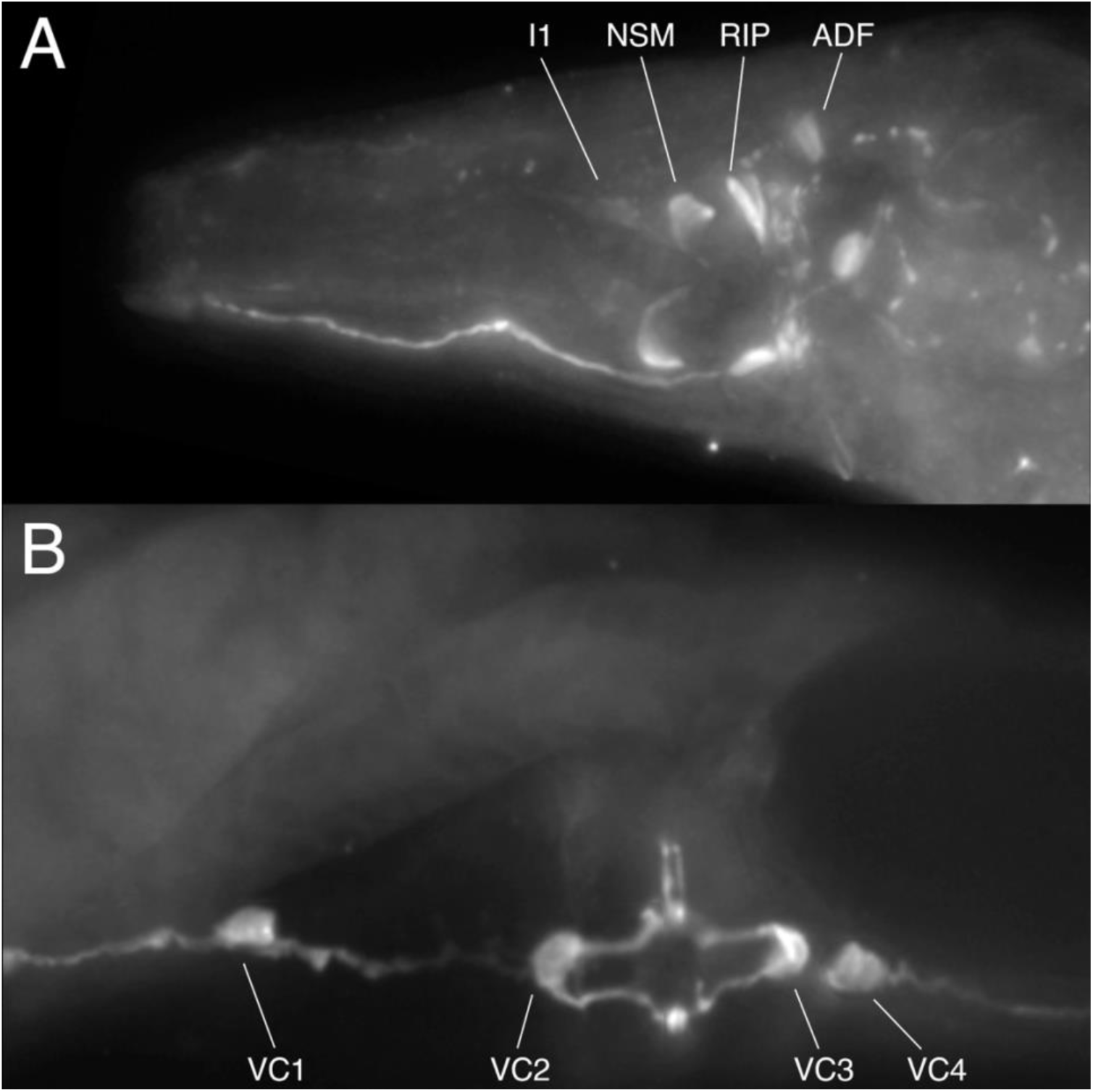
Serotonin-IR is unchanged in *mod-5* mutants. Anti-serotonin staining in *mod-5(tu587)* mutant. Anterior to the left. The same result was seen with the other allele, *mod-5(tu586)*. (A) Adult hermaphrodite head, MaxIP of several focal planes, approximately dorsal-ventral view. All serotonin-IR cells (I1s, NSMs, RIPs, and ADFs) are seen, as in wildtype. Staining of the unpaired ventral serotonin-IR neuron, which is weak and unreliable, was not seen in wildtype worms in these experiments, so this cell might still be uptake-dependent. (B) Adult midbody vulval region, ventro-lateral view. VC neurons stain as in wildtype. We observed the same results for worms treated with SERT blockers fluoxetine or imipramine at various concentrations; no cells were affected except at the highest concentration used, which eliminated or greatly reduced serotonin staining uniformly in all cells, a likely non-specific effect since these drugs can bind other targets beside SERT.

**Suppl Fig S7.**
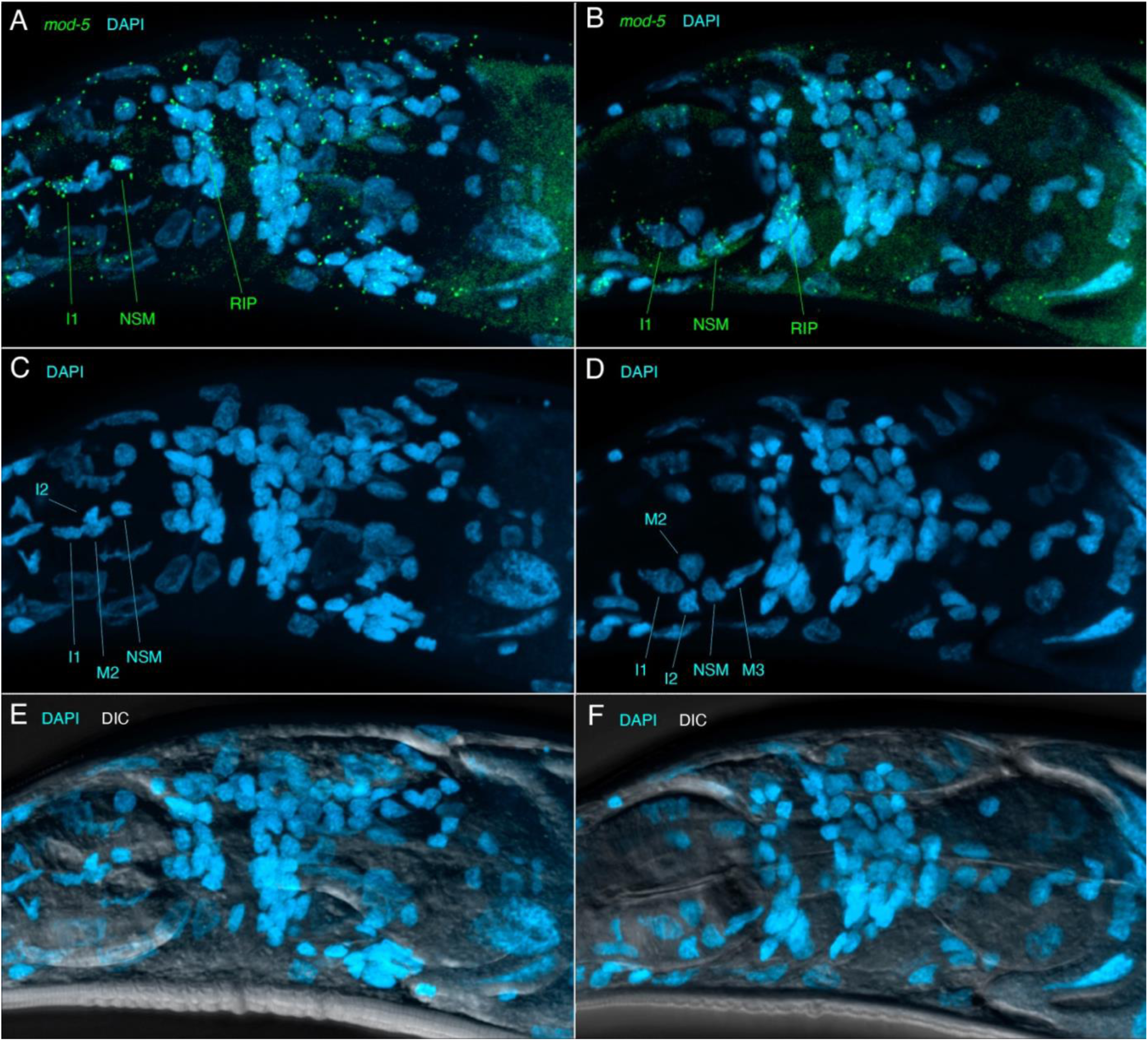
Serotonergic neurons in the head express *mod-5/SERT* transcripts. Anterior to the left, ventrolateral views with ventral approximately down. MaxIPs of a few focal planes on left (A,C,E) and right sides (B, D, F) of head. (A) Expression of *mod-5* transcripts (green) in identified pharyngeal (I1, NSM) and anterior ganglion (RIP) serotonin neurons, left side, with DAPI (cyan). (B) Same on the right side. (C) DAPI alone, same as A, with other adjacent DAPI-stained pharyngeal neurons identified. (D) Same on the right side. (E) Including DIC to show outlines of pharyngeal bulbs. (F) Same on the right side.

**Suppl Fig S8.**
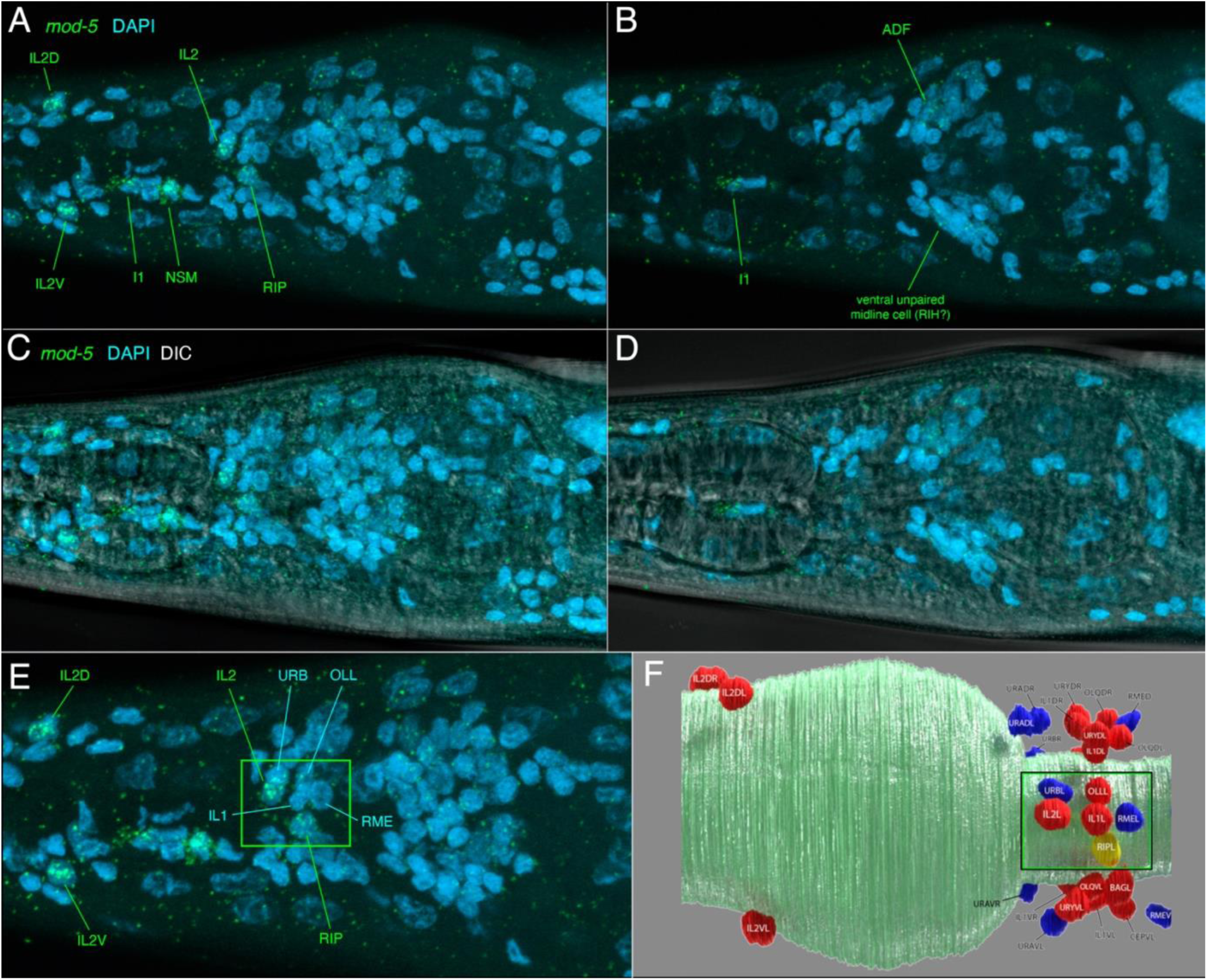
Head serotonergic and other neurons in the head sometimes express *mod-5/SERT*. Anterior to the left, ventral down, all images. MaxIPs of the same few focal planes (A, C, E), or a nearby set of focal planes (B, D). (A) Expression of *mod-5* transcripts (green) in identified pharyngeal (I1, NSM) and anterior ganglion (RIP) serotonin neurons, plus IL2 neurons (D, V, lateral), with DAPI (cyan). (B) Nearby focal planes showing *mod-5* transcripts in serotonin neurons posterior to the nerve ring, ADF and possibly RIH. (C, D) Including DIC to show outlines of pharyngeal bulbs. (E) Closeup of anterior ganglion, lateral neurons (green box). Several adjacent DAPI-stained neuronal nuclei are identified by position. (F) Map of identified anterior ganglion neuronal nuclei, left side lateral view (boxed as in E), 3D rendering from EM reconstruction (Cook et al., 2025).

**Suppl Fig S9.**
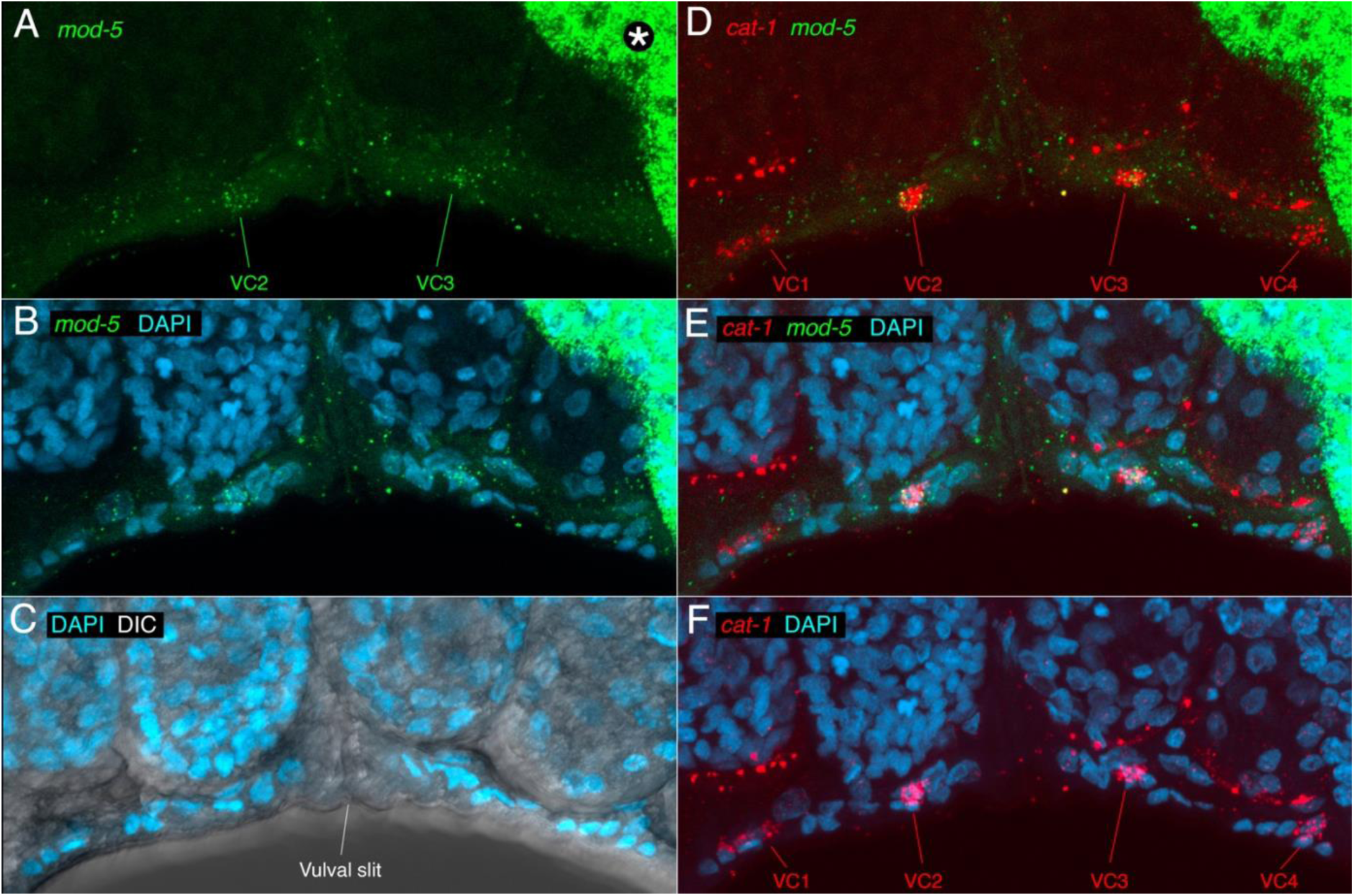
Vulval-proximal VC neurons express *mod-5/SERT*. Closeup of vulval region; anterior to the left, ventral down. Images are MaxIPs of several focal planes. (A) *mod-5* transcripts expressed in VC2 and VC3 (proximal VCs) in the VNC, but not obviously in distal VCs (VC1, VC4). Asterisk indicates intestine, which displays high background and autofluorescence in this preparation. (B) *mod-5* transcripts are associated with compact VNC nuclei shown with DAPI staining. (C) DIC reveals location of vulval opening. Four embryos in the uterus are seen above the VNC. (D) *cat-1* transcripts are expressed in VC1-4, but colocalized with *mod-5* only in proximal VCs. (E) DAPI staining of vulval region, showing compact nuclei of VNC. (F) *cat-1* transcripts colocalized with DAPI-stained VNC nuclei.

**Suppl Fig S10.**
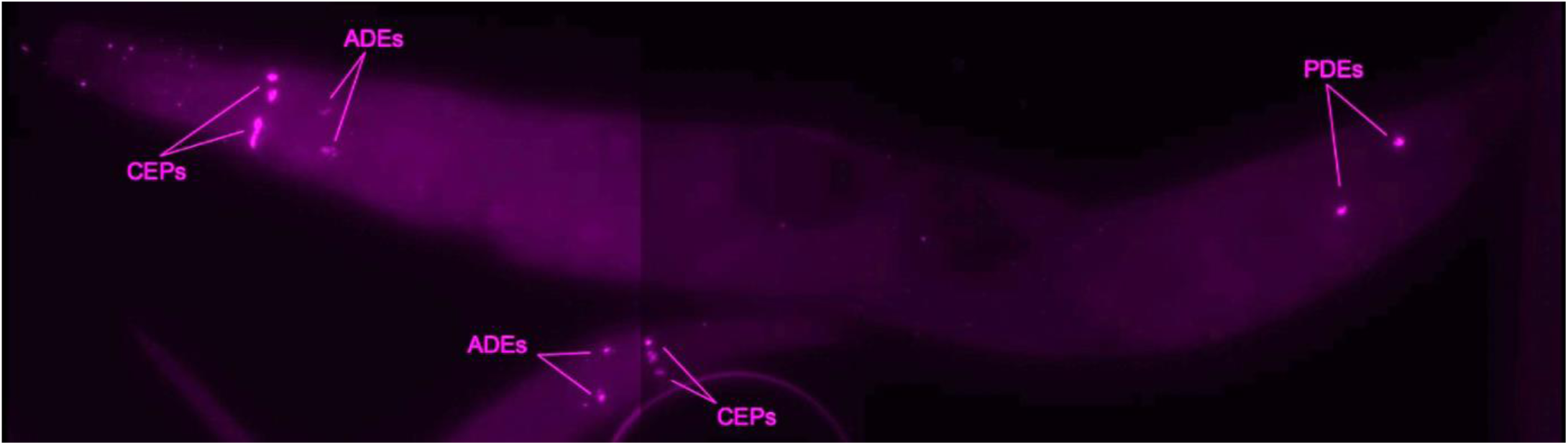
Expression of cat-2 transcripts in identified dopaminergic neurons. Whole adult (anterior to the left) and larval head (anterior to right) expression of *cat-2* transcripts (magenta), MaxIP. Head neurons CEPs and ADEs are seen both in the adult head (top) and larval head (below). The posterior body shows PDE neurons on left and right sides.

**Suppl Fig S11.**
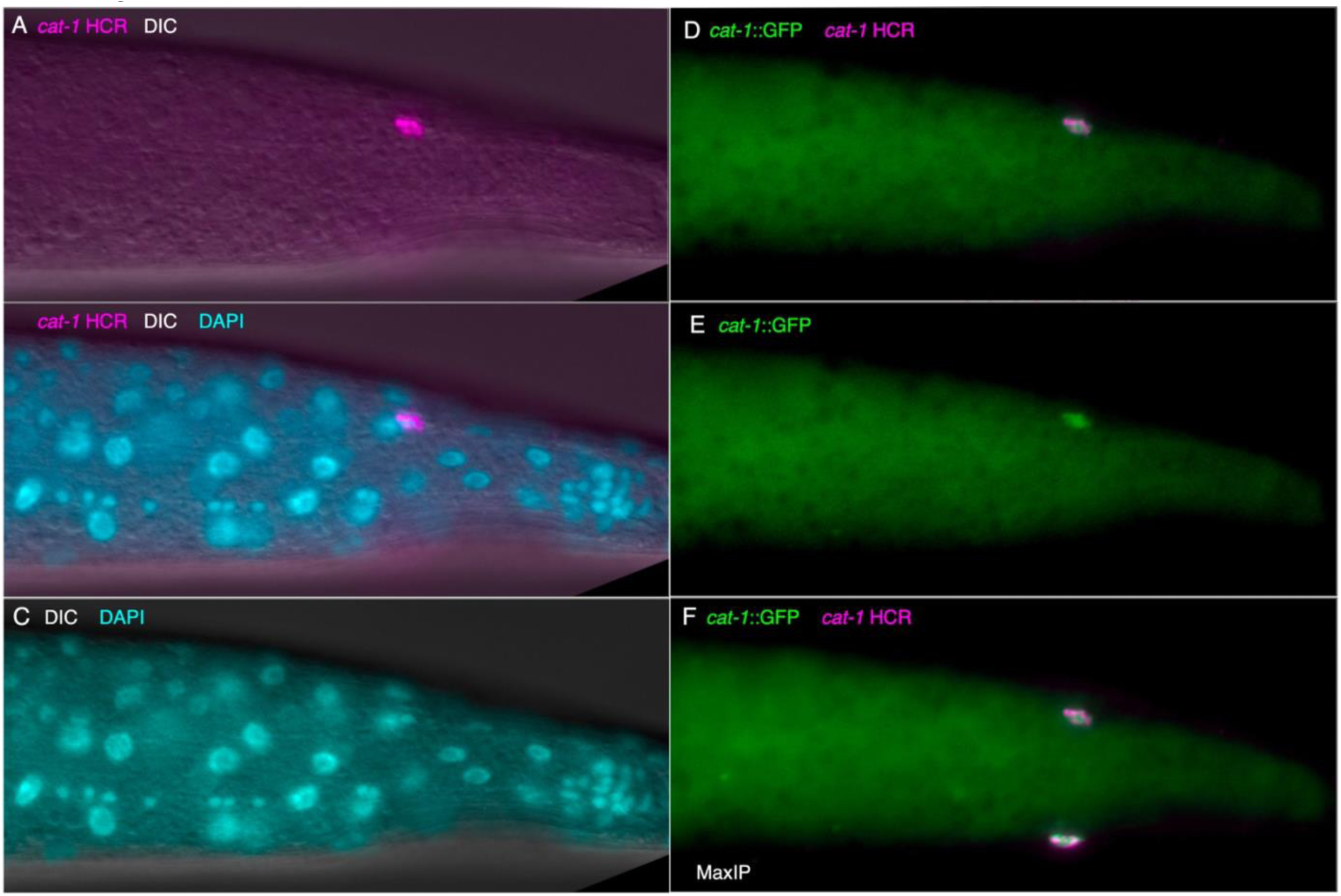
Colocalization of *cat-1 transcripts* and *cat-1*::GFP reporter in posterior body dopaminergic neurons (PDEs). Anterior is to the left in all panels. (A-E) Single focal plane. (A) *cat-1* HCR fluorescence (magenta) and DIC. (B) As in A, with DAPI. (C) DIC and DAPI, which shows a small, compact nucleus associated with the *cat-1* HCR signal. (D) *cat-1*::GFP (green) and *cat-1* transcripts. (E) *cat-1*::GFP alone (F) MaxIP of several focal planes to show colocalization in PDE neurons on both sides of the body wall.

**Suppl Fig S12.**
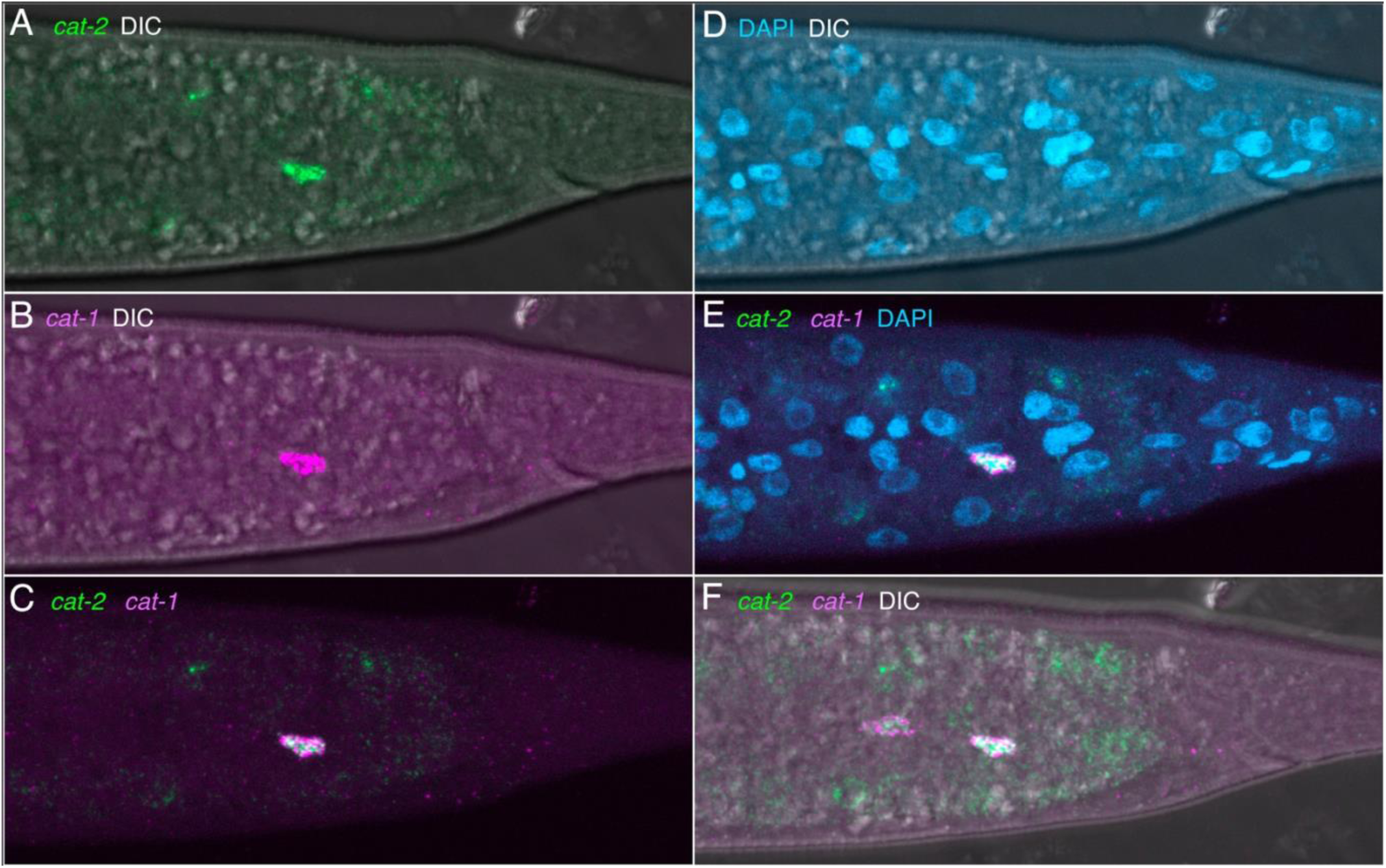
Co-expression of *cat-2* and *cat-1* transcripts in PDE neurons. Anterior is to the left in all panels. (A-E) Right side of posterior body, MaxIP of same set of focal planes. (A) *cat-2* HCR fluorescence (green) and DIC. (B) *cat-1* HCR fluorescence (magenta) and DIC. (C) Colocalization of *cat-2* and *cat-1* transcripts in PDE neuron. (D) DAPI and DIC (E) *cat-2* and *cat-1* transcripts are associated with compact PDER nucleus associated with the *cat-1* HCR signal. (F) MaxIP including left side focal planes to show colocalization in PDE neurons on both sides of the body.

**Suppl Fig S13.**
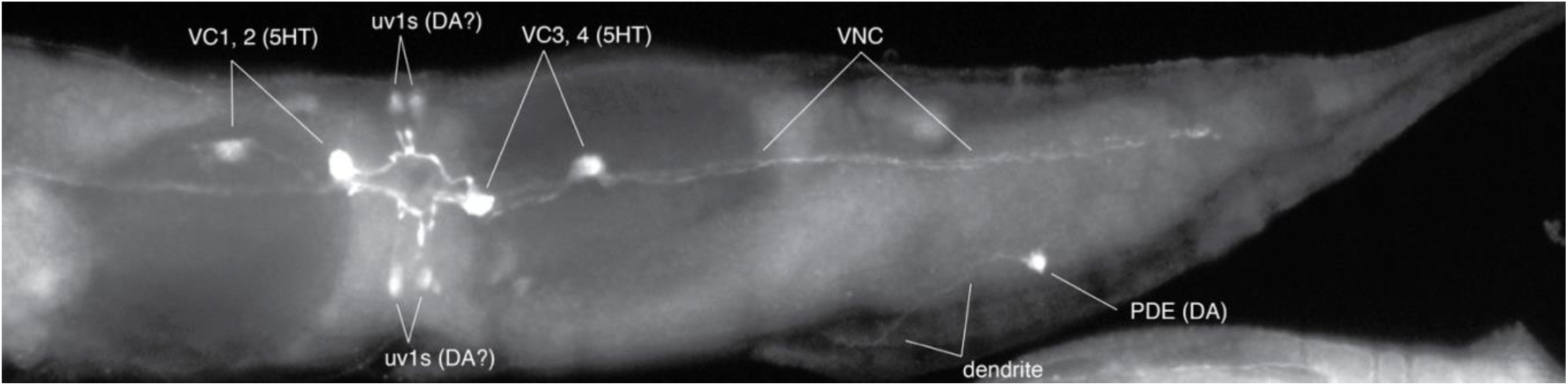
Monoaminergic cells in the central and posterior body of *P. pacificus*. Anterior is to the left. Serotonergic and presumptive dopaminergic cells and in the central and posterior body of adult hermaphrodite *P. pacificus* revealed by 5-HTP-induced serotonin immunoreactivity; animal incubated with 5-HTP and subsequently stained with anti-serotonin. Dopaminergic neurons take up 5-HTP and convert it to serotonin. In the ventral nerve cord, VC motor neurons stain for serotonin (and are also seen *without* 5-HTP treatment). On either side of the vulva in the lateral body wall are teardrop-shaped cells that are likely homologs of *C. elegans* uterine vulva uv1 cells. In the mid-posterior lateral body wall is the dopaminergic PDE mechanosensory neuron, which extends a long dendrite anteriorly and laterally (with a slight turn dorsally at the end, in contrast to that seen in other free-living nematodes in which a much shorter dendrite extends dorsally from the soma (Rivard et al., 2010). The PDE dendrite is seen more clearly in Fig. 6E-F, showing *cat-1p::gfp* expression.

**Suppl Fig S14.**
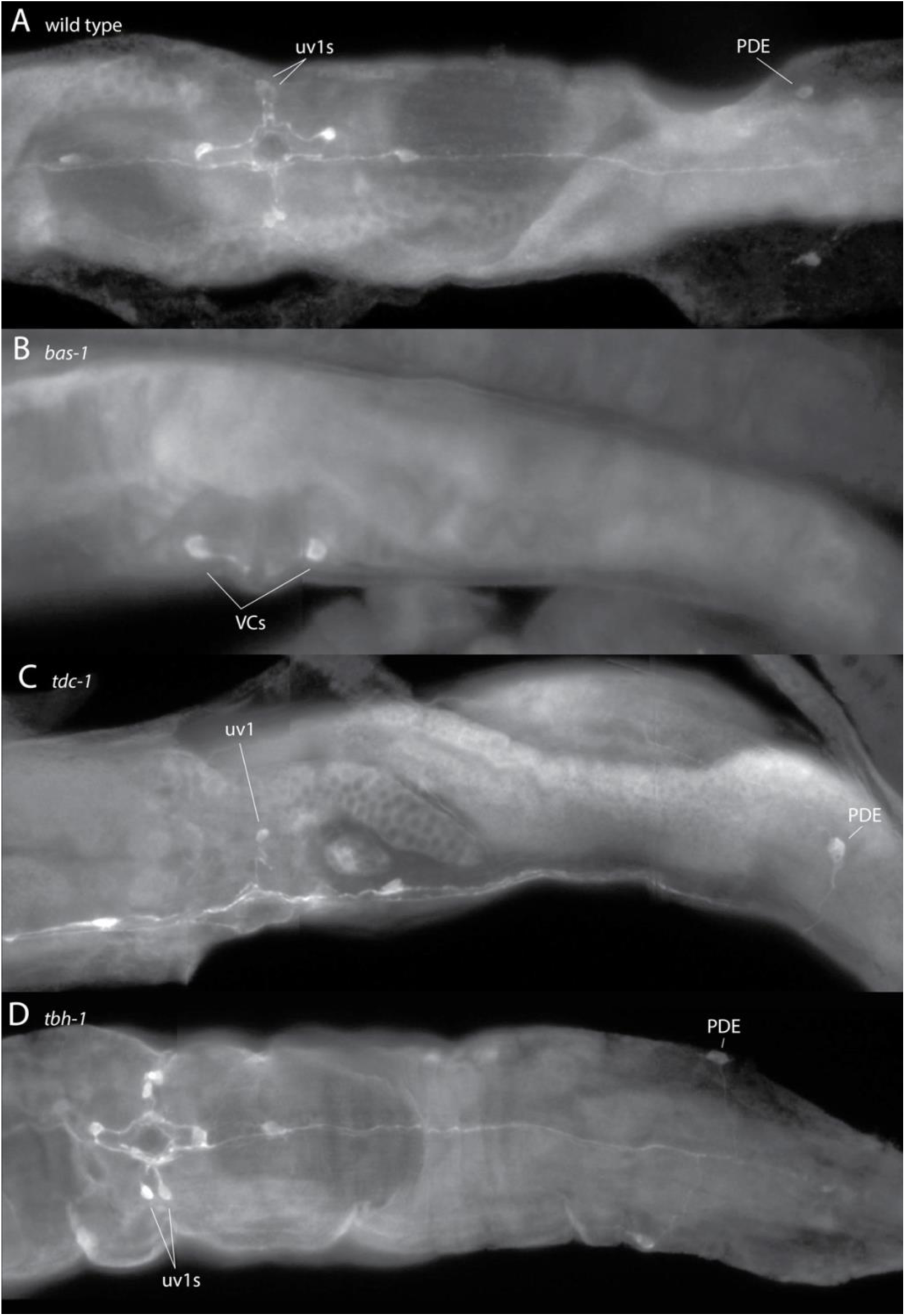
5HTP-induced serotonin immunoreactivity requires *bas-1* / AAADC function. Anterior is to the left, each panel shows anti-serotonin staining in adult hermaphrodite central and posterior body in worms exposed to 5mM 5-HTP in NGM plates for 24 hr. Such treatment typically results in high background staining in nearly all tissues. (A) Wild type (PS312) worms show both serotonin-IR uv1s and PDEs. Ventral view (uv1s and PDEs seen on both sides). (B) *bas-1(tu629)* mutant worms show no stained uv1s or PDEs, showing the requirement for AAADC function to decarboxylate 5HTP to 5HT. Interestingly, *bas-1* mutants show weak to moderately-stained VC neurons and sometimes NSM neurons in the head, independent of 5HTP exposure (not shown). Lateral view. (C) *tdc-1(tu1007)* worms show staining in uv1s and PDEs. Note that in some worms (unrelated to 5HTP exposure), as shown here, only a single uv1 cell is found on a given side. Lateral view. (D) *tbh-1(cbh32)* worms show staining in uv1s and PDEs. Ventral view (uv1s and PDEs seen on both sides).

**Suppl Fig 15.**
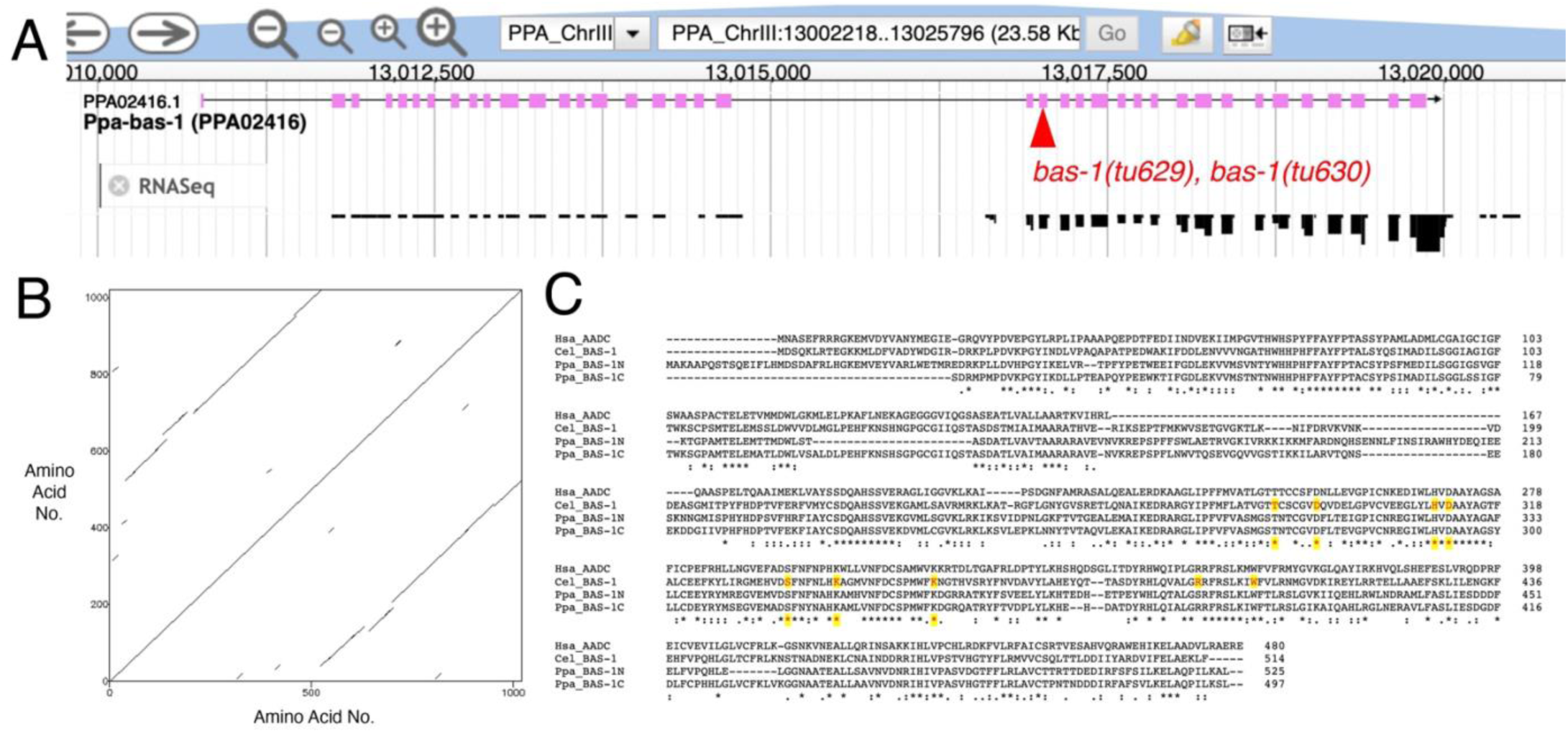
*Ppa bas-1* locus is a tandem duplication – *bas-1* mutants may have residual function. (A) *Ppa-bas-1* locus has a tandem duplication of BAS-1/AADC coding sequence (tracks from Wormbase genome browser, WS297). Gene model showing coding exons (pink boxes, upper track) and RNASeq coverage (lower track); existing *bas-1* mutations (causing premature stops) are in the second section of coding sequence (C-terminal duplicate) indicated by red triangle. Differences in RNASeq coverage suggest that the regions are expressed independently. (B) Dot plot of 1022 AA predicted protein (made with https://www.bioinformatics.nl/cgi-bin/emboss/dotmatcher) showing duplicated region starting around 500 AAs. (C) Multiple sequence alignment of human AADC, *C. elegans* BAS-1, *Ppa* N-terminal BAS-1 and C-terminal BAS-1 proteins. Examples of known critical function AAs are highlighted (red letters with yellow background); all but one are conserved in the N-terminal BAS-1 which appears likely to be functional, either by itself or as a truncated internally duplicated protein.

**Suppl Fig S16.**
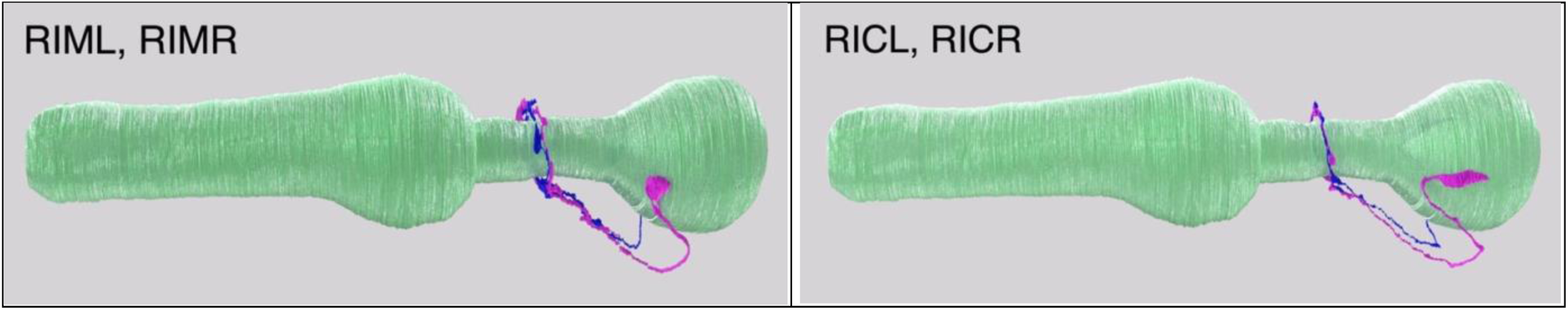
*P. pacificus* RIM and RIC neuron 3D renderings. Putative tyraminergic (RIM) and octopaminergic (RIC) neurons, lateral views. Anterior to the left. Pharynx in light green, left neuron in magenta, right side blue (Cook et al., 2025).

**Suppl Fig S17.**
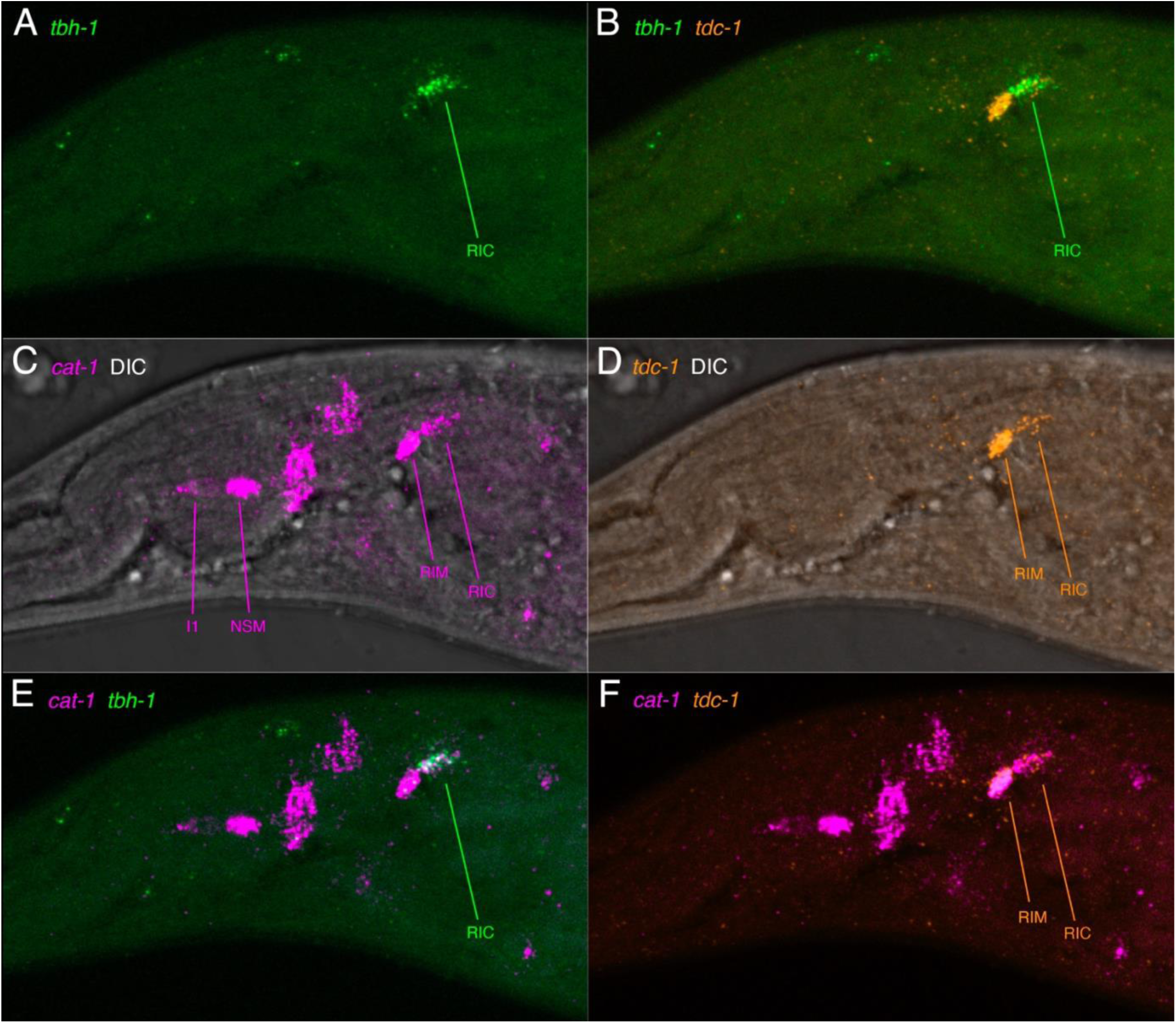
*Colocalization of cat-1* transcripts with *tdc-1* and *tbh-1* transcripts in RIM and RIC neurons. Anterior left, ventral down, ventro-lateral view. All images of the same adult hermaphrodite head, MaxIP of a few focal planes showing one side. (A) *tbh-1* transcripts (green) expressed in RIC. (B) *tdc-1* transcripts (orange), showing two cells in lateral ganglion, RIM and RIC. (C) *cat-1* transcripts (magenta) with DIC to show location of the pharynx as a landmark. Some *cat-1*-positive cells are identified. (D) *tdc-1* with DIC, expression in RIM and RIC. (E) Colocalization of *cat-1* with *tbh-1* transcripts in RIC. (F) Colocalization of *cat-1* with *tdc-1* transcripts in RIM and RIC.

**Suppl Fig S18.**
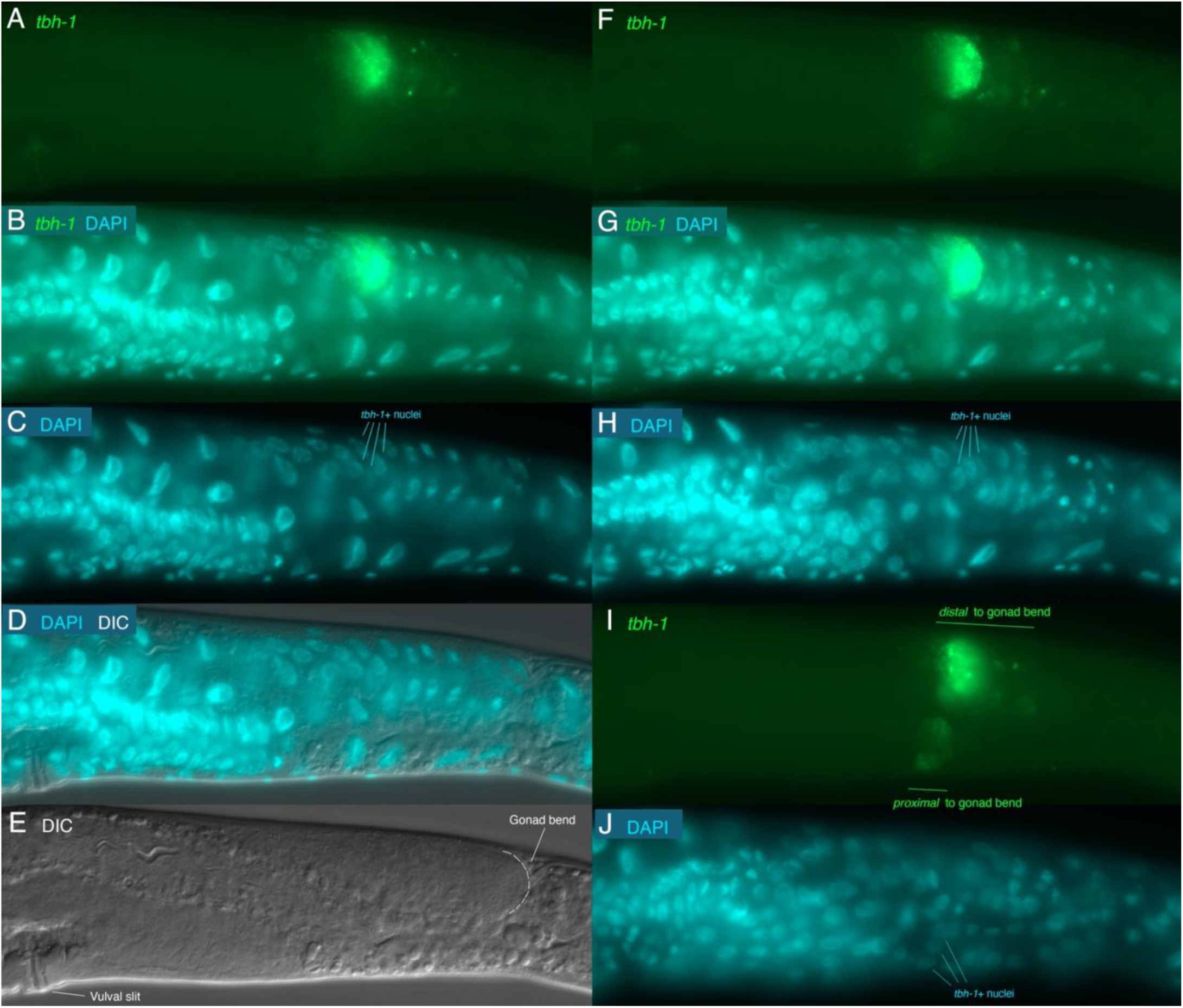
Expression of *tbh-1* transcripts in the gonad. Anterior to the left, ventral down, showing posterior body. (A-H) distal (and dorsal) gonad staining; (A-E) same single focal plane; (F-H) different focal plane. (I, J) same single focal plane, showing proximal (ventral) gonad. ‘Proximal’ indicates closer to the vulva, distal is further away within the gonad. Regions identified based on (Rudel et al. 2005). (A, F) HCR of *tbh-1* transcripts (green) shows expression in likely gonadal sheath cells. (B, G) Both *tbh-1* and DAPI signals. (C, H) DAPI staining with nuclei strongly *tbh-1*-expressing indicated. Some adjacent nuclei (in front and behind) appear to express *tbh-1* at a lower level. (D) DAPI and DIC. (E) DIC showing that the region of *tbh-1* expression is distal to the gonad bend where the gonad shifts from dorsal to ventral. (I) *tbh-1* transcripts are expressed at a lower level in a second region after the gonad bend (proximal to the vulva). (J) DAPI staining with some *tbh-1*-expressing nuclei indicated.

**Suppl Fig S19.**
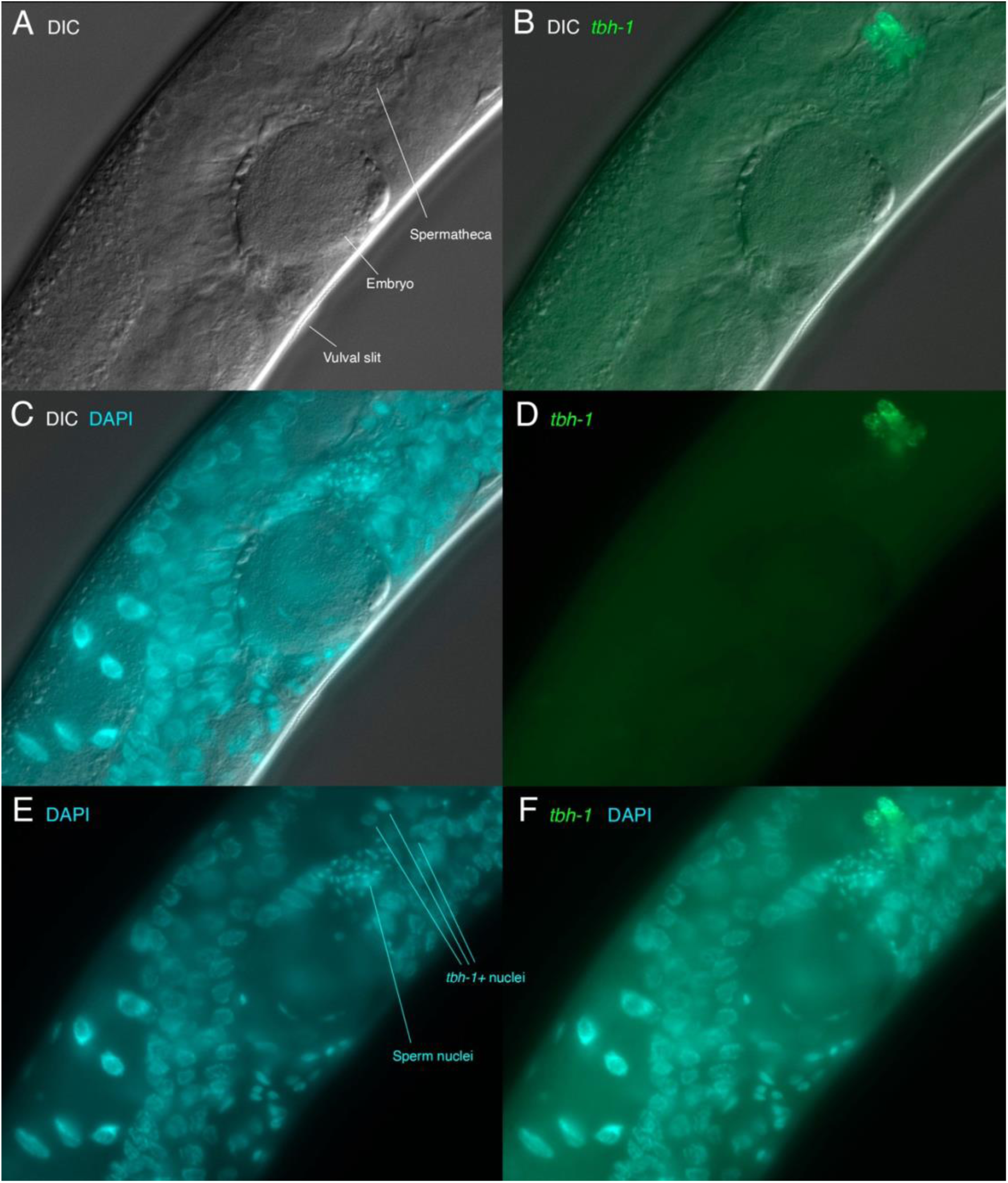
Expression of *tbh-1* transcripts in the spermatheca (proximal gonad). Adult hermaphrodite with embryos, vulval region, dorsal to the upper left, ventral is lower right. Same focal plane for each panel. (A) DIC showing locations as indicated. A single embryo is present in the uterus between the vulval opening and more distal spermatheca. (B, D, F) Expression of *tbh-1* transcripts by HCR. (B) *tbh-1* transcripts (green) + DIC. (C) DIC and DAPI (cyan) to show nuclei. (D) *tbh-1* alone. (E) DAPI with sperm nuclei within spermatheca indicated. Just distally is gonad constriction with *tbh-1*-expressing nuclei indicated. (F) *tbh-1* transcripts + DAPI staining.

**Suppl Fig S20.**
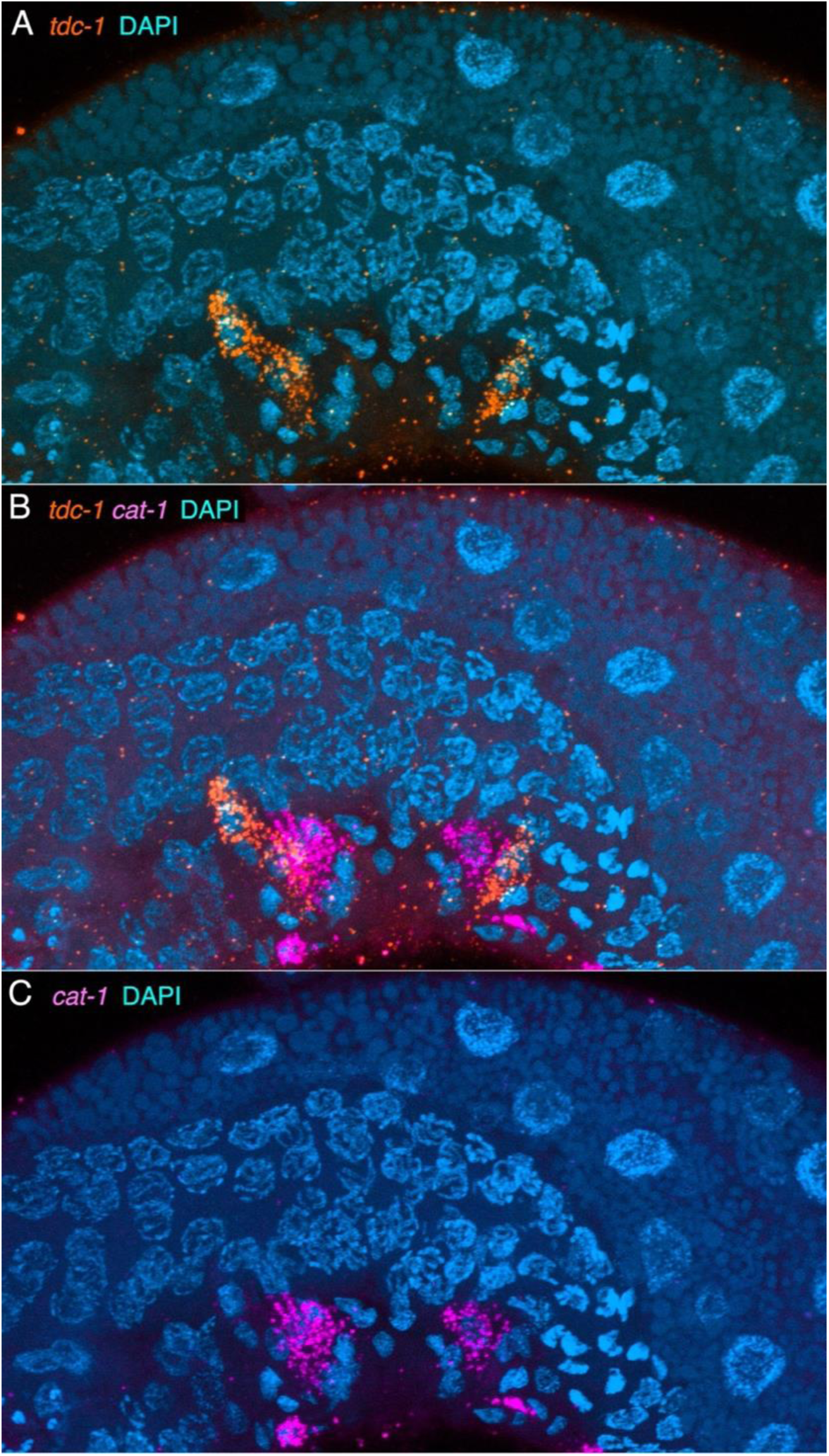
Expression of *tdc-1* and *cat-1* transcripts is in adjacent vulval region cells. Adult hermaphrodite vulval region, anterior to the right and ventral down, MaxIP of 21 z-planes on the left side to approximately the midline. (A) *tdc-1* transcripts (orange) associated with two nuclei (DAPI, cyan) per quandrant. (B) Co-labeling shows that *cat-1* transcripts (magenta) are associated with two nuclei closer to the vulval pore in each quadrant. Although this image suggests possible overlap in expression in some cells (left / anterior), most preparations show clear separation of expression in adjacent cells, like that seen here in the posterior cells. (C) *cat-1* transcripts expressed close to the vulval pore, lateral to the ventral nerve cord. Some smaller VC neurons expressing *cat-1* are seen at the bottom of the image.

**Suppl Fig S21.**
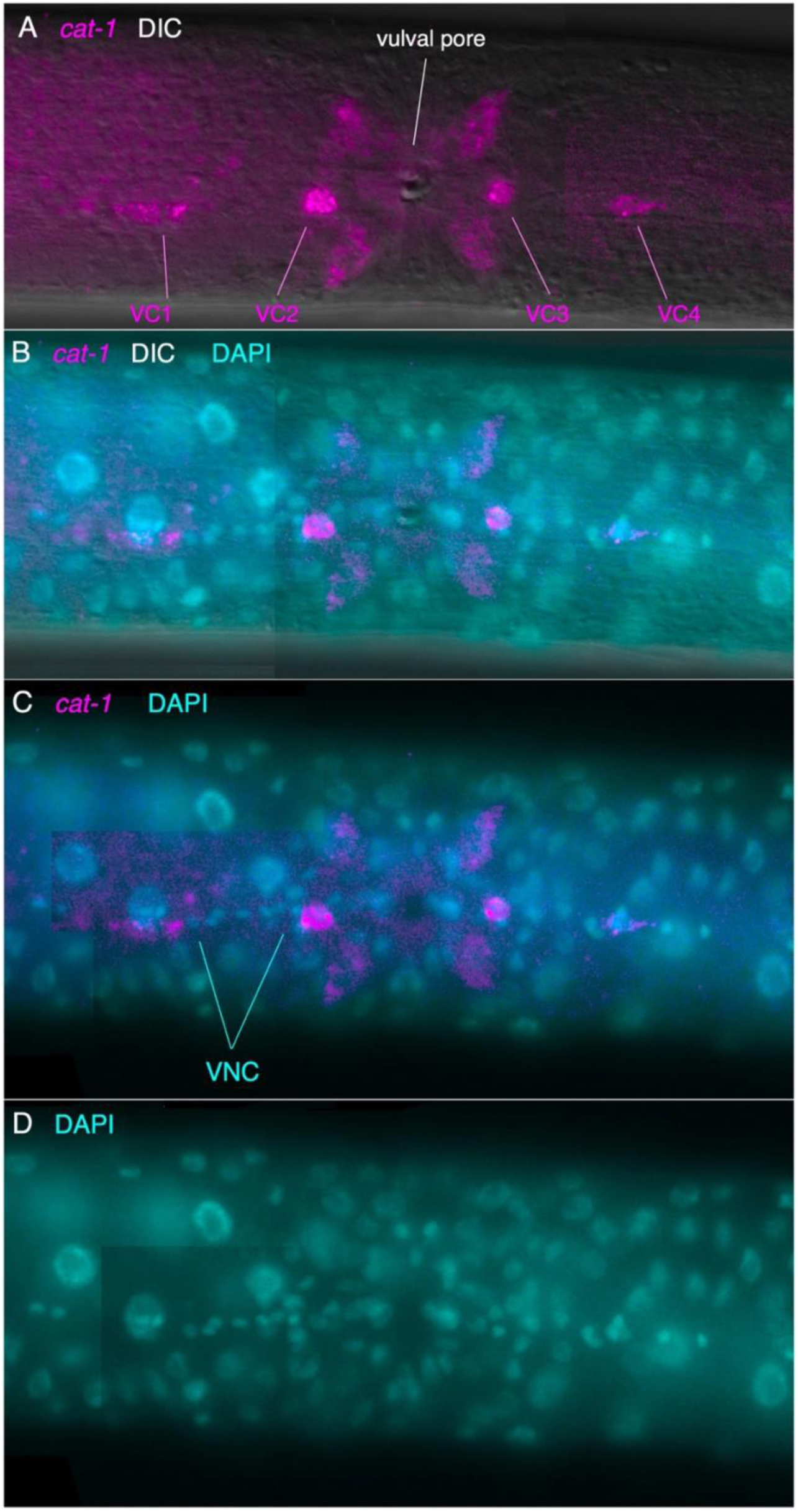
Expression of *cat-1* transcripts in the midbody VC neurons and vulval cells. Ventral view; anterior is to the left in all images. Each panel is the same single focal plane (panels B-D, however, a partly montages that include a small region from a nearby focal plane to better show the ventral nerve cord). (A-C) HCR for *cat-1* transcripts (magenta) with DIC and/or DAPI (cyan). (D) DAPI alone to better show vulval region and VNC.

**Suppl Fig S22.**
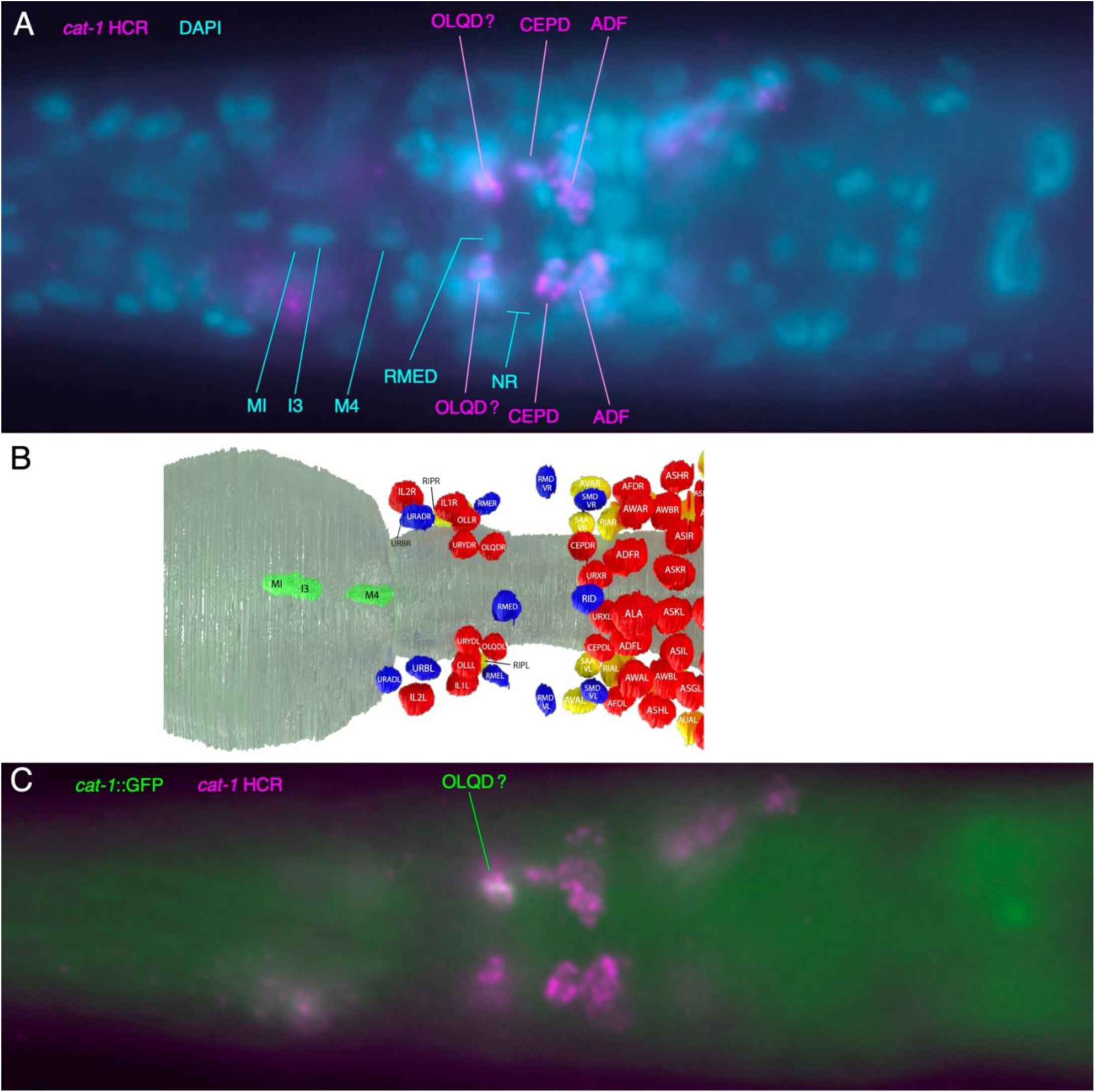
Other *cat-1*-expressing cells in the head. Expression of *cat-1* transcripts via HCR in dorsal head in *cat-1*::GFP strain. Anterior to the left. (A, C) same dorsal focal plane. (A) *cat-1* HCR fluorescence and DAPI, particularly showing likely identification of OLQDs as dorsal anterior ganglion nuclei just anterior to the nucleus-free region of the nerve ring (NR). Readily identifiable DAPI stained nuclei in the dorsal anterior pharyngeal bulb are indicated. (B) Dorsal view map of dorsal neuronal nuclei around NR, including dorsal pharyngeal neurons (green) identified in (A) from EM reconstruction (Cook et al., 2025); pharynx, partly transparent, in darker green. Other head nuclei (non-pharyngeal) colored by apparent function: sensory (red), motor (blue), interneuron (yellow). (C) *cat-1*::GFP transgene expression and *cat-1* HCR fluorescence colocalization in one putative OLDQ neuron. Same focal plane as (A).

**Suppl Fig S23.**
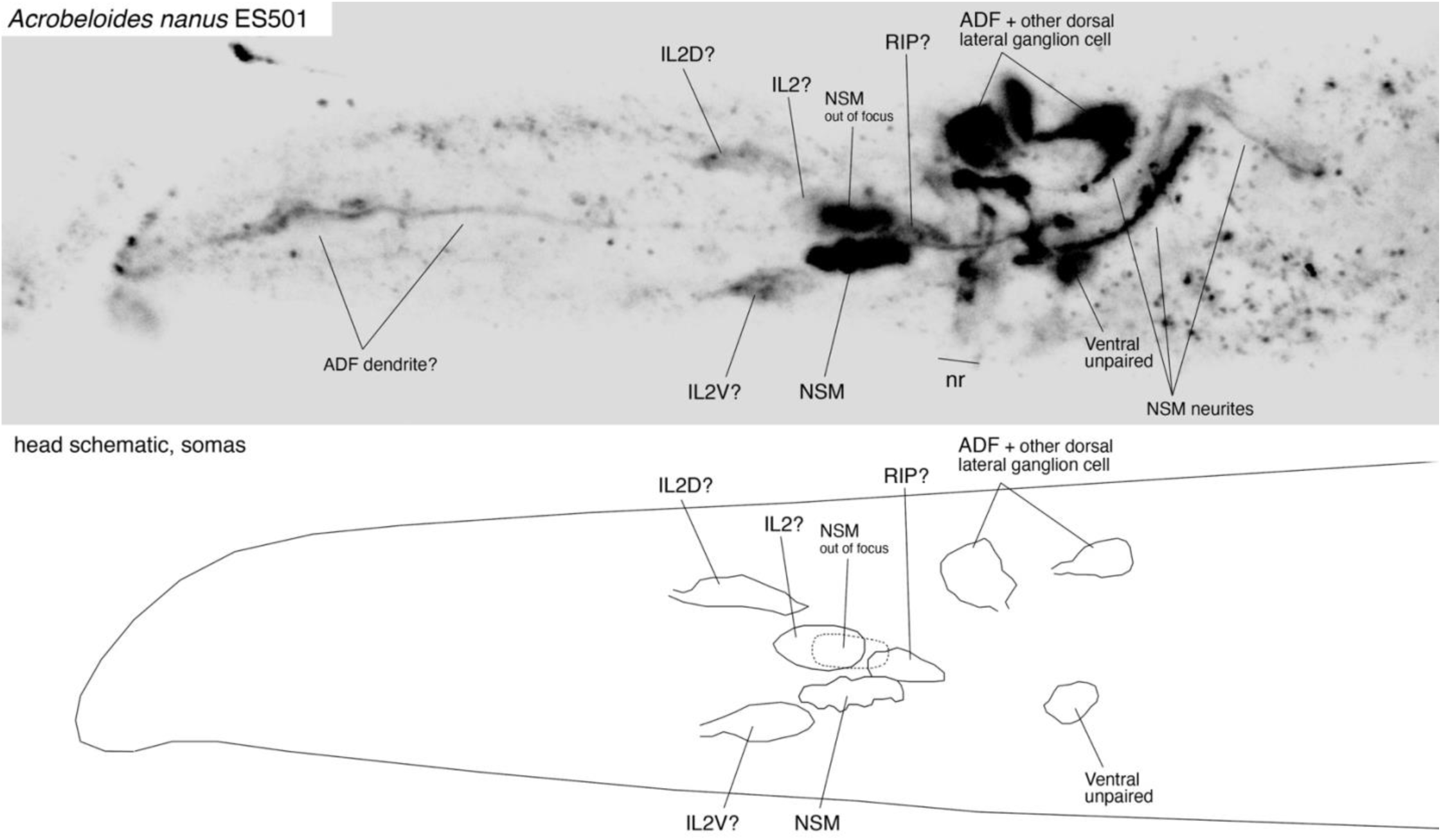
*Acrobeloides nanus* ES501 head serotonin-IR neurons. Anterior left, ventral down, lateral view; confocal MaxIP of a few focal planes on left side (top); likely cell and neurite identifications as indicated. Brighter cells and neurites are over-exposed to show less brightly stained somas. nr – location of nerve ring. Schematic showing outlines of somas matching image (below).

**Suppl Fig S24.**
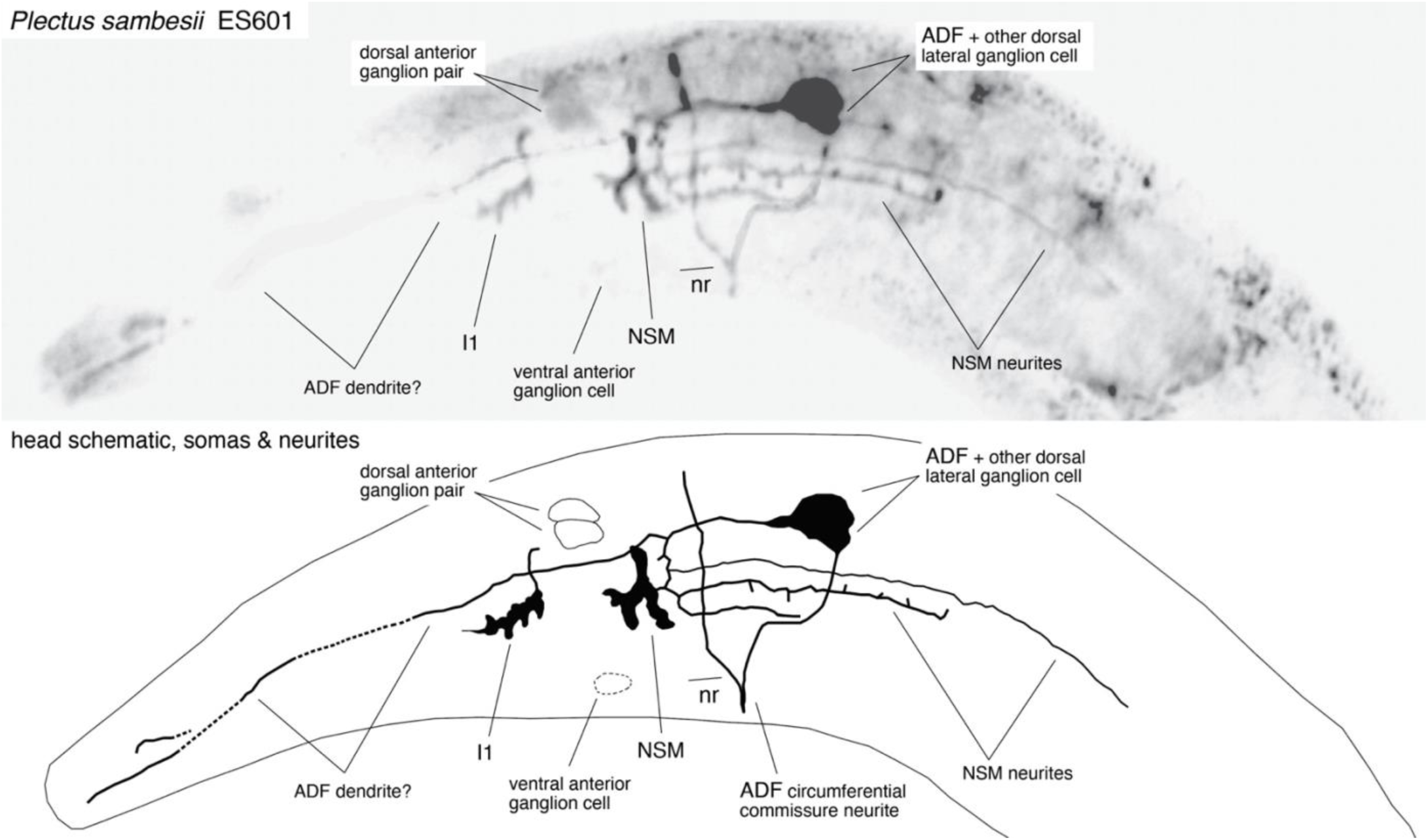
*Plectus sambesii* ES601 head serotonin-IR neurons. Anterior left, ventral down, lateral view; montage of confocal focal planes on left side (top); likely cell and neurite identifications as indicated. nr – location of nerve ring. Schematic showing outlines of somas and neurites matching image (below).

**Suppl Fig S25.**
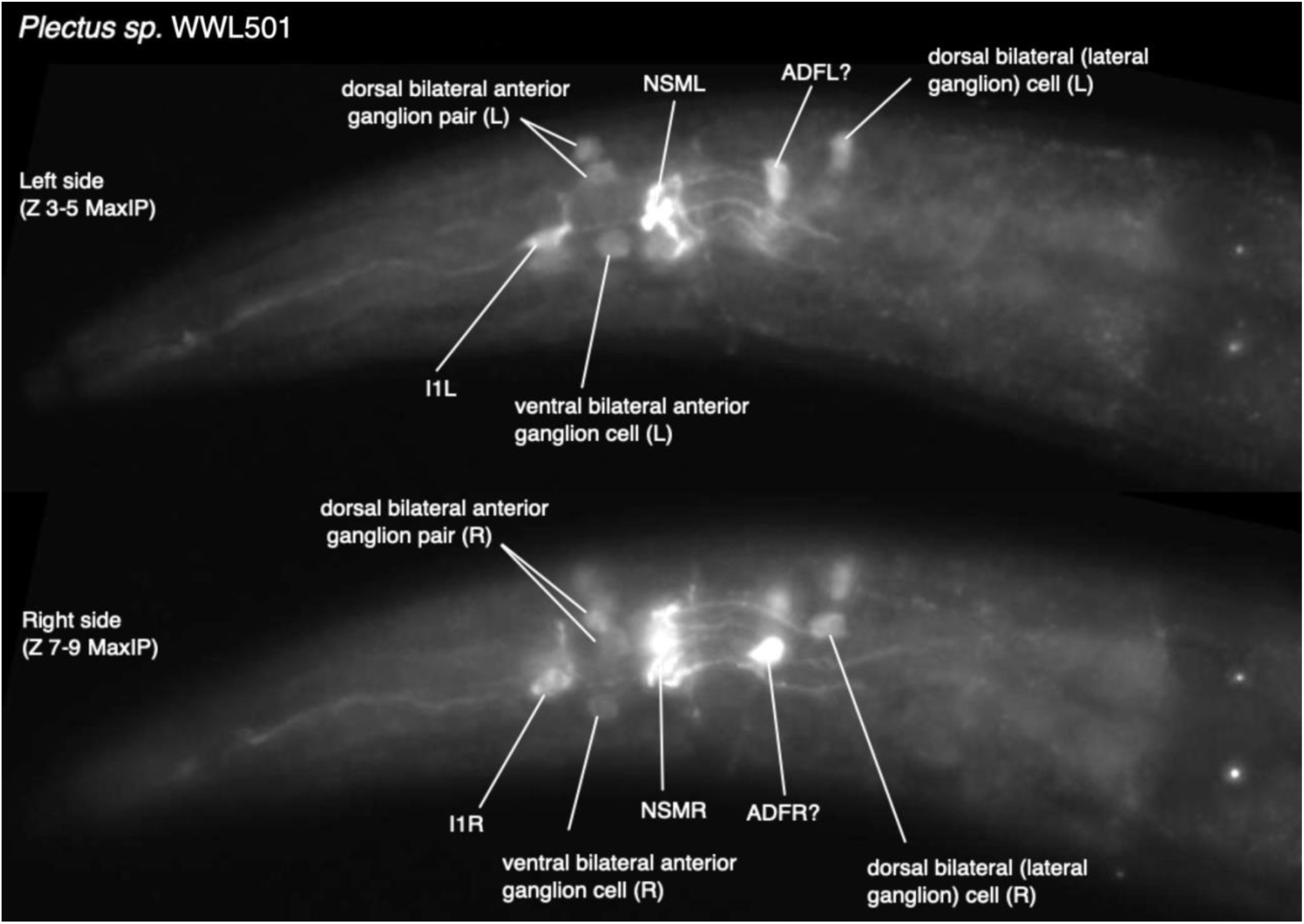
*Plectus sp.* WWL501 head serotonin-IR neurons. Anterior left, ventral down, lateral views; confocal MaxIPs of a few focal planes on left (top) and right (bottom) sides. Likely cell identifications as indicated.

**Suppl Fig S26A.**
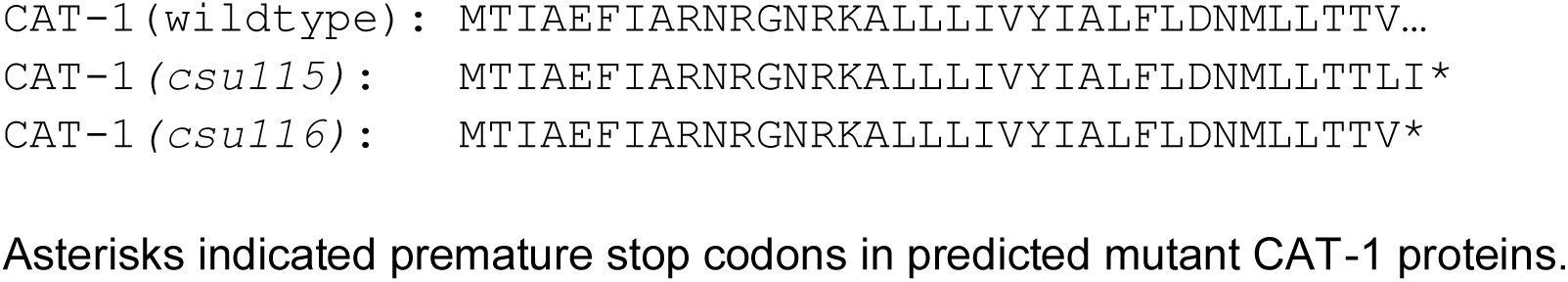
*P. pacificus* CAT-1 mutant alignments.

**Suppl Fig S26B.**
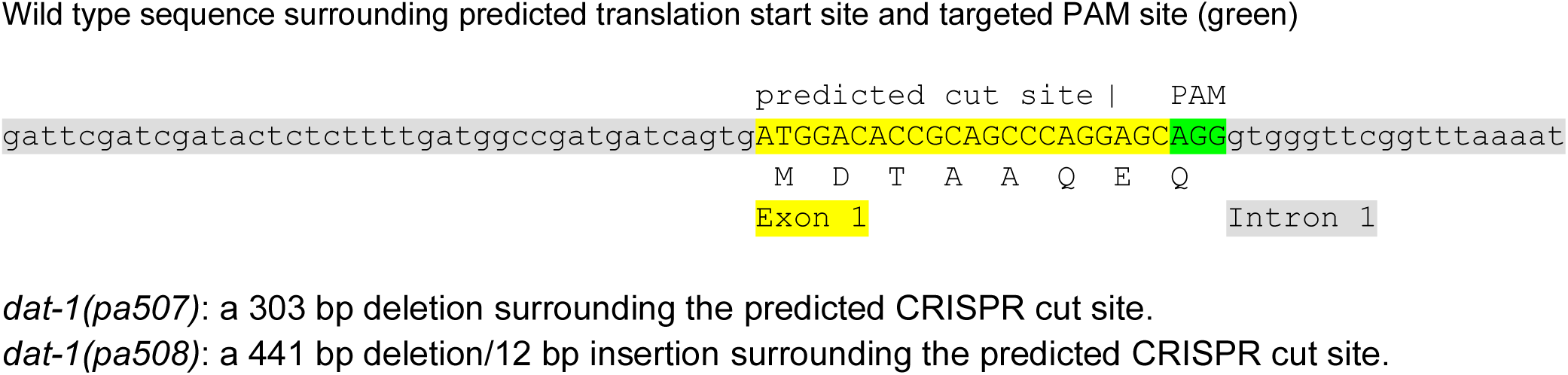
*P. pacificus* DAT-1 mutants (N-terminal deletions)

**Suppl Fig 27.**
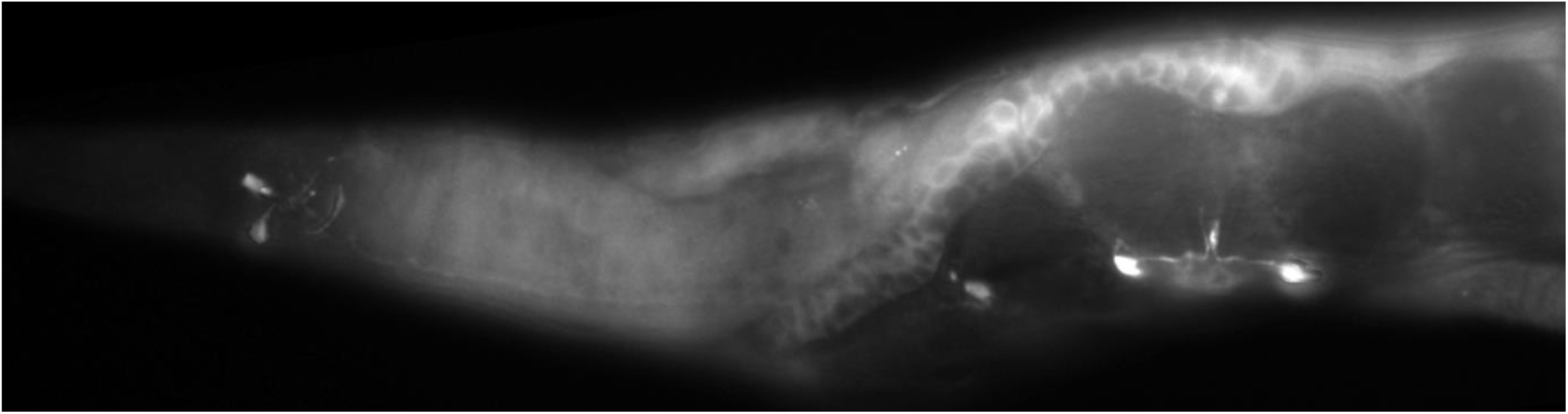
Anti-5HT staining in *P. pacificus cat-1* mutants is variably reduced. Anterior to the left, ventro-lateral view, montage. Head and midbody VNC region of *cat-1(csu116)* adult hermaphrodite. In this individual, NSM neurons in the head are moderately stained; other serotonin-IR neurons are not seen. VC neurons in VNC stain similarly to wildtype (VC1-3 shown; VC4, which also stains moderately, is out of the plane of focus). Although both *cat-1* mutations likely cause a complete loss of function, we observed a variable reduction in serotonin-IR, as has been seen previously in *C. elegans cat-1* mutants (Desai et al., 1988) (Loer and Kenyon 1993). Remaining serotonin-IR is not surprising since neurons should continue to synthesize serotonin via *tph-1/*TPH and *bas-1/*AADC expression (Figure 1), even if the cells are unable to package the neurotransmitter into synaptic vesicles. In some preparations, we saw loss of serotonin-IR in most or all head neurons and midbody VC neurons; in other preparations (like that above), we observed reduced but considerable staining in all known serotonin-IR cells. The behavioral phenotypes we observed in *P. pacificus cat-1* mutants are consistent with a loss of VMAT function in spite of remaining neurotransmitter within the neurons.

**Suppl Fig S28.**
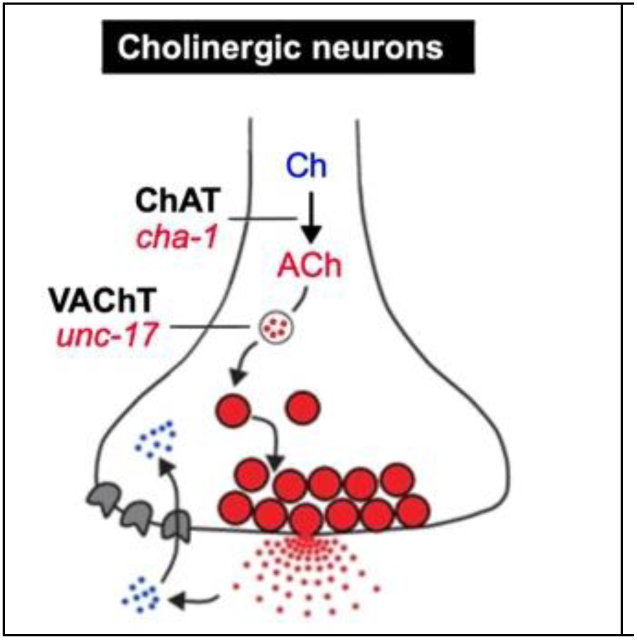
Cholinergic proteins and nematode genes. Proteins encoded by identified nematode genes are indicated in black, bold, capital letters (abbreviations: Ch - choline; ACh - acetylcholine; ChAT - choline acetyltransferase, required for acetylcholine synthesis; VAChT - vesicular acetylcholine transporter, required for packaging acetylcholine into synaptic vesicles); nematode gene names (originally from *C. elegans*) are italicized below the protein name abbreviations.

**Suppl Fig S29.**
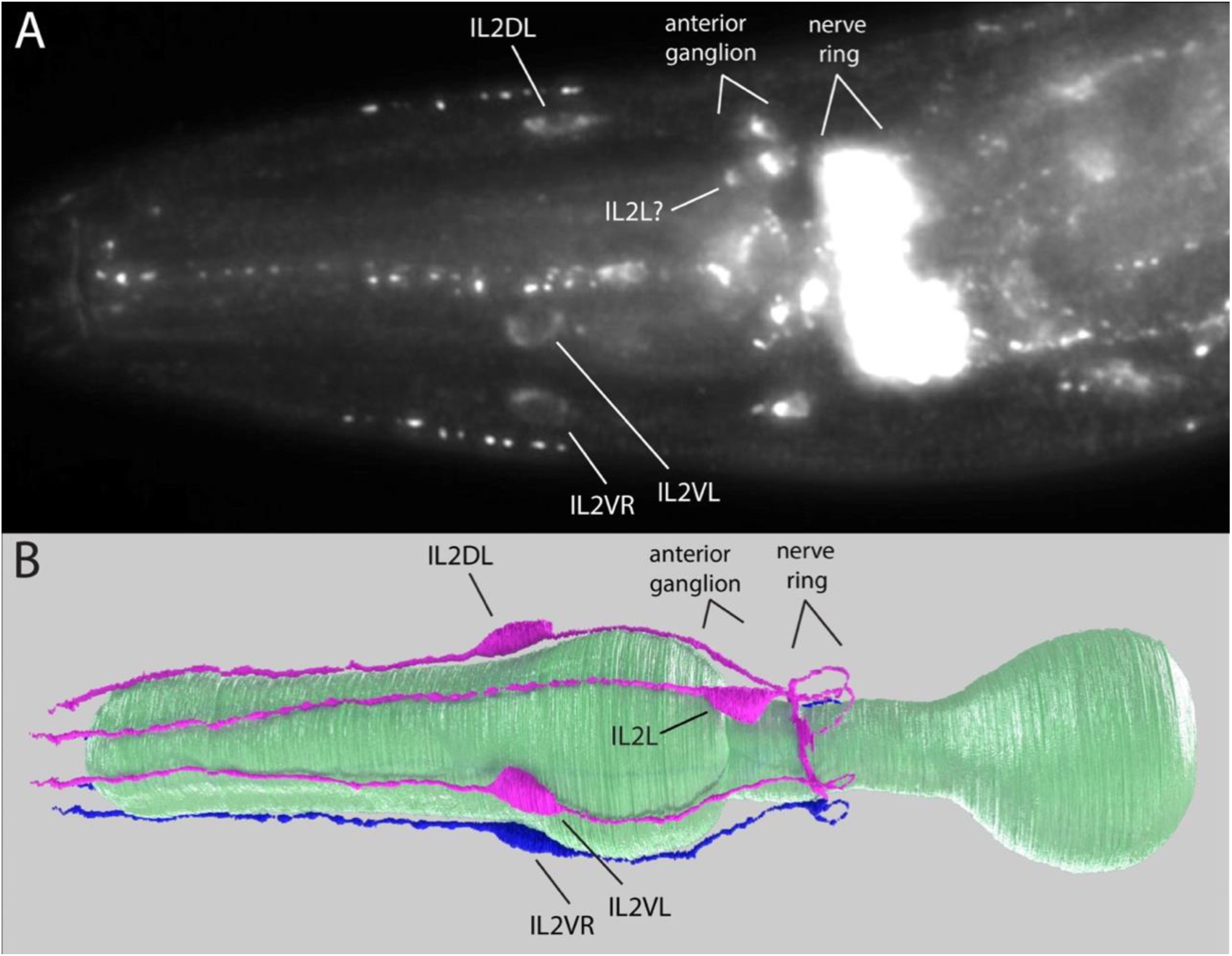
Cholinergic neurons and nerves in the head of *P. pacificus*. Both images are left side views, anterior to the left. (A) Wildtype adult hermaphrodite head staining with a ‘cholinergic mix’ of monoclonal antibodies to *C. elegans* cholinergic proteins CHA-1 (ChAT) and UNC-17 (VAChT) shows extensive staining in the nervous system. The nerve ring is very strongly stained; head ganglia surrounding the nerve ring, including the anterior ganglion, are moderately stained. Although it is difficult to identify individual neuronal somas, the somas of IL2D and IL2V neurons are well isolated from all other neurons, being just in front of the pharyngeal anterior bulb, so can be clearly identified. In this view, the IL2DL (left) is seen, and both IL2Vs (L & R). Another soma in the anterior ganglion could be IL2L. (B) 3D rendering of IL2 neurons, left lateral view (and slightly ventral), anterior to the left. IL2Ds and IL2Vs are further anterior; IL2 L & R are within the anterior ganglion near to the nerve ring. Left side IL2s are rendered in magenta; right side IL2s in blue. The pharynx is in green.

**Suppl Fig S30.**
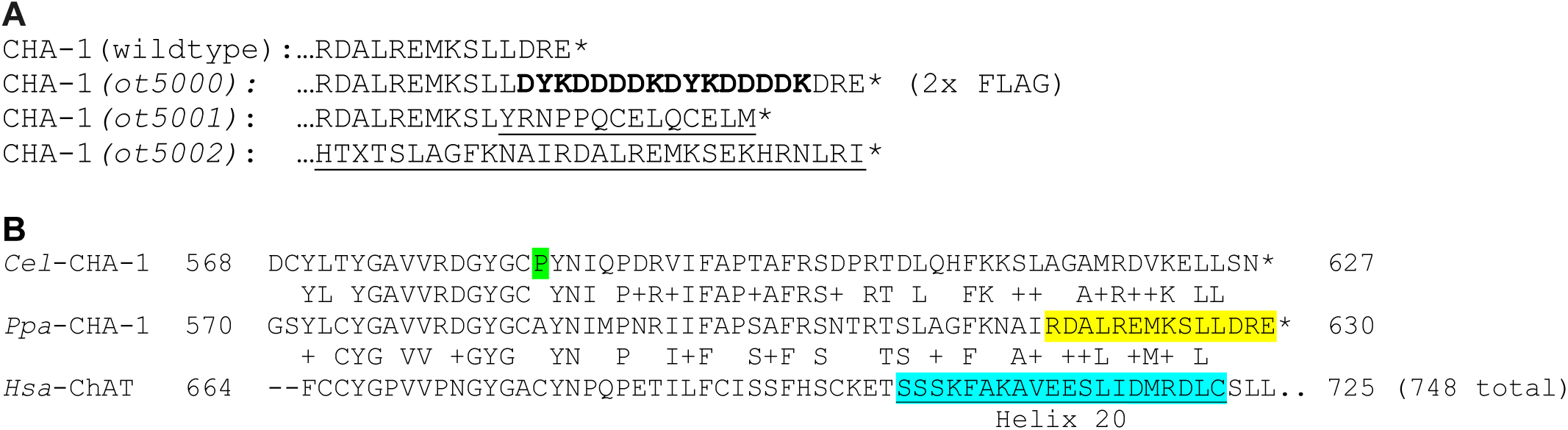
CHA-1 C-terminus alignments of mutant alleles and wildtype proteins. (A) Alignments of wildtype *P. pacificus* CHA-1 protein showing alterations in mutant alleles. * denotes stop codon; **bolded** region shows in-frame 2x FLAG epitope tag insertion (DYKDDDDK); <u>underlined</u> regions indicate results of frame-shift from indel mutations. Alignments of C-termini from nematode and mammalian proteins. There is considerable conservation of the C-terminus. Between each sequence shows identical and similar (+) AAs. *Cel – C. elegans, Ppa – P. pacificus, Hsa – Homo sapiens. Cel* sequence highlight: P584L missense mutation results in 99% loss of ChAT activity (Rand and Russel, 1984). *Ppa* sequence highlight: *Ppa* CHA-1 wildtype C-terminus portion shown in *Ppa* mutant alignments (A). *<u>Hsa</u>*<u>-ChAT highlight</u>: region of Helix 20 in rat ChAT crystal structure (Govindasamy et al., 2004).

**Suppl Fig S31.**
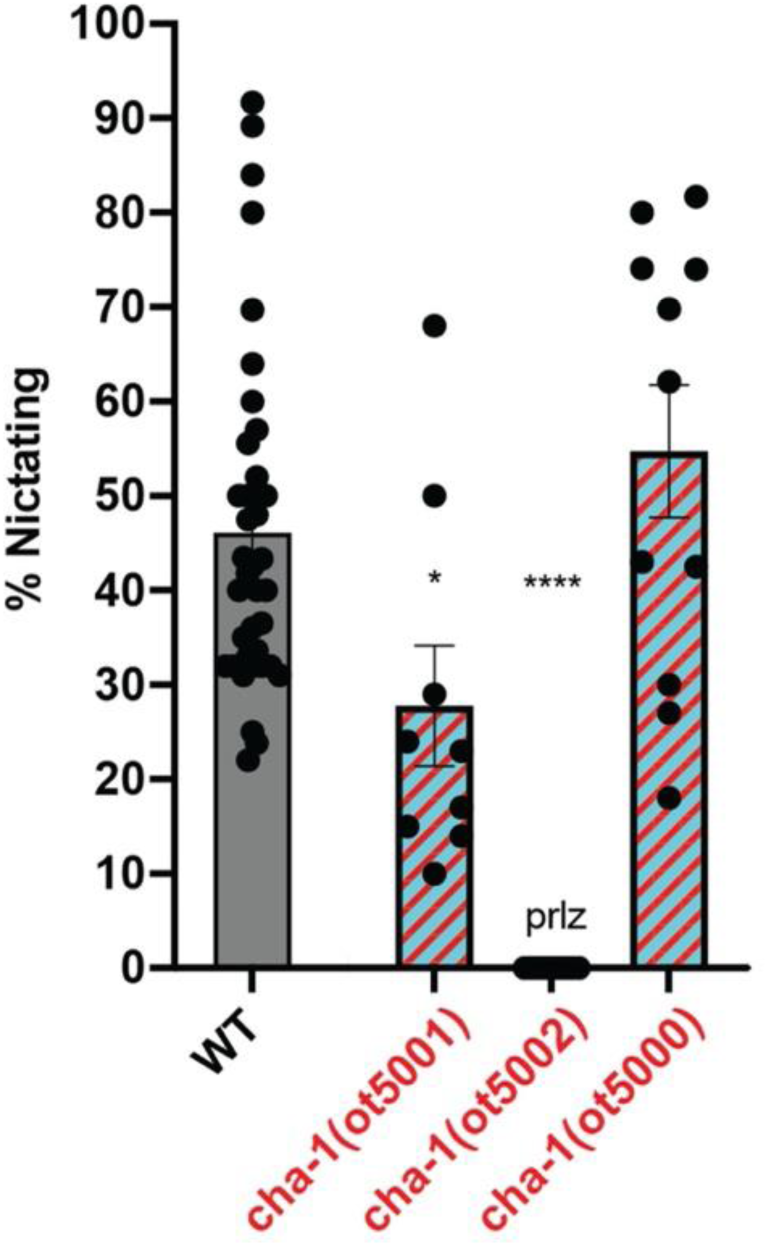
Effect of cholinergic function loss on nictation is inconclusive. Reduction-of-function alleles in the acetylcholine biosynthesis gene *cha-1*, including the C-terminal 2x-FLAG strain (*ot5000*), and two C-terminal indel mutants were found in the same CRISPR knock-in screen (*ot5001, ot5002*) (Suppl Figure 19) have temperature-sensitive phenotypes.^†^ In these nictation assays, the alleles showed a partial nictation phenotype (*ot5001*), paralysis of locomotion (*ot5002*), or no effect (*ot5000,* in frame FLAG insertion). Between 30-60 animals participate in each nictation assay and at least 6 assays were performed for each genotype. *P<0.05, ***P<0.001, ****P<0.0001 Dunnett’s multiple comparisons test show significant difference to wildtype (WT). ^†^ All three *cha-1* strains were found to be temperature-sensitive; this may reflect a temperature-sensitive process in CHA-1 protein function such has been observed in all *C. elegans cha-1* mutants (Duerr et al., 2021). Whereas the mutants appeared to move similarly to wildtype when raised at 20°C, mutants raised continuously at 25°C or higher became extremely uncoordinated or paralyzed, and progressively become sterile, particularly at temperatures above 25°C. Therefore, to test *cha-1* dauers, we let cultures starve at the permissive 20°C, then shifted to 25°C after food was depleted but before dauer formation, so that dauers were formed at the higher temperature. Given the pleiotropic phenotype of the *cha-1* alleles at 25°C, however, including defects in locomotion, whether the phenotype is truly specific to nictation is inconclusive. Furthermore, the nictation phenotype could also result simply from a slower response to the sand substrate beyond the 30-minute incubation time of the assay.

## Notes

### Competing Interest Statement

The authors have declared no competing interest.

